# KDS2010, a newly developed reversible MAO-B inhibitor, as an effective therapeutic candidate for Parkinson’s disease

**DOI:** 10.1101/2020.07.06.190579

**Authors:** Min-Ho Nam, Jong-Hyun Park, Hyo Jung Song, Ji Won Choi, Siwon Kim, Bo Ko Jang, Hyung Ho Yoon, Jun Young Heo, Doo-Wan Cho, Young-Su Yang, Su-Cheol Han, Sangwook Kim, Soo-Jin Oh, Sang Ryong Jeon, Ki Duk Park, C. Justin Lee

## Abstract

**Background and Purpose:** Monoamine oxidase-B (MAO-B) is a long-standing therapeutic target for Parkinson’s disease (PD), however, previous clinical studies demonstrated discouraging effects of currently available irreversible MAO-B inhibitors. Since KDS2010, a novel, potent, selective, and reversible MAO-B inhibitor, has been developed, here we tested its therapeutic potential in animal models of PD.

**Experimental Approach:** We designed and synthesized α-aminoamide derivatives and compared the specificity to MAO-B and reversibility of each compound with KDS2010. To investigate the in vivo therapeutic effect, we used MPTP mouse model with two different regimes of 3-day administration (pre-treatment or post-treatment) and 30-day administration. We assessed the therapeutic potential using behavioral and immunohistochemical analyses. Additionally, the functional recovery by KDS2010 was tested in 6-hydroxydopamine-induced and A53T-alpha-synuclein overexpression models. Lastly, to validate the potential as a clinical drug candidate, we investigated the pharmacokinetics and toxicity of KDS2010 in non-human primates.

**Key Results:** KDS2010 showed the highest potency, specificity, and reversibility among the α-aminoamide derivatives, with high bioavailability (>100%) and BBB permeability. KDS2010 also showed significant neuroprotective and anti-neuroinflammatory effects in the nigrostriatal pathway, leading to an alleviation of MPTP-induced parkinsonism in all administration regimes. In particular, the therapeutic effect of KDS2010 was superior to selegiline, an irreversible MAO-B inhibitor. KDS2010 also showed a potent therapeutic effect in 6-hydroxydopamine and A53T models. Moreover, KDS2010 showed virtually no toxicity or side-effect in non-human primates.

**Conclusion and Implications:** KDS2010 shows excellent therapeutic potential and safety in various PD animal models. KDS2010, therefore, could be a next-generation therapeutic candidate for PD.

**Representative Schematic:** 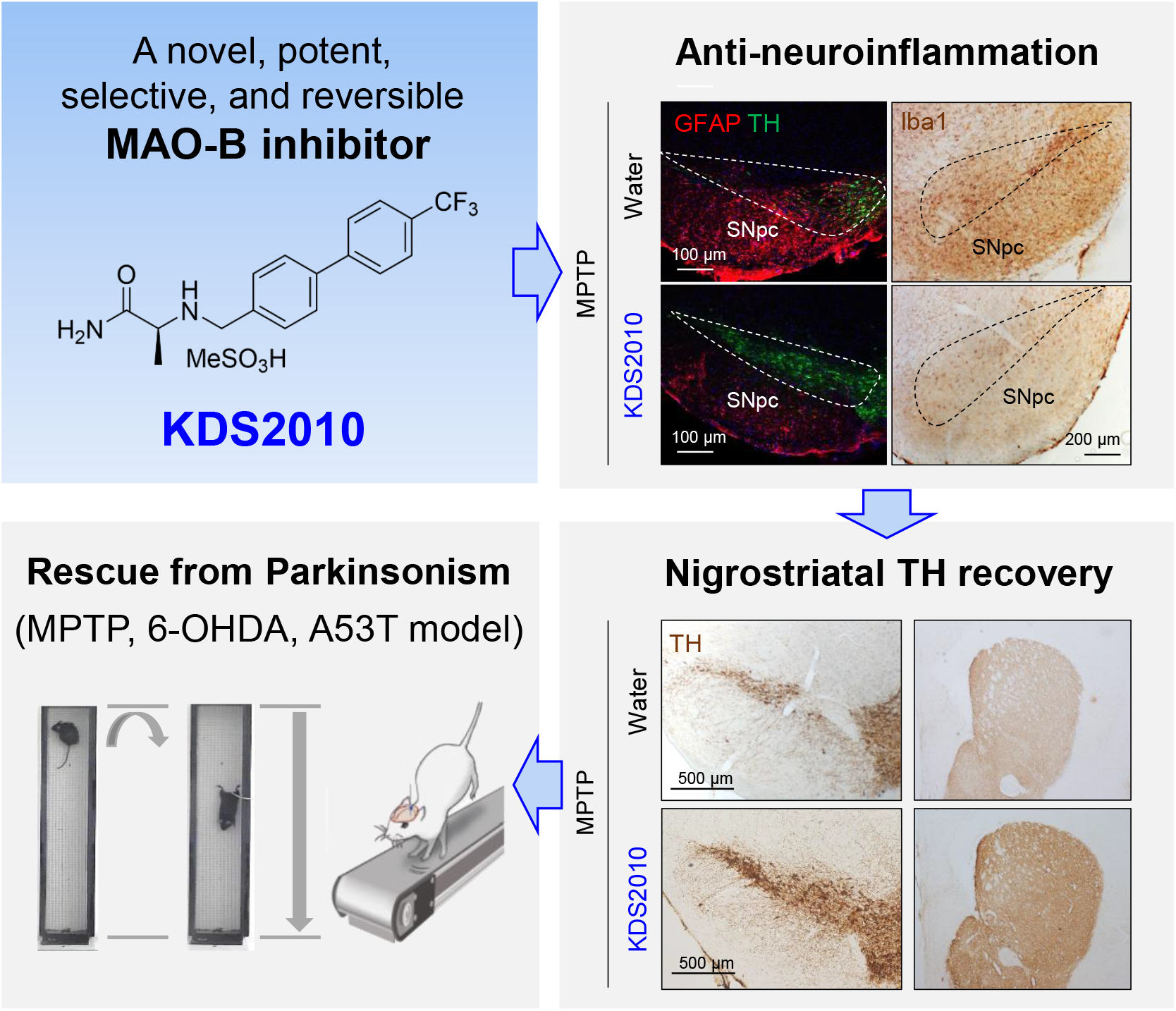

**What is already known:** KDS2010 is a recently developed potent, selective, and reversible MAO-B inhibitor.

MAO-B is critical for PD pathology through astrocytic GABA and H_2_O_2_ synthesis.

**What this study adds:** KDS2010 treatment dramatically recovers from PD-related pathology and motor deficit after pre- and post-treatment regimes in several animal models of PD.

KDS2010 exhibits low toxicity and excellent pharmacokinetic profile in non-human primates.

**What is the clinical significance?:** KDS2010 is a safe and promising therapeutic candidate for Parkinson’s disease.

Reversible MAO-B inhibitors could be more effective for treatment of Parkinson’s disease, overcoming the short-lived actions of irreversible MAO-B inhibitors.

## 1 Introduction

Parkinson’s disease (PD) is a neurodegenerative disorder with motor dysfunctions such as rigidity, bradykinesia, tremor at rest, and gait disturbance (Dickson, 2012). These motor symptoms, also called, parkinsonism, are caused by dopamine deficiency at the nigrostriatal pathway (Dickson, 2012). For alleviating these motor symptoms of patients with PD, various drugs have been developed and prescribed in clinics including dopamine precursors (Nagatsua & Sawadab, 2009), dopamine receptor agonists (Foley, Gerlach, Double & Riederer, 2004), catechol-*O*-methyl transferase (COMT) inhibitors (Widnell & Comella, 2005), and monoamine oxidase B (MAO-B) inhibitors (Robottom, 2011). Among these drugs, MAO-B inhibitors are one of the most frequently prescribed drugs.

MAO-B, an enzyme located in the outer mitochondrial membrane of astrocytes in the brain, catalyzes the oxidative deamination of biogenic amines (Greenawalt & Schnaitman, 1970). Therefore, MAO-B has been believed to degrade dopamine which is a monoamine neurotransmitter and MAO-B inhibitors have been believed to exert anti-Parkinsonian effects by blocking the degradation of dopamine (Schapira, 2011). In contrast, recent studies have demonstrated that MAO-B expression is dramatically increased in reactive astrocytes and that MAO-B is responsible for astrocytic GABA synthesis through the putrescine degradation pathway (Heo et al., 2020; Jo et al., 2014; Nam et al., 2020; Yoon et al., 2014a). A recent study revealed that MAO-B-mediated astrocytic GABA is critically involved in the pathology of PD. In detail, excessive GABA from reactive astrocytes in the substantia nigra pars compacta (SNpc) tonically inhibits neighboring dopaminergic (DA) neurons to suppress the synthesis and release of dopamine (Heo et al., 2020). These recent previous findings raise a possibility of an alternative mode-of-action of MAO-B inhibitors in PD through inhibition of astrocytic GABA synthesis.

Until now, several irreversible MAO-B inhibitors, such as selegiline and rasagiline, have been developed and prescribed to the patients with PD in clinics. Even though they have been reported to be effective in delaying the need for levodopa treatment, several studies still have cast doubt on the efficacy and the long-term balance of benefits and risks (Ives et al., 2004; Olanow et al., 2009), which is possibly related to their irreversibility. To overcome this shortfall of irreversible MAO-B inhibitors, a reversible MAO-B inhibitor, safinamide, was developed as a PD drug. However, safinamide is known to have some undesirable actions including a blockade of voltage-gated sodium and calcium channels, which possibly causes severe adverse events (Marzo et al., 2004). Meanwhile, we recently developed a potent, highly selective, and reversible MAO-B inhibitor, KDS2010, which is an α-amino amide derivative containing a biphenyl moiety. KDS2010 was reported to circumvent the shortcomings of irreversible MAO-B inhibitors and significantly rescue the memory deficit in a mouse model of Alzheimer’s disease (Park et al., 2019). In the current study, we screened more α-amino amide derivatives to search for more potent and selective MAO-B inhibitor candidates and tested the therapeutic potential of KDS2010 in several PD animal models. Furthermore, we studied the pharmacokinetics and toxicity of KDS2010 in non-human primates to validate its safety as a clinical drug.

## 2 Methods

### 2.1 Chemical synthesis

Compound synthesis and analysis methods are listed in Supplemental Information.

### 2.2 *in vitro* MAO-A and MAO-B enzyme assay

Analysis of the 50% inhibitory concentration of compounds (IC_50_) for *in vitro* MAO-A and MAO-B enzyme activity was performed as described previously (Choi et al., 2015). In brief, human recombinant MAO-A (hMAO-A) and MAO-B (hMAO-B) enzymes (Sigma-Aldrich) were diluted in 50 mM phosphate buffer (final protein amount, ~ 0.3 μg protein/well for MAO-A and ~ 2.5 μg protein /well for MAO-B), the test compound was treated after dissolving in DMSO (0.1 nM - 10 μM). The amount of hydrogen peroxide (H_2_O_2_) released after reaction with the substrate (*p*-tyramine for MAO-A; benzylamine for MAO-B) was quantified using a microplate fluorescence reader.

### 2.3. Animals

All mice were kept in a temperature- and humidity-controlled environment with a 12-hour light/12-hour dark cycle, and mice had free access to food and water. Handling and animal care were performed according to the directives of the Animal Care and Use Committee of the Institutional Animal Care and Use Committee of Korea Institute of Science and Technology (KIST) (Seoul, Korea). In all animal experiments, mice were treated by oral administration with each concentration for KDS2010, selegiline, or safinamide at 10 mg/kg per day. The amount of each drug (in mg) was calculated according to each mouse’s weight (in kg) and was dissolved in 100 μl of water. Each oral administration was performed once per day.

All MPTP experiments used the acute regimen consisting of four times of intraperitoneal injection of MPTP-HCl (M0896, Sigma Aldrich, 2 mg/mL in saline, 20 mg/kg for one injection) in a day with 2 h intervals. MPTP use and safety precautions were strictly followed as previously described (Jackson-Lewis & Przedborski, 2007).

6-OHDA model was prepared as previously described (Yoon et al., 2016). Under general anesthesia, all rats received unilateral injections of 8 mg 6-OHDA (Sigma-Aldrich) in 4 mL of saline with 0.1% ascorbic acid into the right medial forebrain bundle (AP −2.2 mm, L +1.5 mm relative to the bregma, and V −8.0 mm from the dura) (Paxinos & Watson, 1998), with the tooth bar set at +4.5 mm. First, to confirm if 6-OHDA model is successfully prepared, the apomorphine (0.25 mg/kg, s.c. administration, Sigma aldrich)-induced rotation test was performed with automated Rotameter (Panlab, Barcelona, Spain). Inclusion criteria was above 6 rpm.

Stereotaxic injections of AAV-CMV::A53T or control AAV-CMV::eGFP virus (packaged by KIST Virus Facility) were performed (right mouse SN: AP −3.2 mm, ML −1.3 mm, DV −4.0 mm; right rat SN; AP −5.3 mm, ML −2.3 mm, DV −7.6 mm relative to the bregma; 0.2 mL/min, total 2 mL) (Paxinos & Watson, 1998) under general anesthesia induced by chloral hydrate (rat) (Sato et al., 2011).

### 2.4 MAO-B assay with brain tissues

Measurements of MAO-B enzyme activity in tissue of 6-OHDA model was assessed using a commercially available kit (A12214, Thermo Fisher Scientific), following the manufacturer’s protocol. To obtain the mitochondrial fraction from the SNpc tissue, the tissues were homogenized in buffer A solution (250 mM sucrose, 2 mM HEPES (pH 7.4), 0.1 mM EGTA). The homogenized tissues were centrifuged at 571 G for 10 min and the supernatants were centrifuged at 14290 G for 10 min. Then, the pellet was re-suspended in buffer B solution (25 mM potassium phosphate, 5 mM MgCl_2_) and centrifuged at 15339 G for 10 min. The pellet was re-suspended in reaction buffer (0.05 M sodium phosphate). Protein concentration was determined using a BCA (Bicinchoninic acid) protein assay kit (#23228 and #23224, Thermo Scientific). The fluorimetric assay to assess the MAO-B activity was initiated by adding 100 μl of a reaction mixture containing Amplex Red reagent (400 μM), horseradish peroxidase (HRP; 2 U/ml) and benzylamine (2 mM), a specific substrate of MAO-B. Plates were incubated for 30 minutes at 37°C, protected from light, and the absorbance was measured at 570 nm using a microplate reader (Infinite M 200 Pro, Tecan). H_2_O_2_ (10 μM) was used as a positive control, and reaction buffer alone was used as a negative control.

### 2.5 DAB staining

The 30-μm-thick coronal sections for striatum and SNpc were immunostained with a DAB staining kit (TL-060-QHD, Thermo, MA, USA). The sections were incubated in Hydrogen Peroxide Block (TA-060-HP, Thermo) for 10 min, washed in PBS 3 times, incubated for 5 min in Ultravision Block (TA-060-UB, Thermo) and washed in PBS 3 times again. Then the samples were immunostained with a primary antibody (Rabbit anti-TH, Pel-freez, p40101-0, 1:500; Rabbit anti-Iba1, Wako, 019-19741, 1:500) in a blocking solution (0.3% Triton-X, 2% ready-to-use donkey serum (GTX30972, Genetex, CA, USA) in 0.1 M PBS) at 4°C on a shaker overnight. After washing in PBS 3 times, sections were incubated in Primary Antibody Amplifier Quanto (TA-060-QPB, Thermo) for 5 min, and washed in PBS again. The sections were incubated in HRP Polymer Quanto for 1 h and washed 4 times in PBS. DAB+ chromogen and DAB+ substrate buffer (K3468, Dako, Denmark) were mixed in 1:10 ratio and the sections were dipped in the mixture for 30 seconds and then washed. Finally, sections were mounted with mounting solution and dried. A series of bright field images were obtained with an Olympus microscope.

An unbiased stereological estimation of the total number of TH+ neurons in the SN area was performed using the optical fractionator method. The counted sections covered the entire SN, from the rostral tip of the SNpc to the caudal end of the SNpr (substantia nigra pars reticulate). An unbiased counting frame of known area (47.87 × 36.19 μm = 1733 μm^2^) was placed randomly on the first counting area and systematically moved through all counting areas until the entire delineated area was sampled. Counting was performed using a low-magnification objective lens (×10). The total number of neurons was estimated according to the optical fractionator formula.

### 2.6 Slice immunostaining for confocal microscopy

Sections were first incubated for 1 h in a blocking solution (0.3% Triton-X, 2% normal serum in 0.1 M PBS) and then immunostained with a mixture of primary antibodies (Rabbit anti-TH, Pel-freez, p40101-0, 1:500; Chicken anti-GFAP, Millipore, AB5541, 1:500) in a blocking solution at 4°C. After extensive washing, sections were incubated with corresponding fluorescent secondary antibodies for 2 h and then washed with PBS 3 times. Finally, sections were mounted with fluorescent mounting medium (S3023, Dako) and dried. A series of fluorescent images were obtained with an A1 Nikon confocal microscope, and Z-stack images in 3-μm steps were processed for further analysis using ImageJ program (NIH, MD, USA). Any alterations in brightness or contrast were equally applied to the entire image set. Specificity of primary antibody and immunoreaction was confirmed by omitting primary antibodies or changing fluorescent probes of the secondary antibodies.

### 2.7 Behavioral tests

To assess the motor deficits of MPTP-induced mouse model of PD, we performed vertical grid test as described in the previous study (Kim, Son, Choi, Ji & Hwang, 2010). Briefly, a mouse was gently placed inside the apparatus at 3 cm from the top, facing upward, and was allowed to turn around and climb down after 2-day habituation. To assess the motor deficits of 6-OHDA or A53T-mutated alpha-synuclein overexpression-induced rat models of PD, stepping test was performed as previously described with slight modifications (Yoon et al., 2014b). Briefly, both hind limbs and one forelimb were firmly fixed in the two hands of the experimenter, and the rostral part of the rat was lowered onto a treadmill moving at rate of 18 cm/s. The rat’s body remained stationary while the unilateral forelimb was allowed to spontaneously touch the moving treadmill track for 10 s. All of the experiment sessions were video-recorded to allow the number of adjusted steps taken in the backward direction to be counted. The number of adjusting steps was averaged across the four trials in each session.

### 2.8 Pharmacokinetics and toxicity study in non-human primates

All non-human primate studies were conducted in the GLP (Good Laboratory Practice)-level laboratory at Korea Institute of Toxicology (KIT, Jeollabuk-do, Korea) and approved by their Institutional Animal Care and Use Committee. Cynomolgus monkeys (*Macaca fascicularis*; from Nafo Vanny, Dong Nai Province, Vietnam) were at least 24 month-old. Cynomolgus monkeys were housed individually in a stainless cage (543W × 715L × 818H in mm) during acclimation, pre-treatment and dosing periods. The environment of the animal room was automatically controlled according to the Standard Operating Procedures (SOPs) of KIT (target range: temperature 20—29°C, relative humidity 30—70%, approximately 12-hour light cycle with 300—700 Lux, ventilation 10—20 times/hour, and air pressure: negative pressure). Male and female were assigned to treatment groups in a stratified manner using the Pristima System based on the most recent body weight. Clinical signs including mortality, moribundity, general appearance and behavior changes were recorded with date/time of finding, and duration. Body weights were measured prior to dosing on the day of dosing. For oral administration, KDS2010 was dissolved in water. We conducted four cynomologus studies as listed in Table 2 and described below:

**Table 1.**
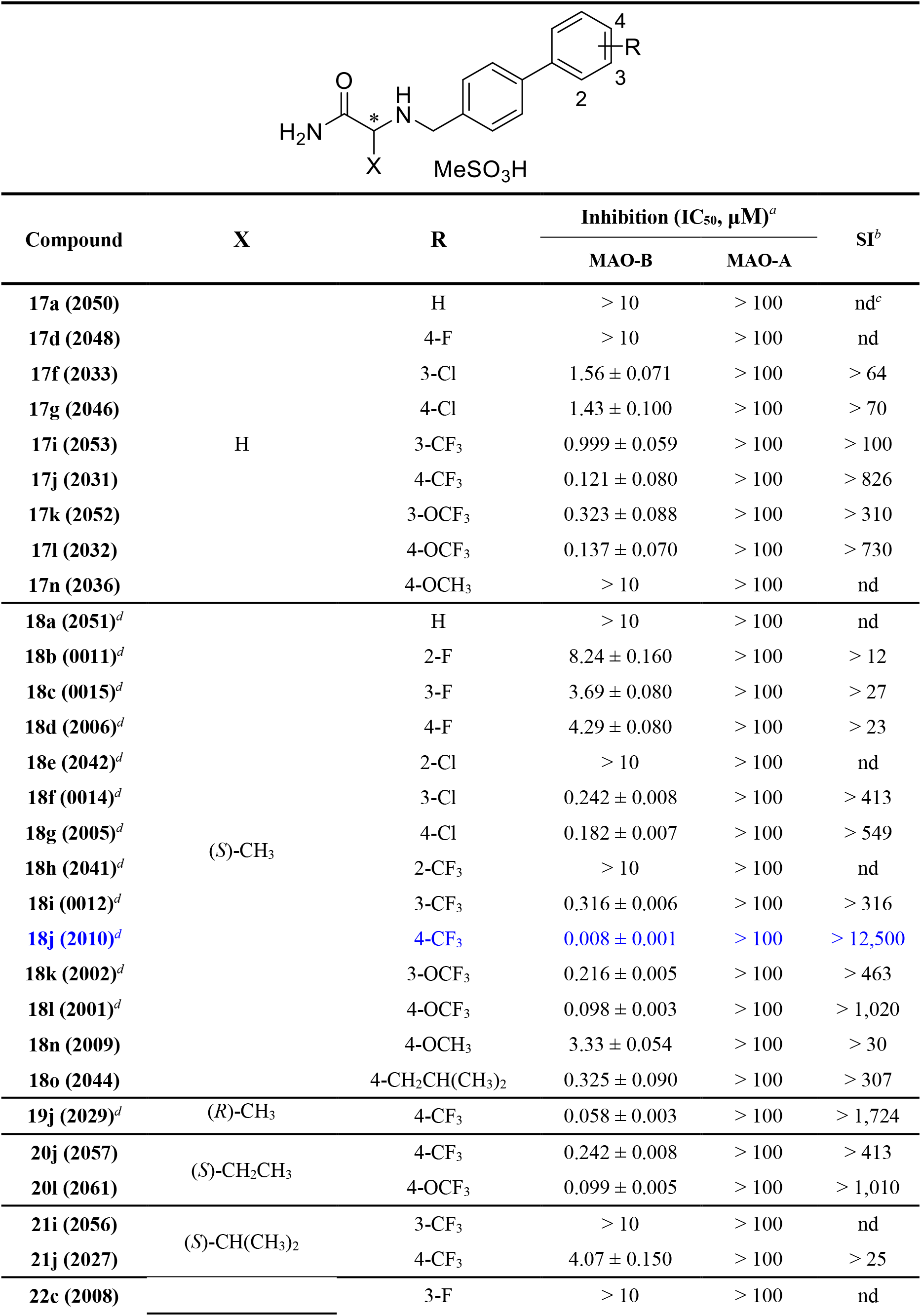
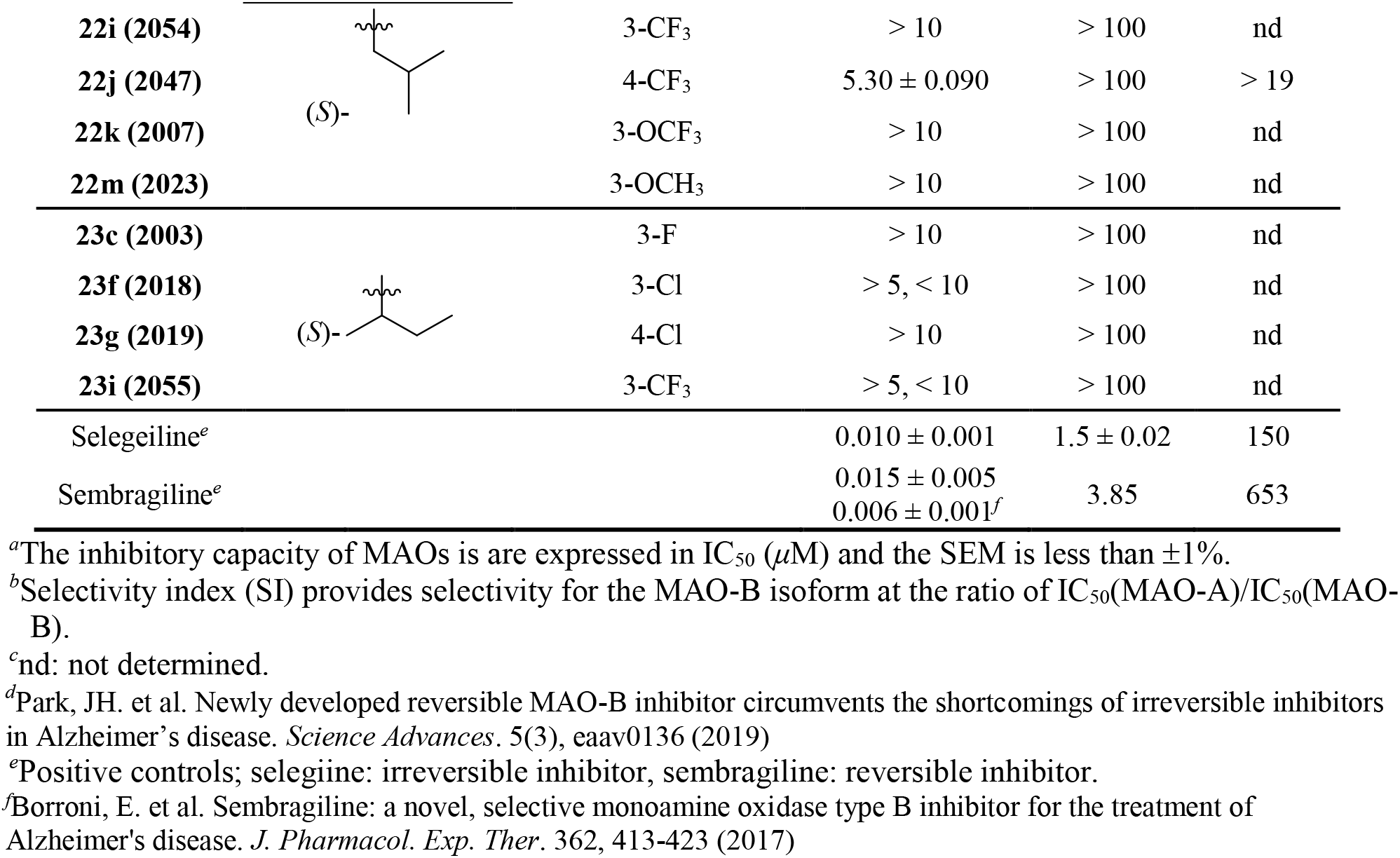
Inhibitory effects of α-aminoamide derivatives on hMAO enzymes

**Table 2.** Overview of non-human primates (cynomolgus monkey) studies

| Study No. | Animals per group<br>Main/recovery | Duration of dosing | Doses<br>(mg/kg/day) | Terminal<br>Sacrifice day |  |
| --- | --- | --- | --- | --- | --- |
| N118029 | 3 males/- | 1 time | 10 *p.o.* | - | PK |
|  | 3 males/- |  | 30 *p.o.* |  |  |
| N217019 | 1 male/-<br>1 female/- | 1 time in 1<sup>st</sup> week | 25 *p.o.* | 22 | Single dose-<br>escalation |
|  |  | 1 time in 2<sup>nd</sup> week | 50 *p.o.* |  |  |
|  |  | 1 time in 3<sup>rd</sup> week | 100 *p.o.* |  |  |
| G217013 | 4 males/- | Once a day | 0, 25, 50, 100 | 15 | Dose range-<br>finding |
|  | 4 females/- | for 2 weeks | *p.o.* |  |  |
| G218027 | 16 males/4 males<br>16 females/4 females | Once a day | 0, 10, 20, 40<br>*p.o.* | 22/36 | Repeated toxicity |
|  |  | for 4 weeks |  |  | for 4 weeks |
|  |  | 2 weeks for recovery |  |  | Recovery<br>for 2 weeks |
|  |  |  |  |  | Toxicokinetics |

In the pharmacokinetic study (**N118029**), six male monkeys were assigned into 2 groups. The animals were orally administered once with KDS2010 at dose levels of 10 or 30 mg/kg. Approximately 0.5 mL of blood was collected from the cephalic vein at several time points: pre-dose (0), 0.5, 1, 2, 4, 6, 8, 10, and 24-hour post-dose. The collected blood was put into the tubes with anti-coagulant (EDTA-2K). Blood samples were mixed gently and stored in wet-ice/cryo rack and then separated by centrifugation (approximately 13,200 rpm, 5 min, 4°C) to obtain plasma. The separated plasma was aliquoted into polypropylene tubes and stored frozen until analysis.

In the single-dose escalation study (**N118029**), each male and female monkey was orally administered with KDS2010 at increasing doses of 25, 50 and 100 mg/kg/day once a week. Mortality and general symptoms were observed during the trial period. Body weight was also measured. Blood was collected for hematology, coagulation and clinical chemistry tests at several time points: pre-dose (0), 8, 15 and 22-day post-dose. After administration of 100 mg/kg/day (day 22), an autopsy was performed to observe individual macroscopic findings.

In the repeated 2-week oral administration toxicity study (**G217013**) for 4-week dose range finding (DRF), monkeys were assigned to 4 groups of one male and one female. Each group was orally administered with KDS2010 at doses of 0, 25, 50 or 100 mg/kg/day once a day for 2 weeks. During the test period, general symptoms were observed, weight and feed intake was measured, and ophthalmological tests were performed. Clinical pathological tests (hematology, coagulation, blood biochemistry, and urine tests) were also conducted. After performing autopsy on all animals, individual macroscopic findings were recorded, organ weights were measured and pathological histologic examination was performed.

In the repeated 4-week oral administration toxicity study with 2-week recovery test and toxicokinetic test (**G218027**), monkeys were assigned to 4 groups with 3 males and 3 females. Each group was orally administered with KDS2010 at doses of 0, 10, 20 or 40 mg/kg/day once a day for 4 weeks. The experiments for toxicity test were conducted as described for the study G217013. For toxicokinetics, the blood of all animals were collected at pre-dose (0) and 1, 2, 4, 6, 8, 10, 24-hour after oral administration on the first day (Day 1) and the last day (Week 4; Day 28).

A non-compartmental analysis module in Phoenix^®^ WinNonlin^®^ (version 6.4) (Certara Inc., CA, USA) was used to calculate PK parameters. Systemic exposure to KDS2010 was calculated by applying the linear trapezoidal rule to the area under the plasma concentration-time curve from the time zero to the last quantifiable time-point (*AUC*_last_), and the maximum observed peak plasma concentration (*C*_max_) and the time to reach *C*_max_ (*T*_max_) were determined based on the observed data. The apparent terminal elimination half-life (*t*_1/2_) was calculated via apparent terminal elimination rate constant using [*t*_1/2_ = 0.693/kel].

## 3 Results

### 3.1 KDS2010, the most potent and selective MAO-B inhibitor among α-amino amide derivatives

In the previous study, to develop a novel MAO-B inhibitor we synthesized α-amino amide derivatives by introducing biphenyl groups with various substituents (Park et al., 2019). And we demonstrated that the introduction of electron-withdrawing group at the para-position is key to the MAO-B inhibitory effects. In this study, we designed and synthesized various derivatives by diversifying the alkyl group of α-amino amide compounds (Scheme 1). The synthetic pathways for α-amino amides (Compound # **17**-**23**) containing both various alkyl groups at X position and various functional groups on biphenyl ring B are displayed in Scheme 1.

We first replaced the hydrogen group at the X position instead of the methyl group which was introduced in the previously synthesized derivatives and compared their inhibitory effects on MAO-B (Table 1). Substitution with a H instead of CH_3_ group in compound **18j** (KDS2010) resulted in a 15-fold reduction in the MAO-B inhibitory effect (**17j**: 121 nM vs **18j**: 8 nM). Next, we introduced bulky alkyl groups, such as ethyl, isopropyl, isobutyl and sec-butyl groups to the X position, and found that the inhibitory effect dramatically decreased with increasing alkyl group size. As in the previous study, the compounds with electron-withdrawing groups introduced at the para position of biphenyl ring B increased the MAO-B inhibitory effects in the order of CF_3_>OCF_3_>Cl>F. Whereas, the compounds with electron-donating groups (OCH_3_ and CH_2_CH(CH_3_)_2_) significantly reduced the inhibitory efficacy. Overall, **18j** (KDS2010) showed the best inhibitory effect on MAO-B among the synthesized α-amino amide derivatives, and much greater selectivity for MAO-A than the well-known MAO-B inhibitors (selegiline and sembragiline). Furthermore, the (*S*)-stereoisomer (**18j**) exhibited 8-times higher inhibitory effect than (*R*)-isomer (**19j**). Therefore, we selected KDS2010 (**18j**) to evaluate for *in vivo* therapeutic effect in PD using various PD-related mice models and to investigate the PK and toxicity in non-human primates for as a potential clinical drug candidate.

### 3.2 Short-term KDS2010 treatment alleviates Parkinsonism

To validate the *in vivo* efficacy of the KDS2010 in PD, we first utilized an MPTP (1-methyl-4-phenyl-1,2,3,5-tetrahydropyridine)-induced mouse model which is one of the most widely used animal models for PD (Jackson-Lewis & Przedborski, 2007). MPTP is known to be converted to MPP+, which is a neurotoxin causing a significant damage to DA neurons in SNpc, through the enzymatic action of MAO-B (Chiba, Trevor & Castagnoli, 1984). Therefore, the therapeutic action of MAO-B inhibitor in MPTP model of PD is believed to be mediated by blocking the conversion of MPTP to MPP+ in astrocytes (Meredith & Rademacher, 2011). In this regard, most of the previous studies, animals were treated with MAO-B inhibitors before MPTP administration (Heo et al., 2020; Nam et al., 2017; Yeon et al., 2018). According to the broadly known protocol, we also treated the animals with KDS2010 (10 mg/kg/day) for 3 consecutive days from one day before MPTP administration (pre-treatment; Figure 1b). To test if KDS2010 pre-treatment alleviates MPTP-induced parkinsonian motor symptoms, we performed vertical grid test, which allows a sensitive examination of MPTP-induced motor deficit in mice as previously described (Figure 1a) (Kim, Son, Choi, Ji & Hwang, 2010). As expected, the pre-treatment of KDS2010 significantly reduced the total time, time to turn and rate of missed steps during the vertical grid test (Figure 1c-e), indicating that KDS2010 pre-treatment alleviated the MPTP-induced parkinsonian motor dysfunction.

**Figure 1.**
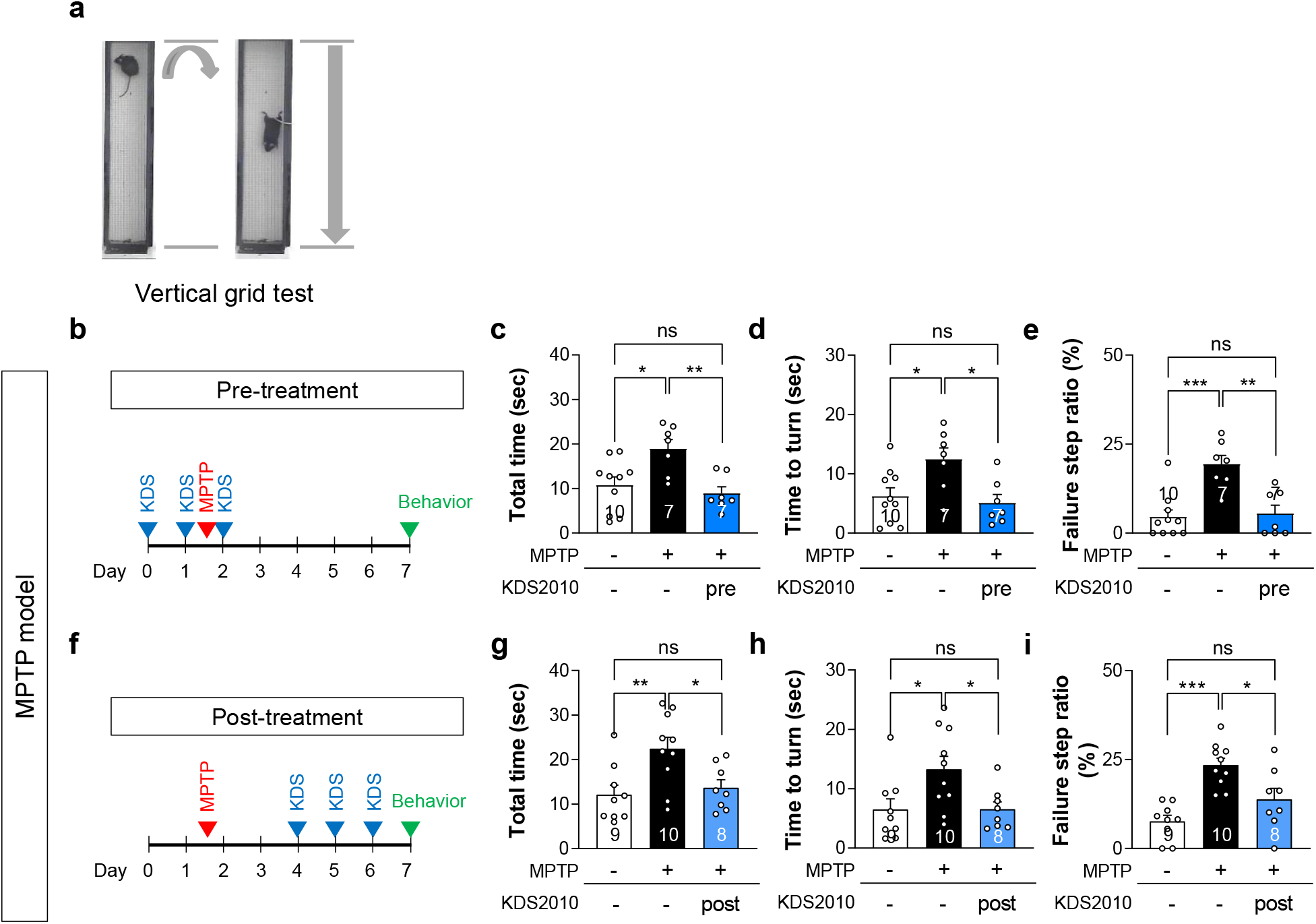
Pre- and post-treatments with KDS2010 alleviate parkinsonian motor deficits in MPTP model. (A) Schematic diagram of vertical grid test. (B) Timeline of pre-treatment with KDS2010. (C-E) Total time, time to turn, and failure step ratio assessed by vertical grid test. Pre-treatment with KDS2010 reduced total time, time to turn, and failure step ratio in MPTP-treated animals. (F) Timeline of post-treatment with KDS2010. (G-I) Total time, time to turn, and failure step ratio assessed by vertical grid test. Post-treatment with KDS2010 reduced total time, time to turn, and failure step ratio in MPTP-treated animals.

If the therapeutic effect of KDS2010 in MPTP-induced PD model is solely attributed to the blockade of MPTP conversion into MPP+, KDS2010 might not be an appropriate candidate for clinical PD drug. This is because there are many other causes of PD than MPTP, and because patients are not likely to be treated with MAO-B inhibitors before they are exposed to MPTP. Therefore, we tried to bypass the MAO-B’s action on MPTP to MPP+ conversion by treating with KDS2010 three days after MPTP treatment (post-treatment; Figure 1f). Three days are enough time for MPTP to be converted to MPP+ (Sundstrom, Luthman, Goldstein & Jonsson, 1988), so that the effect of KDS2010 should not be mediated by a blockade of MPTP conversion into MPP+. We found that the post-treatment of KDS2010 also significantly reduced the total time, time to turn and rate of missed steps during the vertical grid test (Figure 1g-i). Consistently, we have demonstrated that selegiline also showed a significant therapeutic effects in the MPTP model, by post-treatment as well as pre-treatment regime (Heo et al., 2020). These results indicate that the therapeutic effect of KDS2010 in MPTP model could be attributed to blocking other actions of MAO-B.

### 3.3 Short-term KDS2010 treatment alleviates nigrostriatal TH loss and neuroinflammation

To test whether the KDS2010-induced functional recovery in MPTP model is associated with nigrostriatal level of tyrosine hydroxylase (TH), the key dopamine-synthesizing enzyme, we assessed the nigrostriatal TH level of KDS2010-treated MPTP model. TH is widely accepted as a marker of DA neurons of SNpc projecting to the striatum. We found that MPTP administration significantly reduced the TH-positive cell number in SNpc and the optical density of TH in the striatum, which validates our MPTP model (Figure 2). We also found that both pre-treatment and post-treatment with KDS2010 significantly recovered the reduced TH-positive cell number in SNpc and the optical density of TH in the striatum (Figure 2). These findings indicate that the motor functional recovery by KDS2010 treatment is associated with nigrostriatal TH recovery.

**Figure 2.**
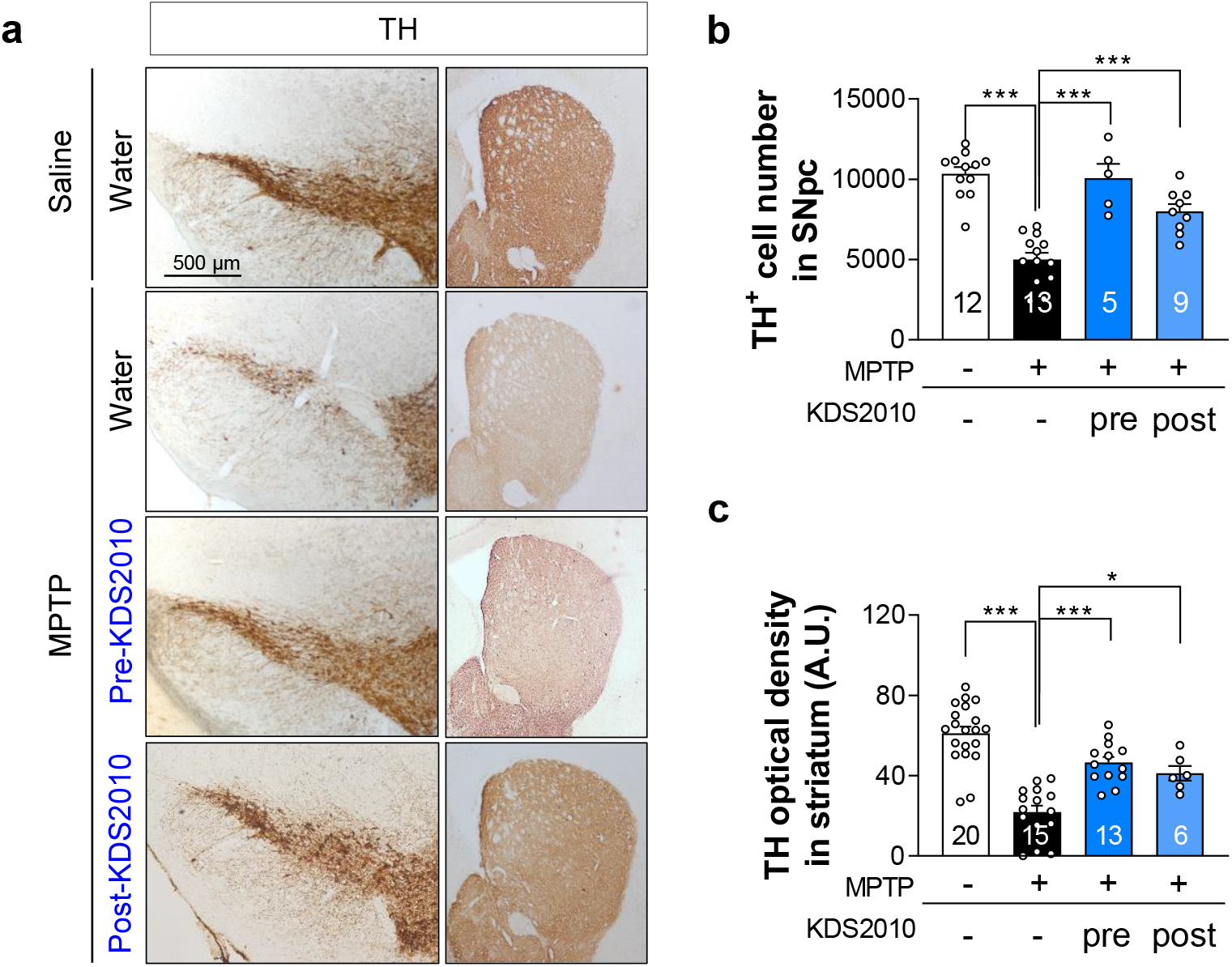
Pre-treatment and post-treatment with KDS2010 rescue nigrostriatal TH loss in MPTP model. (A) Representative images of TH-stained SNpc and striatum tissues. (B) Quantification of TH-positive cell number in SNpc. Both pre-treatment and post-treatment with KDS2010 significantly rescued the number of TH-positive dopaminergic neurons in SNpc of MPTP-treated animals. (C) Quantification of TH optical density in striatum. Both pre-treatment and post-treatment with KDS2010 significantly mitigates the TH loss in striatum of MPTP-treated animals.

In addition to nigrostriatal TH loss, neuroinflammation is a well-known pathology in PD (McGeer & McGeer, 2008). Indeed, it has been well documented that astrocytes and microglia are found to be reactive in the brains of PD animal models such as MPTP model and PD patients (Heo et al., 2020; Joe, Choi, An, Eun, Jou & Park, 2018). We previously reported that the increased astrocytic reactivity is mediated by MAO-B in the brains of PD as well as AD (Heo et al., 2020; Park et al., 2019), and that KDS2010 treatment dramatically reduces astrocytic reactivity in AD (Park et al., 2019). Therefore, we tested if KDS2010 treatment also reduces astrocytic as well as microglial reactivity in the MPTP model of PD. We performed immunohistochemistry with antibodies against glial fibrillary acidic protein (GFAP) and Iba1 for astrocytes and microglia, respectively. We found that MPTP treatment significant increased the intensities of both GFAP and Iba1, and KDS2010 significantly reduced both (Figure 3). These results are consistent with our previous results in AD (Park et al., 2019) and indicate that KDS2010 is effective in reducing neuroinflammation which can be critical for DA neuronal suppression and degeneration.

**Figure 3.**
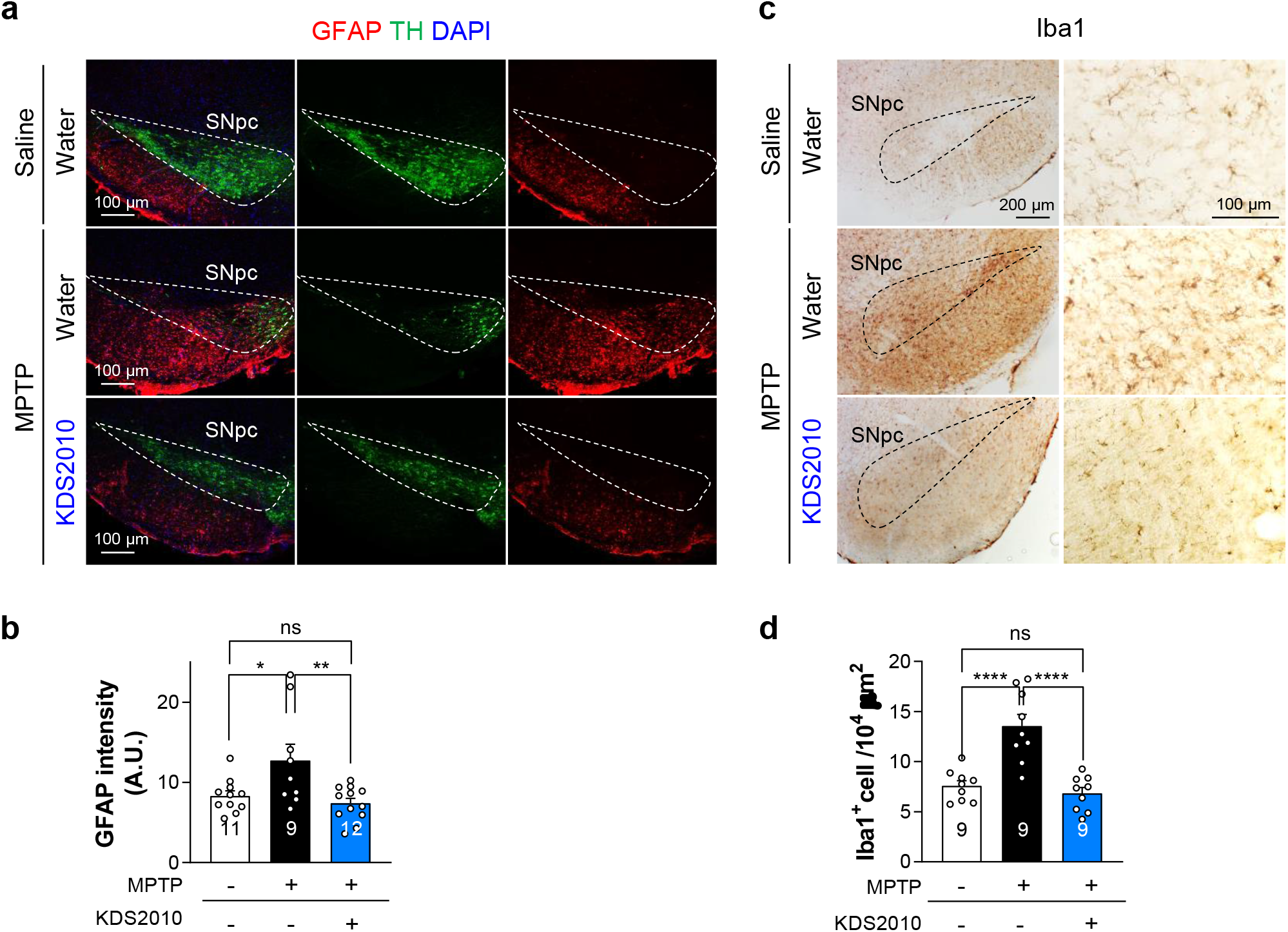
KDS2010 treatment alleviates glial reactivity. (A) Representative confocal images of GFAP and TH-stained SNpc tissues. (B) Quantification of GFAP intensity in SNpc area. MPTP significantly increased the GFAP intensity which was reversed by KDS2010 treatment. (C) Representative images of Iba1-stained SNpc tissues. (D) Quantification of Iba1 intensity in SNpc area. MPTP significantly increased the Iba1 intensity which was reversed by KDS2010 treatment.

### 3.4 Long-term KDS2010 treatment mitigates the PD-like pathology and symptoms in MPTP

We previously reported that irreversible MAO-B inhibitors have a critical drawback of turning on compensatory GABA synthetic mechanism when treated for a long period of time, while reversible MAO-B inhibitors do not (Park et al., 2019). The short-lived action of selegiline in blockade of astrocytic GABA synthesis is also attributed to its irreversibility (Jo et al., 2014; Park et al., 2019). Therefore, we also tested the long-term effect of KDS2010 in MPTP model, comparing with selegiline and safinamide, another reversible MAO-B inhibitor. We treated with the MAO-B inhibitors for 29 days before MPTP injection and the drug treatment was continued for one more day (Figure 4a). We found that KDS2010 and safinamide showed significant recovery of TH-positive cells in SNpc, while only partial recovery was found in selegiline-treated group (Figure 4b, c). In terms of motor recovery, total time and time to turn were significantly reduced by all three MAO-B inhibitors in MPTP model (Figure 4d, e), while the failure step ratio was only significantly reduced by reversible MAO-B inhibitors such as KDS2010 and safinamide, but not by selegiline (Figure 4f). These findings indicate that the reversible MAO-B inhibitors are more effective in mitigating PD-like pathology and symptoms in MPTP model when long-term treated, compared to the irreversible MAO-B inhibitor, selegiline.

**Figure 4.**
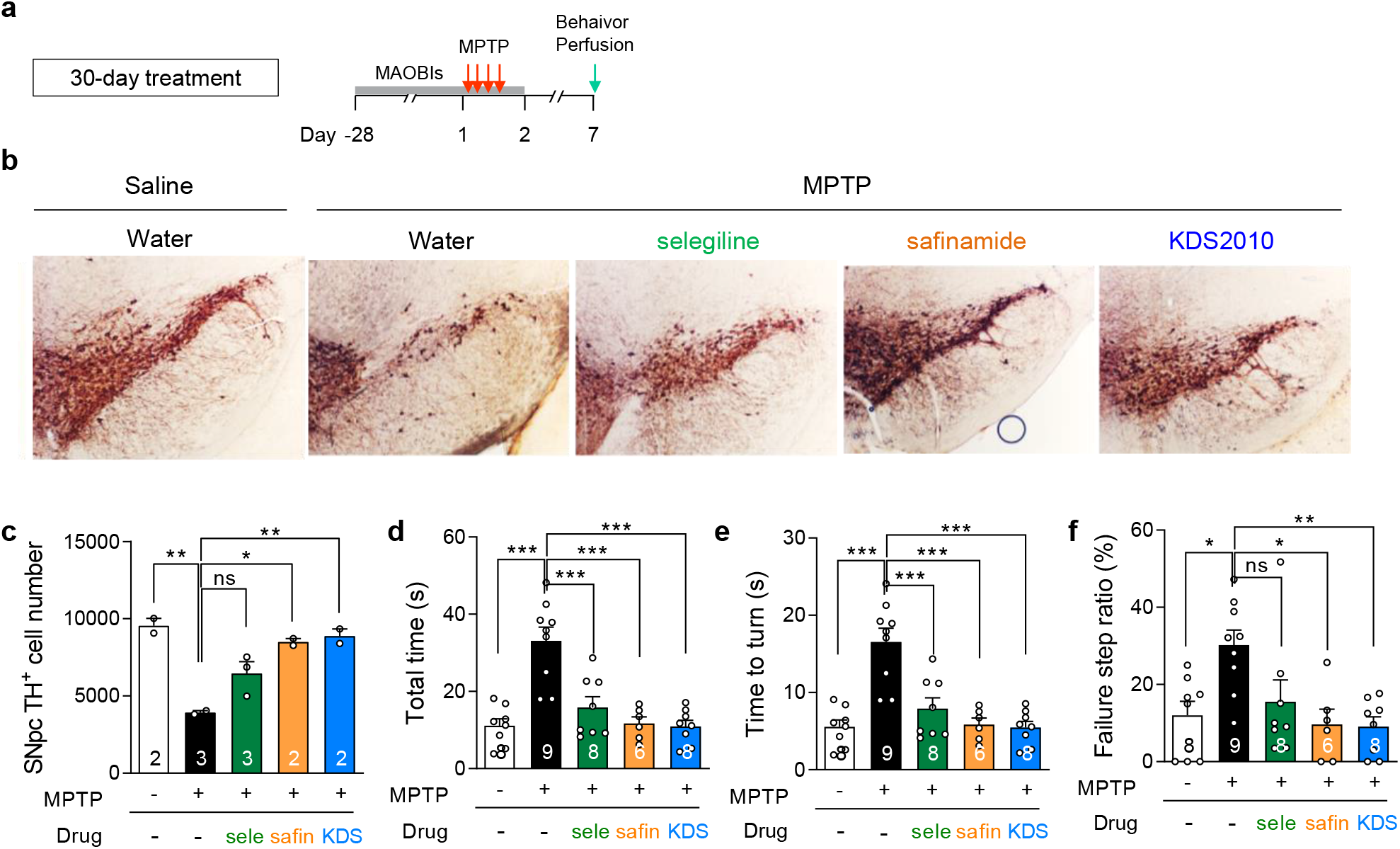
Long-term treatment of reversible MAO-B inhibitors significantly alleviates TH loss and parkinsonian motor deficits in MPTP model. (A) Timeline of long-term (30-day) treatment with MAO-B inhibitors, including selegiline, safinamide, and KDS2010. (B) Representitive images of TH-stained SNpc tissues. (C) Quantification of TH-positive cell number in SNpc. Long-term treatment with safianamide and KDS2010 significantly rescued the number of TH-positive dopaminergic neurons in SNpc of MPTP-treated animals, while selegiline only showed a non-significant trend of recovery. (D-F) Total time, time to turn, and failure step ratio assessed by vertical grid test. Long-term treatment with safianamide and KDS2010 significantly rescued all items assessed in MPTP-treated animals, while selegiline only showed a non-significant reducing trend in failure step ratio.

### 3.5 KDS2010 is effective in other animal models of PD

Next, we tested the therapeutic efficacy of KDS2010 in 6-OHDA rat model, which exhibits more extensive DA neurodegeneration and motor deficits. For assessment of motor behaviors of rats, we performed the stepping test which is well-validated for assessing the parkinsonian motor deficits (Yoon et al., 2014b) (Figure 5a). After pre-screening for properly generated 6-OHDA model animals with apomorphine-induced rotation test, the selected animals were treated with KDS2010 (10 mg/kg/day) for 15 days (Figure 5b). While vehicle-treated 6-OHDA model rats showed severely impaired stepping behavior of the contralateral forepaw, KDS2010-treated rats showed significantly improved stepping behavior (Figure 5c). This is in marked contrast to the results from the previous study, in which selegiline treatment did not show any improvement in stepping behavior (Heo et al., 2020).

**Figure 5.**
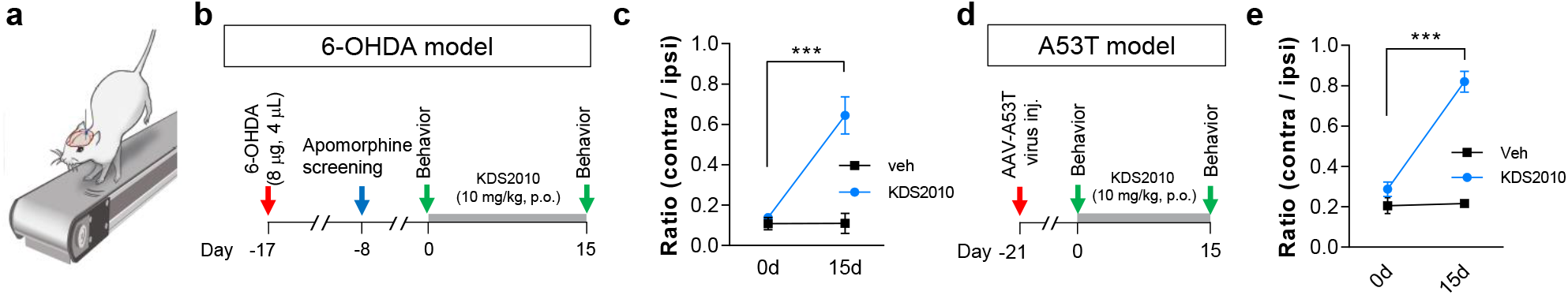
Treatment with KDS2010 alleviates parkinsonian motor deficits in 6-OHDA and A53T models. (A) Schematic diagram of stepping test. (B) Timeline of post-treatment with KDS2010 for 15 days in 6-OHDA model. (C) Quantification of the ratio of contralateral stepping numbers over ipsilateral ones. KDS2010 treatment significantly recovered the stepping ratio. (D) Timeline of post-treatment with KDS2010 for 15 days in A53T-SNCA over-expressed model of PD. (E) Quantification of the ratio of contralateral stepping numbers over ipsilateral ones. KDS2010 treatment significantly recovered the stepping ratio.

We also tested the therapeutic efficacy of KDS2010 in A53T-alpha-synuclein-overexpressed rat PD model (A53T rat model) (Sato et al., 2011). This model was generated by AAV-A53T virus injection into unilateral SNpc of a rat as previously described (Heo et al., 2020). Unlike MPTP and 6-OHDA models, this is an alpha-synuclein-induced PD model which can mimic human PD pathology better. The animals were also treated with KDS2010 (10 mg/kg/day) for 15 days (Figure 5d). We found that A53T rat models showed impaired stepping behavior in contralateral forepaw, whereas KDS2010 treatment significantly rescued it (Figure 5e). Taken together, these findings indicate that KDS2010 is highly effective for treating parkinsonian motor symptoms, regardless of which animal model is used.

### 3.6 KDS2010 shows an excellent PK profile and no toxicity issues in non-human primates

To select the accurate and safe dose for the first-in-human (FIH) study, we performed a pharmacokinetic (PK) test in cynomolgus monkeys based on non-clinical PK data performed in rats (Park et al., 2019). A PK analysis performed after a single oral administration of KDS2010 to male monkeys at doses of 10 and 30 mg/kg showed a *C*_max_ of 2232.0 ± 142.7 ng/mL and 4845.1.0 ± 567.5 ng/mL, respectively (Table 3). Particularly, the *C*_max_ of KDS2010 at 10 mg/kg in monkeys was 2.34-fold higher than the *C*_max_ in rats (952.1±80.3 ng/mL) (Table 3). In addition, when compared at the same dose (10 mg/kg), the total blood concentration (*AUC*_all_) in cynomolgus monkey was 6 times higher than that in rats, indicating that the drug exposure was significantly superior to oral administration of KDS2010 in primate than rodent (Table 3). Next, we conducted single- and 4-week repeated-dose oral toxicity studies. In a single dose toxicity study (N217019), 25, 50 and 100 mg/kg of KDS2010 were administered in escalating dose daily for 3 days. As a result, no change was observed in general symptoms and clinical pathological and gross organ examination. For 4-week repeated-dose oral toxicity study, we first conducted a 2-week dose range founding (DRF) study (G217013) and determined the maximum dose to be 50 mg/kg/day. KDS2010 was administered orally to male and female cynomolgus monkeys at a dose of 0, 10, 20 or 40 mg/kg/day, once a day, for 4 weeks. The toxicokinetic analysis revealed that the mean value of T_max_ was 4.72 ± 1.93 hours after administration (Figure 6a), and that the calculated systemic exposure (*AUC*_last_) of KDS2010 was dose-dependently increased in a slightly less than proportional manner (Figure 6b). During the test period, no significant changes in body weight were observed in all male and female groups (Figure 6c). Moreover, no adverse effect in relation to KDS2010 was observed in various toxicity examinations. Taken together, KDS2010 has the No Observed Adverse Effect Level (NOAEL) of 40 mg/kg/day for both sexes. These results indicate a high potential of KDS2010 as a clinical candidate drug for PD.

**Table 3.** *In vivo* pharmacokinetic parameters of KDS2010

| Compd. | PK<sup>a</sup> |  |  |  |
| --- | --- | --- | --- | --- |
|  | 10 mg/kg *p.o.* |  | 30 mg/kg *p.o.* |  |
|  | *C*<sub>max</sub> | *AUC*<sub>all</sub> | *C*<sub>max</sub> | *AUC*<sub>all</sub> |
|  | (ng/mL) | (ng*h/mL) | (ng/mL) | (ng*h/mL) |
| **KDS2010** | 2232.0±142.7 | 30587.0±421.6 | 4845.1.0±567.5 | 80770.0±5374.7 |

| Compd. | PK<sup>b</sup> |  |  |  |  |  |  |
| --- | --- | --- | --- | --- | --- | --- | --- |
|  | 1 mg/kg *i.v.* |  |  |  | 10 mg/kg *p.o.* |  | *F*<br>(%) |
|  | *AUC*<sub>all</sub> | *CL* | *V*<sub>ss</sub> | *t*<sub>1/2</sub> | *C*<sub>max</sub> | *AUC*<sub>all</sub> |  |
|  | (ng*h/mL) | (mL/min/kg) | (L/kg) | (h) | (ng/mL) | (ng*h/mL) |  |
| **KDS2010** | 421.1±42.6 | 39.9±4.0 | 10.1±0.8 | 3.3±0.2 | 952.1±80.3 | 5201.5±458 | 123.5 |

<sup>a</sup>Monkeys (*n* = 3) were dosed with 10 mg/kg for *p.o.* or 30 mg/kg for *p.o.*. Parameters were calculated from composite mean plasma concentration-time data. Data are expressed as the mean ± SD. %.
<sup>b</sup>Rats (*n* = 5) were dosed with 1 mg/kg for *i.v.* and 10 mg/kg for *p.o.*. Parameters were calculated from composite mean plasma concentration-time data. Data are expressed as the mean ± SD. %. from Park, JH. et al. Newly developed reversible MAO-B inhibitor circumvents the shortcomings of irreversible inhibitors in Alzheimer's disease. *Science Advances*. 5(3), eaav0136 (2019)
**Abbreviations:** Compd, compound; *AUC*, area under the plasma concentration–time curve; *CL*, time-averaged total body clearance; *V*<sub>ss</sub>, apparent volume of distribution at steady state; *t*<sub>1/2</sub>, elimination half-life; *C*<sub>max</sub>, maximum concentration of the drug; *F*, bioavailability

**Figure 6.**
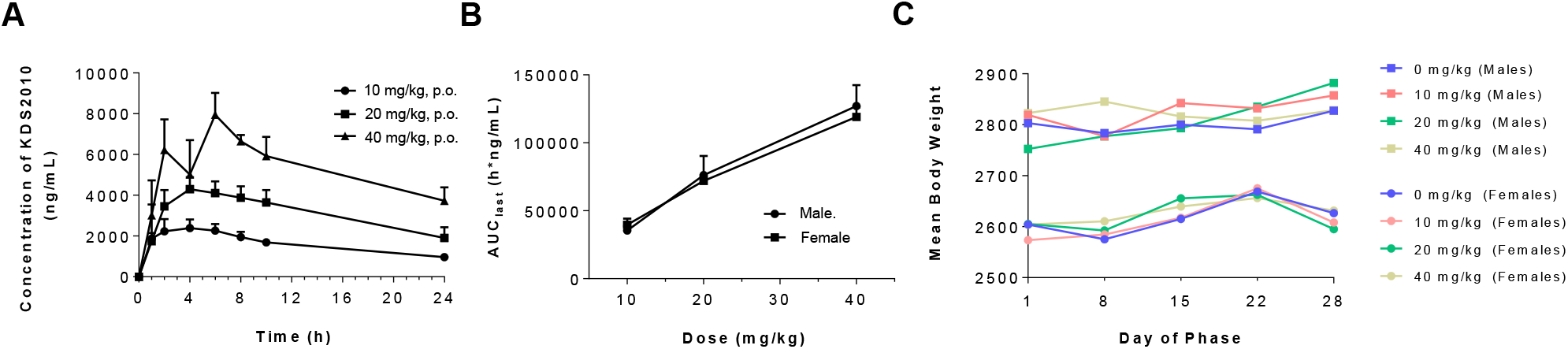
KDS2010 has an excellent PK profile in non-human primates and is safe for 4 weeks of repeated dosing. (A) Composite serum KDS2010 concentration-time profiles after single *p.o*. doses of 10, 20 and 40 mg/kg in cynomolgus monkey (B) Increased amount of systemic exposure (AUC_last_) compared to dose (C) Changes in body weight by group during the 4-week repeated dosing period

## 4 Discussion

In this study, we have demonstrated that KDS2010 is the most optimized MAO-B inhibitor among various alpha-amino amide derivatives, as revealed by the lowest IC_50_ and highest selectivity to MAO-B over MAO-A. Moreover, we found that 3-day treatment of KDS2010 treatment dramatically prevented or reversed the MPTP-induced PD-like pathology including nigrostriatal TH loss, neuroinflammation such as astrogliosis and microgliosis, and parkinsonian motor deficits. Particularly, in terms of long-term treatment, the therapeutic effect of KDS2010 in MPTP model was superior to selegiline, a widely prescribed irreversible MAO-B inhibitor. We also validated KDS2010 as a potential therapeutic agent for patients with PD with an excellent PK profile and safety in the extensive pharmacokinetics and toxicity studies with non-human primates.

In a previous study, we developed KDS2010, an α-amino amide derivative containing a biphenyl moiety, and confirmed the superb effect on MAO-B inhibition, selectivity to MAO-B, and reversibility (Park et al., 2019). Moreover, this novel compound exhibited the highest potency in both *in vitro* and *in vivo* rodent model systems and have also shown better ADME/Tox profiles than other chemical scaffolds (Park et al., 2019). The newly developed KDS2010 showed excellent PK and blood-brain-barrier permeability, molecular target specificity, and *in vivo* safety, which are important preclinical requirements for CNS drug candidates (Park et al., 2019).

What is the possible mechanism of the therapeutic effect of KDS2010 in *in vivo* PD models? First, the therapeutic effect of KDS2010 could be attributed to blockade of the conversion of MPTP to MPP+, because MPTP is reported to be converted to a neurotoxin, MPP+, by MAO-B in astrocytes (Chiba, Trevor & Castagnoli, 1984). This possibility can explain the effect of KDS2010 in pre-treatment regime. However, the post-treatment of KDS2010, which starts after 3 days from MPTP injection to allow a sufficient conversion of MPTP to MPP+, also dramatically reversed nigrostriatal TH level and parkinsonian motor symptoms. Our results raise an alternative action of MAO-B, other than the conversion of MPTP to MPP+. Second, the therapeutic effect of KDS2010 could be attributed to a blockade of dopamine degradation, because MAO-B is believed to be responsible for dopamine degradation. However, conflicting results have also been reported. For example, no difference in dopamine level between MAO-B-deficient mice and wild-type mice (Fornai, Chen, Giorgi, Gesi, Alessandri & Shih, 1999), and no alteration in dopamine level by selegiline treatment (Lamensdorf, Youdim & Finberg, 1996). In addition to these two possibilities, we have reported the critical role of MAO-B in GABA synthesis in reactive astrocytes (Heo et al., 2020; Jo et al., 2014; Nam et al., 2020; Yoon et al., 2014a). Reactive astrocytes produce excess amount of GABA through MAO-B and tonically release GABA to suppress neighboring DA neurons in SNpc, which leads to reduced TH level and DA synthesis, finally causing parkinsonian motor symptoms (Heo et al., 2020). We also demonstrated the evidence of TH-negative dormant DA neurons, which is suppressed by astrocytic GABA, in the SNpc of PD brains. Moreover, the effects of MAO-B inhibitors such as selegiline and safinamide in PD animal models were attributed to the blockade of tonic inhibition of SNpc DA neurons by inhibiting astrocytic GABA synthesis (Heo et al., 2020). Additionally, reactive astrocytes produce excessive amount of H_2_O_2_ when they synthesize GABA through MAO-B, leading to severe neuroinflammation and neuronal death. In this regard, the therapeutic effect of KDS2010 is also very likely to be mediated by a blockade of astrocytic GABA and H_2_O_2_ synthesis, rather than conversion of MPTP to MPP+ or clearance of dopamine.

The therapeutic effect of KDS2010 was previously tested in APP/PS1 transgenic mouse, a mouse model of AD. KDS2010 effectively ameliorated the memory impairment in APP/PS1 mice, regardless of whether they were treated with KDS2010 for short-term (3 days) or long-term (30 days). This effect was attributed to suppressing tonic inhibition of hippocampal neurons by blocking astrocytic GABA synthesis (Park et al., 2019). In the same study, we also found that long-term treatment (> 2 weeks) of selegiline did not show any therapeutic effect, which is due to compensatory up-regulation of astrocytic GABA synthesis (via diamine oxidase) following chronic treatment with an irreversible MAO-B inhibitor, while long-term treatment of reversible MAO-B inhibitor did not turn on this compensatory metabolism (Park et al., 2019). Furthermore, in our recent study a long-term treatment of KDS2010 showed a dramatic effect on post-stroke recovery when it was accompanied by rehabilitation training (Nam et al., 2020). The difference in therapeutic effects in AD between reversible and irreversible MAO-B inhibitors is consistent with current findings in PD. These findings suggest that reversible MAO-B inhibitor is more appropriate for clinical use, because most of the medicines are administered for long time to the patients with neurodegenerative disorders.

In summary, we have demonstrated that MAO-B is a key molecular element of astrogliosis in both PD and AD. Therefore, reversible MAO-B inhibitors are effective for alleviating and preventing astrogliosis. Furthermore, various brain disorders with astrogliosis could be possible indications for reversible MAO-B inhibitors. Among several reversible MAO-B inhibitors, KDS2010 has shown excellent potency and druggability with safety. We propose KDS2010 as a highly effective therapeutic candidate for Parkinson’s disease as well as other neuroinflammatory brain disorders.

## Acknowledgements

This work is supported by Institute for Basic Science (IBS), Center for Cognition and Sociality (IBS-R001-D2) to C.J.L.; and the National Research Council of Science & Technology (NST) grant by the Korea government (MSIP) (No. CRC-15-04-KIST), the National Research Foundation of Korea (NRF-2018M3A9C8016849) to K.D.P.

## Author contributions

M.H.N., J.H.P., K.D.P. and C.J.L. conceived and designed the research. M.H.N., J.H.P., S.K. and J.Y.H. and S.J.O. performed *in vivo* study. H.J.S. and B.K.J. synthesized the chemicals. J.H.P. and J.W.C. performed pharmacological assays. D.W.C., Y.Y.S. and S.C.H. performed toxicity test in non-human primates. M.H.N., J.H.P., K.D.P. and C.J.L. wrote the manuscript. All authors provided ongoing critical review of results and commented on the manuscript.

## Declaration of transparency and scientific rigour

This Declaration acknowledges that this paper adheres to the principles for transparent reporting and scientific rigour of preclinical research as stated in the BJP guidelines for Design & Analysis and Animal Experimentation, and as recommended by funding agencies, publishers and other organisations engaged with supporting research

## Data availability

All data underpinning this study is available from the authors upon reasonable request.

---

**Scheme 1.**
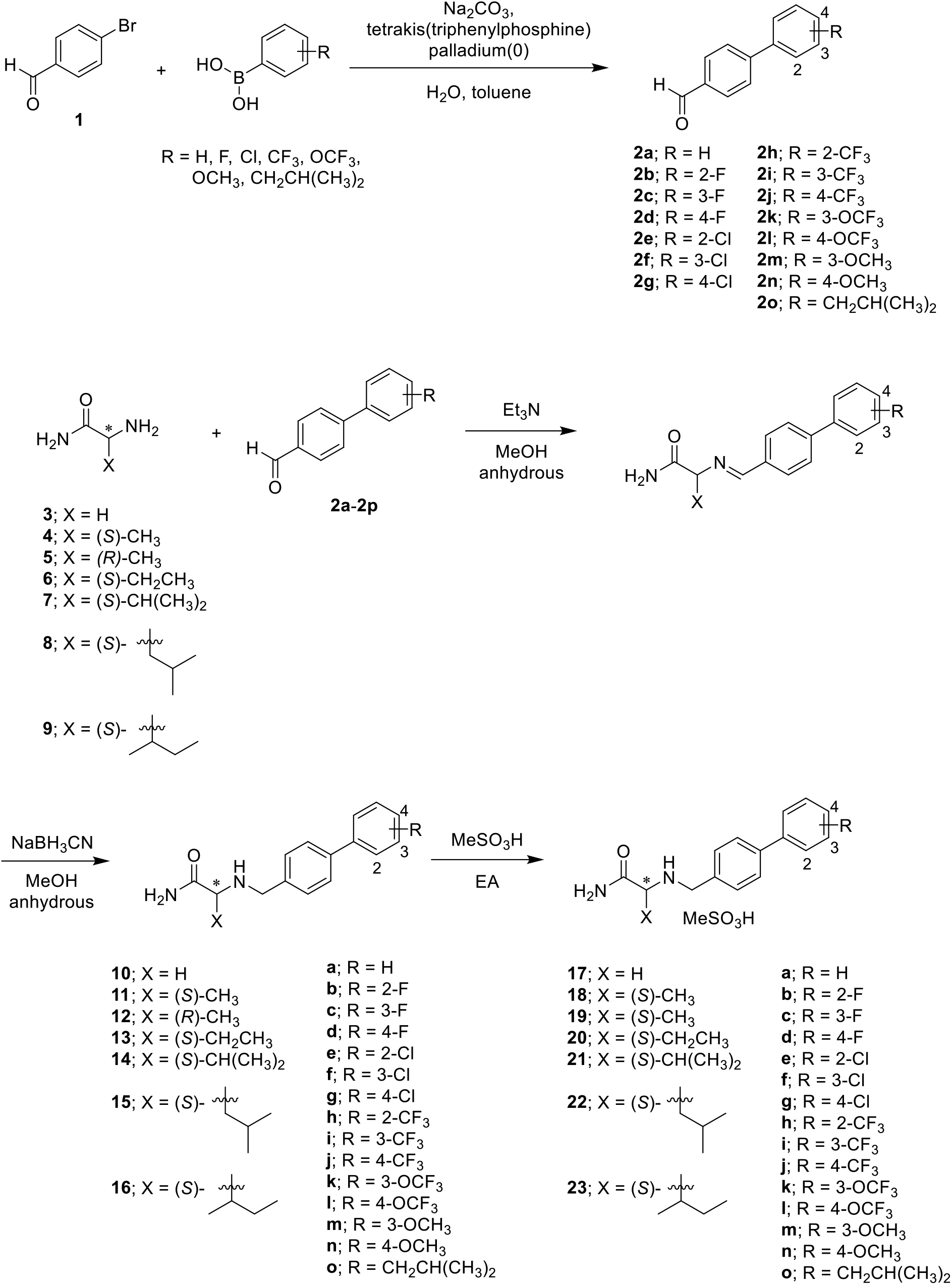
Synthetic scheme for α-aminoamide derivatives **17**–**23**

