## Supplementary materials for "KDS2010, a newly developed reversible MAO-B inhibitor, as an effective therapeutic candidate for Parkinson’s disease"

### EXPERIMENTAL SECTION

#### General Methods

The melting points were measured in capillary tubes using a Standford Research System Opti melt apparatus. The reactions were checked by analytical thin-layer chromatography (TLC) (Merck, Cat # 1.05715) and analyzed with ultraviolet light (254 nm) and were purified by flash column chromatography using silica gel (Merck, Cat # 1.07734 & 1.09385). NMR spectra were obtained at either 300 MHz ($^{1}H$) and 75 MHz ($^{13}C$) or 400 MHz ($^{1}H$) and 100 MHz ($^{13}C$) using BRUKER apparatus. The chemical shifts ($\delta$) were calibarated from tetramethylsilane (TMS) in parts per million (ppm) and the coupling constants ($J$) were reported in hertz and assigned as follows : singlet (s), doublet (d), triplet (t), quartet (q) and broad singlet (br s). All reagents and solvents used as purchased from commercial sources without further purification. HPLC was analyzed using a Waters E2695 system provided with the following column: SHISEIDO capcell pak $C_{18}$ MG II column (4.6 mm × 150 mm; 5 μm) and SHISEIDO capcell pak® column (4.6 mm × 75 mm; 3μm). HPLC data were recorded using following parameters: 1% acetic acid in $H_2O$/MeCN, 90/10 → 0/100 in 10 min, + 10 min isocratic, flow rate of 1 mL/min, λ = 254 and 280 nm. Compounds were confirmed by TLC, $^{1}H$ and $^{13}C$ NMR. The TLC, NMR, HPLC and the analytical data checked the purity of the products was ≥ 95%.

#### General procedure for the preparation of aldehyde compounds (2a-2p) (Method A)

A mixture of 4-bromobenzaldehyde, the desired arylboronic acid (1.3 equiv), tetrakis(triphenylphosphine)palladium(0) (4–8 mol%), $Na_2CO_3$ (4.9 equiv) in degassed toluene/$H_2O$ (7/3) was refluxed 18 h. The mixture was concentrated *in vacuo* and the resulting solution was washed with EtOAc (200 mL) and $H_2O$ (2 × 200 mL). The organic layer was dried with anhydrous $Na_2SO_4$ and concentrated *in vacuo*. The residue was purified by column chromatography on Silica gel.

#### General procedure for the preparation of free amine compounds (10a-16i) (Method B)

To a solution of the desired $\alpha$-amino amide hydrochloride (1.2 equiv), triethylamine (TEA, 1.5 equiv) in anhydrous methanol was added, and then desired biphenyl aldehyde was added at room temperature for 3 h. The reaction mixture was concentrated *in vacuo* and dissolved in EtOAc and washed with brine. The organic layer was dried over $Na_2SO_4$ and then, sodium cyanoborohydride (4–6 equiv) was added to the reaction mixture at 0 ˚C, and then the mixture was stirred at room temperature for 18 h.

The reaction mixture was concentrated *in vacuo* and then the residue was dissolved in EtOAc (150 mL) and washed with brine (2 × 150 mL). The organic layer was dried with anhydrous $Na_2SO_4$ and concentrated *in vacuo*. The residue was purified by column chromatography on Silica gel.

**[1,1'-Biphenyl]-4-carbaldehyde (2a)**

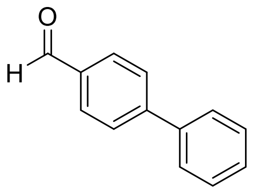


Using method A, 4-bromobenzenealdehyde (**1**) (3.27 g, 20.84 mmol), (4-formylphenyl)boronic acid (4.00 g, 26.67 mmol), tetrakis(triphenylphosphine)palladium(0) (0.96 g, 0.83 mmol) and $Na_2CO_3$ (10.74 g, 101.3 mmol) in toluene/$H_2O$ (175 mL/25.2 mL) gave **2a** as a white solid (3.03 g, 80%); $R_f$ = 0.30 (n-Hexane/EtOAc 10/1); mp 41–44 ˚C; $^1$H NMR (300 MHz, DMSO–d$_6$) $\delta$ 10.06 (s, C(O)**H**), 8.00 (d, *J* = 8.2 Hz, 2Ar**H**), 7.91 (d, *J* = 8.2 Hz, 2Ar**H**), 7.77 (d, *J* = 7.3 Hz, 2Ar**H**), 7.41–7.56 (m, 3Ar**H**); $^{13}$C NMR (75 MHz, DMSO–d$_6$) $\delta$ 193.1 (**C**(O)H), 146.3, 139.3, 135.5, 130.6, 129.6,129.0, 127.8, 127.6 (Ar**C**).

**2'-Fluoro-[1,1'-biphenyl]-4-carbaldehyde (2b)**

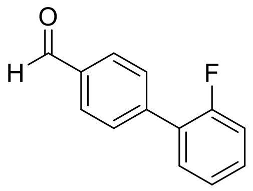


Using Method A, 4-bromobenzaldehyde (**1**) (1.00 g, 5.40 mmol), 2-fluorophenylboronic acid (0.97 g, 6.92 mmol), tetrakis(triphenylphosphine)palladium(0) (0.25 g, 0.216 mmol) and $Na_2CO_3$ (2.78 g, 26.3 mmol) in toluene/$H_2O$ (50 mL/7.2 mL) gave **2b** as a white solid (0.26 g, 24%); $R_f$ = 0.65 (n-Hexane/EtOAc 9/1); mp 43–45 °C; $^1$H NMR (300 MHz, DMSO–*d*$_6$) $\delta$ 10.07 (s, C(O)**H**), 7.98-8.06 (m, 2Ar**H**), 7.79 (dd, *J* = 1.6, 8.2 Hz, 2Ar**H**), 7.62 (td, *J* = 1.7, 7.9 Hz, 1Ar**H**), 7.49–7.55 (m, 1Ar**H**), 7.44–7.49 (m, 2Ar**H**); $^{13}$C NMR (100 MHz, $CDCl_3$) $\delta$ 191.9

(**C**(O)H), 159. 8 (d, $J_{C\text{–}F}$ = 247.7 Hz), 142.0, 135.4, 130.7 (d, $J_{C\text{–}F}$ = 14.4 Hz), 130.2 (d, $J_{C\text{–}F}$ = 8.3 Hz), 129.8, 129.7 (d, $J_{C\text{–}F}$ = 7.5 Hz), 127.8 (q, $J_{C\text{–}F}$ = 13.1 Hz), 124.6, 116.3 (d, $J_{C\text{–}F}$ = 20.8 Hz) (Ar**C**).

**3'-Fluoro-[1,1'-biphenyl]-4-carbaldehyde (2c)**

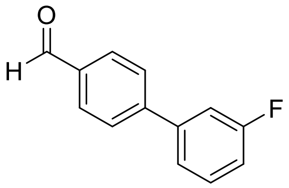


Using Method A, 4-bromobenzaldehyde (**1**) (2.98 g, 16.09 mmol), 3-fluorophenylboronic acid (3.91 g, 20.60 mmol), tetrakis(triphenylphosphine)palladium(0) (0.74 g, 0.64 mmol) and $Na_2CO_3$ (8.29 g, 78.2 mmol) in toluene/$H_2O$ (148.9 mL/21.4 mL) gave **2c** as a white solid (2.64 g, 82%); $R_f$ = 0.30 (n-Hexane/EtOAc 15/1); mp 33–35 °C; $^1$H NMR (400 MHz, DMSO–$d_6$) $\delta$ 10.07 (s, C(O)**H**), 8.01 (d, $J$ = 8.3 Hz, 2Ar**H**), 7.96 (d, $J$ = 8.3 Hz, 2Ar**H**), 7.63–7.67 (m, 2Ar**H**), 7.54–7.59 (m, 1Ar**H**), 7.27–7.31 (m, 1Ar**H**); $^{13}$C NMR (75 MHz, DMSO–$d_6$) 193.0 (**C**(O)H), 163.1 (d, $J_{C\text{–}F}$ = 242.4 Hz, **C**F), 144.8 (d, $J_{C\text{–}F}$ = 2.2 Hz), 141.6 (d, $J_{C\text{–}F}$ = 7.9 Hz), 135.9, 131.4 (d, $J_{C\text{–}F}$ = 8.4 Hz), 130.5, 127.9, 123.6 (d, $J_{C\text{–}F}$ = 2.6 Hz), 115.7 (d, $J_{C\text{–}F}$ = 20.9 Hz), 114.3 (d, $J_{C\text{–}F}$ = 22.3 Hz) (Ar**C**).

**4'-Fluoro-[1,1'-biphenyl]-4-carbaldehyde (2d)**

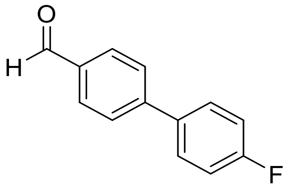


Using Method A, 4-bromobenzaldehyde (**1**) (1.50 g, 8.11 mmol), 4-fluorophenylboronic acid (1.45 g, 10.38 mmol), tetrakis(triphenylphosphine)palladium(0) (0.75 g, 0.65 mmol) and $Na_2CO_3$ (4.18 g, 39.40 mmol) in toluene/$H_2O$ (75 mL/10.8 mL) gave **2d** as a white soild (1.51 g, 93%); $R_f$ = 0.25 (n-Hexane/EtOAc 10/1); mp 79–80 °C; $^1$H NMR (400 MHz, DMSO–$d_6$) $\delta$ 10.05 (s, C(O)**H**), 7.99 (d, $J$ = 8.1 Hz, 2Ar**H**), 7.90 (d, $J$ = 8.1 Hz, 2Ar**H**), 7.84 (d, $J$ = 5.7

Hz, 2Ar**H**), 7.81 (d, $J$ = 5.7 Hz, 2Ar**H**), 7.32–7.37 (m, 2Ar**H**); $^{13}$C NMR (75 MHz, $CDCl_3$) $\delta$ 191.8 (**C**(O)H), 163.2 (d, $J_{C\text{–}F}$ = 246.9 Hz, **C**F), 146.1, 135.9 (d, $J_{C\text{–}F}$ = 3.3 Hz), 135.2, 130.3, 129.1 (d, $J_{C\text{–}F}$ = 8.2 Hz), 127.5, 116.0 (d, $J_{C\text{–}F}$ = 86.1 Hz) (Ar**C**).

**2'-Chloro-[1,1'-biphenyl]-4-carbaldehyde (2e)**

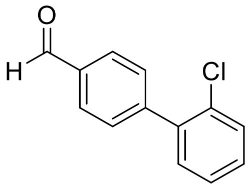


Using method A, , 4-bromobenzaldehyde (**1**) (3.00 g, 16.21 mmol), 2-chlorophenylboronic acid (3.25 g, 20.75 mmol), tetrakis(triphenylphosphine)palladium(0) (0.75 g, 0.65 mmol) and $Na_2CO_3$ (8.35 g, 78.8 mmol) in toluene/$H_2O$ (150 mL/21.6 mL) gave **2e** as a white soild (0.39 g, 11%); $R_f$ = 0.20 (n-Hexane/EtOAc 5/1); mp 71–72 °C; $^1$H NMR (300 MHz, DMSO–$d_6$) $\delta$ 10.08 (s, C(O)**H**), 8.00 (d, $J$ = 7.9 Hz, 2Ar**H**), 7.67 (d, $J$ = 7.9 Hz, 2Ar**H**), 7.58–7.64 (m, 1Ar**H**), 7.44–7.52 (m, 3Ar**H**); $^{13}$C NMR (75 MHz, DMSO–$d_6$) $\delta$ 193.1 (**C**(O)H), 145.0, 139.2, 135.8, 131.6, 130.6, 130.5, 1300.4, 129.8, 128.1 (Ar**C**).

**3'-Chloro-[1,1'-biphenyl]-4-carbaldehyde (2f)**

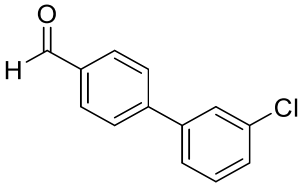


Using Method A, 4-bromobenzaldehyde (**1**) (0.50 g, 2.7 mmol), 3-chlorophenylboronic acid (0.52 g, 3.24 mmol), tetrakis(triphenylphosphine)palladium(0) (0.12 g, 0.11 mmol) and $Na_2CO_3$ (1.39 g, 13.13 mmol) in toluene/$H_2O$ (25 mL/3.6 mL) gave **2f** as a white soild (0.49 g, 84%); $R_f$ = 0.25 (n-Hexane/EtOAc 9/1); mp 49–51 °C; $^1$H NMR (400 MHz, DMSO–$d_6$) $\delta$ 10.07 (s, C(O)**H**), 8.02 (d, $J$ = 8.2 Hz, 2Ar**H**), 7.96 (d, $J$ = 8.2 Hz, 2Ar**H**), 7.85 (s, 1Ar**H**), 7.75 (d, $J$ = 7.3 Hz, 1Ar**H**), 7.57–7.50 (m, 2Ar**H**); $^{13}$C NMR (400 MHz, DMSO–$d_6$) $\delta$ 193.1 (**C**(O)H), 144.6, 141.4, 136.0, 134.4, 131.3, 131.3, 130.6, 128.8, 128.0, 127.4, 127.3, 126.3 (Ar**C**).

**4'-Chloro-[1,1'-biphenyl]-4-carbaldehyde (2g)**

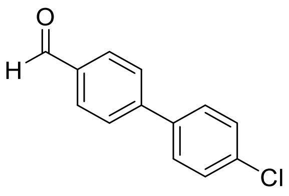


Using Method A, 4-bromobenzaldehyde (**1**) (3.00 g, 16.21 mmol), 4-chlorophenylboronic acid (3.25 g, 20.75 mmol), tetrakis(triphenylphosphine)palladium(0) (0.75 g, 0.65 mmol) and $Na_2CO_3$ (8.35 g, 78.8 mmol) in toluene/$H_2O$ (150 mL/21.6 mL) gave **2g** as a white soild (2.73 g, 78%); $R_f$ = 0.35 (n-Hexane/EtOAc 10/1); mp 142–145 °C; $^1$H NMR (300 MHz, DMSO–$d_6$) $\delta$ 10.06 (s, C(O)**H**), 8.01 (d, $J$ = 8.2 Hz, 2Ar**H**), 7.92 (d, $J$ = 8.2 Hz, 2Ar**H**), 7.81 (d, $J$ = 8.5 Hz, 2Ar**H**), 7.56 (d, $J$ = 8.5 Hz, 2Ar**H**); $^{13}$C NMR (75 MHz, $CDCl_3$) $\delta$ 191.7 (**C**(O)H), 145.8, 138.2, 135.5, 134.8, 130.3, 129.2, 129.1, 128.6, 128.3, 127.9, 127.5 (Ar**C**).

**2'-(Trifluoromethyl)-[1,1'-biphenyl]-4-carbaldehyde (2h)**

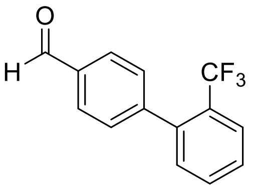


Using Method A, 4-bromobenzaldehyde (**1**) (3.00 g, 16.22 mmol), 2-(trifluoromethyl) phenylboronic acid (3.94 g, 20.76 mmol), tetrakis(triphenylphosphine)palladium(0) (0.76 g, 0.64 mmol) and $Na_2CO_3$ (8.36 g, 78.8 mmol) in toluene/$H_2O$ (150 mL/21.6 mL) gave **2h** as a white solid (0.40 g, 10%); $R_f$ = 0.25 (n-Hexane/EtOAc 15/1); mp 77–80 °C; $^1$H NMR (300 MHz, $CDCl_3$) $\delta$ 10.08 (s, C(O)**H**), 7.93 (d, $J$ = 8.3 Hz, 2Ar**H**), 7.78 (d, $J$ = 7.9 Hz, 2Ar**H**), 7.46–7.65 (m, 4Ar**H**), 7.33 (d, $J$ = 7.32 Hz, 1Ar**H**); $^{13}$C NMR (75 MHz, DMSO–$d_6$) $\delta$ 193.3 (**C**(O)H), 145.7, 139.9, 136.0, 132.9, 132.2, 130.1, 129.5, 129.1, 127.2 (q, $J_{C–F}$ = 29.4 Hz), 126.7 (q, $J_{C–F}$ = 5.2 Hz), 124.5 (q, $J_{C–F}$ = 272.2 Hz) (Ar**C**).

**3'-(Trifluoromethyl)-[1,1'-biphenyl]-4-carbaldehyde (2i)**

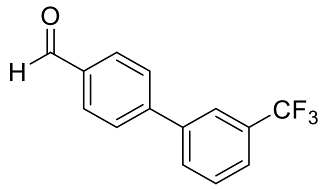


Using Method A, 4-bromobenzaldehyde (**1**) (0.50 g, 2.70 mmol), 3-(trifluoromethyl) phenylboronic acid (0.67 g, 3.51 mmol), tetrakis(triphenylphosphine)palladium(0) (0.13 g, 0.11 mmol) and $Na_2CO_3$ (1.43 g, 13.51 mmol) in toluene/$H_2O$ (25 mL/3.6 mL) gave **2i** as a colorless oil (0.61 g, 97%); $R_f$ = 0.20 (n-Hexane/EtOAc 15/1); $^1$H NMR (400 MHz, DMSO–$d_6$) $\delta$ 10.10 (s, C(O)**H**), 8.00–8.10 (m, 6Ar**H**), 7.73–7.82 (m, 2Ar**H**); $^{13}$C NMR (75 MHz, $CDCl_3$) $\delta$ 191.8 (**C**(O)H), 145.6, 130.6, 135.8, 131.5 (q, $J_{C–F}$ = 32.2 Hz), 130.7, 130.4, 129.6, 127.9, 125.1 (q, $J_{C–F}$ = 3.8 Hz), 124.2 (q, $J_{C–F}$ = 3.7 Hz), 124.0 (q, $J_{C–F}$ = 270.6 Hz) (Ar**C**).

**4'-(Trifluoromethyl)-[1,1'-biphenyl]-4-carbaldehyde (2j)**

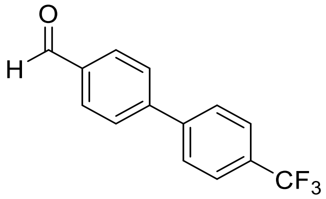


Using Method A, 4-bromobenzaldehyde (**1**) (1.50 g, 8.11 mmol), 4’-(trifluoromethyl)phenylboronic acid (1.97 g, 10.38 mmol), tetrakis(triphenylphosphine)palladium(0) (0.38 g, 0.32 mmol) and $Na_2CO_3$ (4.18 g, 39.4 mmol) in toluene/$H_2O$ (75 mL/10.8 mL) gave **2j** as a white solid(1.17 g, 58%); $R_f$ = 0.80 (EtOAc); mp 73–74 ˚C; $^1$H NMR (300 MHz, $CDCl_3$) $\delta$ 10.09 (s, C(O)**H**), 7.97–8.01 (m, 2Ar**H**), 7.74–7.77 (m, 6Ar**H**); $^{13}$C NMR (75 MHz, $CDCl_3$) $\delta$ 193.2 (**C**(O)H), 144.6, 143.3, 136.2, 130.7, 129.2 (q, $J_{C–F}$ = 31.7 Hz), 128.4, 128.3, 126.4 (q, $J_{C–F}$ = 3.8 Hz), 124.6 (q, $J_{C–F}$ = 264.0 Hz) (Ar**C**).

**3'-(Trifluoromethoxy)-[1,1'-biphenyl]-4-carbaldehyde (2k)**

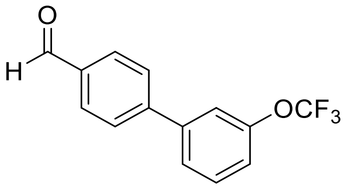


Using Method A, 4-bromobenzaldehyde (**1**) (2.00 g, 10.8 mmol), 3-(trifluoromethoxy) phenylboronic acid (2.85 g, 13.8 mmol), tetrakis(triphenylphosphine)palladium(0) (0.5 g, 0.43 mmol) and $Na_2CO_3$ (5.57 g, 52.5 mmol) in toluene/$H_2O$ (100 mL/14.4 mL) gave **2k** as a colorless oil (2.15 g, 75%); $R_f$ = 0.35 (n-Hexane/EtOAc 10/1); $^1$H NMR (300 MHz, DMSO–$d_6$) $\delta$ 10.08 (s, C(O)**H**), 8.02 (dd, $J$ = 1.9, 6.6 Hz, 2Ar**H**), 7.97 (dd, $J$ = 1.9, 6.6 Hz, 2Ar**H**), 7.81–7.86 (m, 1Ar**H**), 7.76 (s, 1Ar**H**), 7.66 (t, $J$ = 8.0 Hz, 1Ar**H**), 7.42–7.50 (m, 1Ar**H**); $^{13}$C NMR (75 MHz, $CDCl_3$) $\delta$ 191.8 (**C**(O)H), 149.8 (**C**$OCF_3$), 145.5, 141.9, 135.8, 130.4, 130.3, 128.7, 125.7, 120.7, 120.5 (q, $J_{C\text{–}F}$ = 256.0 Hz), 120.0 (Ar**C**).

**4'-(Trifluoromethoxy)-[1,1'-biphenyl]-4-carbaldehyde (2l)**

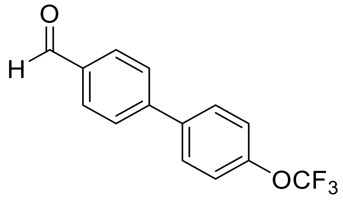


Using Method A, 4-bromobenzaldehyde (**1**) (0.50 g, 2.70 mmol), 4’-(trifluoromethoxy)phenylboronic acid (0.71 g, 3.46 mmol), tetrakis(triphenylphosphine)palladium(0) (0.13 g, 0.108 mmol) and $Na_2CO_3$ (1.39 g, 13.1 mmol) in toluene/$H_2O$ (25 mL/3.6 mL) gave **2l** as a white solid (0.71 g, 98%); $R_f$ = 0.65 (n-Hexane/EtOAc 9/1); mp 31–33 ˚C; $^1$H NMR (300 MHz, DMSO–$d_6$) $\delta$ 10.07 (s, C(O)**H**), 7.96–8.05 (m, 2Ar**H**), 7.88–7.95 (m, 4Ar**H**), 7.51 (d, $J$ = 8.0 Hz, 2Ar**H**); $^{13}$C NMR (75 MHz, $CDCl_3$) $\delta$ 191.7 (**C**(O)H), 149.5266, 145.7, 138.4, 135.5, 130.3, 128.8, 127.7, 121.4, 120.5 (q, $J_{C\text{–}F}$ = 256.1 Hz) (Ar**C**).

**3'-Methoxy-[1,1'-biphenyl]-4-carbaldehyde (2m)**

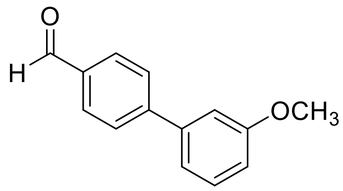


Using Method A, 4-bromobenzaldehyde (**1**) (0.50 g, 2.70 mmol), 3-methoxyphenylboronic acid (0.53 g, 3.46 mmol), tetrakis(triphenylphosphine)palladium(0) (0.13 mg, 0.108 mmol) and $Na_2CO_3$ (1.39 g, 13.1 mmol) in toluene/$H_2O$ (25 mL/3.6 mL) gave **2m** as a colorless oil (0.57 g, 99%); $R_f$ = 0.25 (n-Hexane/EtOAc 10/1); $^1$H NMR (300 MHz, DMSO–$d_6$) $\delta$ 10.06 (s, C(O)**H**), 7.99 (d, $J$ = 8.2 Hz, 2Ar**H**), 7.92 (d, $J$ = 8.2 Hz, 2Ar**H**), 7.43 (t, $J$ = 7.8 Hz, 1Ar**H**), 7.25–7.34 (m, 2Ar**H**), 6.98–7.05 (m, 1Ar**H**), 3.84 (s, OC**H**3); $^{13}$C NMR (75 MHz, $CDCl_3$) $\delta$ 191.8 (**C**(O)H), 160.1, 147.0, 141.2, 135.3, 130.2, 130.0, 127.7, 119.8, 113.8, 113.2 (Ar**C**), 55.3 (O**C**H3).

**4'-Methoxy-[1,1'-biphenyl]-4-carbaldehyde (2n)**

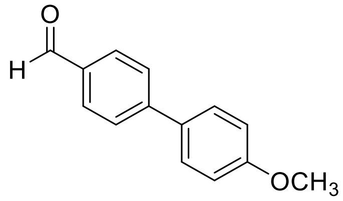


Using Method A, 4-bromobenzaldehyde (**1**) (2.00 g, 10.81 mmol), 3-(trifluoromethyl) phenylboronic acid (2.10 g, 13.84 mmol), tetrakis(triphenylphosphine)palladium(0) (0.5 g, 0.43mmol) and $Na_2CO_3$ (5.57 g, 52.5 mmol) in toluene/$H_2O$ (100 mL/14.4 mL) gave **2n** as a white solid(1.26 g, 55%); $R_f$ = 0.20 (n-Hexane/EtOAc 10/1); mp 105–106 ˚C; $^1$H NMR (300 MHz, DMSO–d6) $\delta$ 10.03 (s, C(O)**H**), 7.96 (d, $J$ = 8.1 Hz, 2Ar**H**), 7.88 (d, $J$ = 8.3 Hz, 2Ar**H**), 7.75 (d, $J$ = 8.8 Hz, 2Ar**H**), 7.08 (d, $J$ = 8.8 Hz, 2Ar**H**), 3.83 (s, OC**H3**); $^{13}$C NMR (75 MHz, DMSO–d6) $\delta$ 193.0 (**C**(O)), 160.3, 146.0, 135.0, 131.4, 130.6, 128.8, 127.1, 115.1 (Ar**C**), 55.8 (O**C**H3).

**4'-Isobutyl-[1,1'-biphenyl]-4-carbaldehyde (2o)**

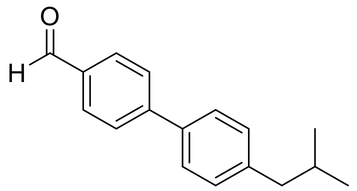

Using Method A, 4-bromobenzaldehyde (**1**) (3.50 g, 18.92 mmol), 4'-isobutylphenylboronic acid (4.31 g, 24.21 mmol), tetrakis(triphenylphosphine)palladium(0) (0.87 g, 0.76 mmol) and $Na_2CO_3$ (9.74 g, 91.9 mmol) in toluene/$H_2O$ (175 mL/25.2 mL) gave **2o** as a white solid (4.16 g, 93%); $R_f$ = 0.30 (n-Hexane/EtOAc 15/1); mp 50–51 ˚C; $^1$H NMR (300 MHz, DMSO–d$_6$) $\delta$ 10.04 (s, C(O)**H**), 7.98 (d, *J* = 8.0 Hz, 2Ar**H**), 7.90 (d, *J* = 8.0 Hz, 2Ar**H**), 7.69 (d, *J* = 7.7 Hz, 2Ar**H**), 7.29 (d, *J* = 7.7 Hz, 2Ar**H**), 2.50 (d, *J* = 6.2 Hz, C**H$_2$**), 1.80–1.98 (m, C**H**), 0.82–0.97 (m, 2C**H$_3$**); $^{13}$C NMR (75 MHz, DMSO–d$_6$) $\delta$ 193.1 (**C**(O)H), 146.3, 142.3, 136.6, 135.3, 130.6, 130.2, 127.5, 127.3 (Ar**C**), 44.7 (C**H$_2$**), 30.0 (**C**H), 22.6 (CH(**C**H$_3$)**$_2$**).

**2-(([1,1'-Biphenyl]-4-ylmethyl)amino)acetamide (10a)**

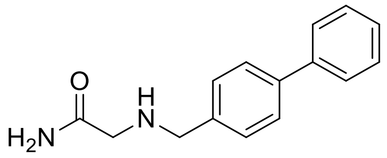

Using method B, glycinamide hydrochloride (**3**) (0.39 g, 3.51 mmol), triethylamine (0.57 mL, 4.11 mmol), **2a** (0.50 g, 2.74 mmol) in MeOH (2.98 mL), and sodium cyanoborohydride (0.73 g, 10.98 mmol) in MeOH (2.74 mL) gave **10a** as a white solid (0.25 g, 38%); $R_f$ = 0.15 (EtOAc); mp 166–167 ˚C; $^1$H NMR (300 MHz, DMSO–*d*$_6$) $\delta$ 7.58–7.63 (m, 4Ar**H**), 7.39–7.50 (m, 4Ar**H**), 7.26–7.39 (m, 1ArH, C(O)N**H**H'), 7.06 (br s, C(O)NH**H'**), 3.71 (s, NHC**H$_2$**Ar), 3.05 (s, C(O)C**H$_2$**NH), 2.60 (br s, N**H**); $^{13}$C NMR (75 MHz, DMSO–*d*$_6$) $\delta$ 173.8 (**C**(O)), 140.6, 140.2, 139.1, 129.4, 129.0, 127.7, 127.0, 126.9 (Ar**C**), 52.8 (C(O)**C**H$_2$NH), 51.8 (NH**C**H$_2$Ar).

**2-(((4'-Fluoro-[1,1'-biphenyl]-4-yl)methyl)amino)acetamide (10d)**

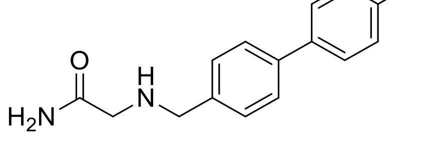

Using method B, glycinamide hydrochloride (**3**) (0.20 g, 1.80 mmol), triethylamine (0.31 mL, 2.25 mmol), **2d** (0.30 g, 1.50 mmol) in MeOH (1.63 mL), and sodium cyanoborohydride (0.40 g, 6.0 mmol) in MeOH (1.5 mL) gave **10d** as a white solid (0.19 g, 49%); $R_f$ = 0.10 (EtOAc); mp 161–162 ˚C; $^1$H NMR (300 MHz, DMSO–$d_6$) $\delta$ 7.65–7.75 (m, 2Ar**H**), 7.63 (d, $J$ = 8.1 Hz, 2Ar**H**), 7.41 (d, $J$ = 8.1 Hz, 2Ar**H**), 7.22–7.34 (m, 2ArH, C(O)N**H**H'), 7.06 (br s, C(O)NH**H'**), 3.71 (s, NHC**H₂**Ar), 3.04 (s, C(O)C**H₂**NH), the remaining peak was not detected and is believed to overlap with $H_2O$ signals; $^{13}$C NMR (75 MHz, DMSO–$d_6$) $\delta$ 173.8 (**C**(O)), 162.2 (d, $J_{C–F}$ = 242.5 Hz), 140.2, 138.0, 137.0 (d, $J_{C–F}$ = 3.0 Hz), 129.0, 128.9 (d, $J_{C–F}$ = 8.4 Hz), 126.9, 116.19 (d, $J_{C–F}$ = 21.1 Hz) (Ar**C**), 52.7 (C(O)**C**$H_2$NH), 51.7 (NH**C**$H_2$Ar).

**2-(((3'-Chloro-[1,1'-biphenyl]-4-yl)methyl)amino)acetamide (10f)**

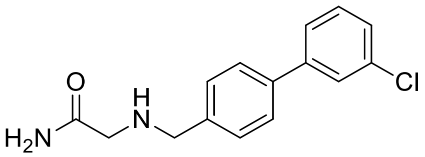


Using method D, glycinamide hydrochloride (**3**) (0.31 g, 2.77 mmol), triethylamine (0.48 mL, 3.46 mmol), **2f** (0.50 g, 2.31 mmol) in MeOH (2.51 mL), and sodium cyanoborohydride (0.89 g, 9.23 mmol) in MeOH (2.31 mL) gave **10f** as a white solid (0.30 g, 47%); $R_f$ = 0.10 (EtOAc); mp 140–141 ˚C; $^1$H NMR (300 MHz, DMSO–$d_6$) $\delta$ 7.59–7.76 (m, 4Ar**H**), 7.37–7.53 (m, 4Ar**H**), 7.33 (br s, C(O)N**H**H'), 7.08 (br s, C(O)NH**H'**), 3.72 (s, NHC**H₂**Ar), 3.06 (s, C(O)C**H₂**NH), 2.70 (br s, N**H**); $^{13}$C NMR (75 MHz, DMSO–$d_6$) $\delta$ 173.8 (**C**(O)), 142.7, 140.9, 137.5, 134.2, 131.2, 129.1, 127.5, 127.1, 126.7, 125.7 (Ar**C**), 52.7 (C(O)**C**$H_2$NH), 51.7 (NH**C**$H_2$Ar).

**2-(((4'-Chloro-[1,1'-biphenyl]-4-yl)methyl)amino)acetamide (10g)**

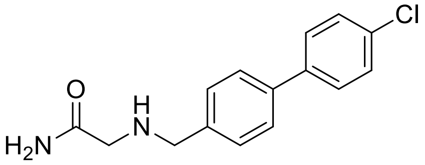


Using method B, glycinamide hydrochloride (**3**) (0.31 g, 2.77 mmol), triethylamine (0.48 mL, 3.46 mmol), **2g** (0.50 g, 2.31 mmol) in MeOH (2.51 mL), and sodium cyanoborohydride (0.89 g, 9.23 mmol) in MeOH (2.31 mL) gave **10g** as a white solid (0.21 g, 34%); $R_f$ = 0.10 (EtOAc);

mp 167–169 ˚C; $^{1}$H NMR (300 MHz, DMSO–$d_6$) $\delta$ 7.69 (d, $J$ = 8.4 Hz, 2Ar**H**), 7.62 (d, $J$ = 8.0 Hz, 2Ar**H**), 7.51 (d, $J$ = 8.4 Hz, 2Ar**H**), 7.43 (d, $J$ = 8.0 Hz, 2Ar**H**), 7.30 (br s, C(O)N**H**H'), 7.06 (br s, C(O)NH**H'**), 3.71 (s, NHC**H**$_2$Ar), 3.03 (s, C(O)C**H**$_2$NH), 2.62 (br s, N**H**); $^{13}$C NMR (75 MHz, DMSO–$d_6$) $\delta$ 173.9 (**C**(O)), 140.6, 139.3, 137.7, 132.6, 129.3, 129.1, 128.8, 126.9 (Ar**C**), 52.7 (C(O)**C**H$_2$NH), 51.7 (NH**C**H$_2$Ar).

**2-(((3'-(Trifluoromethyl)-[1,1'-biphenyl]-4-yl)methyl)amino)acetamide (10i)**

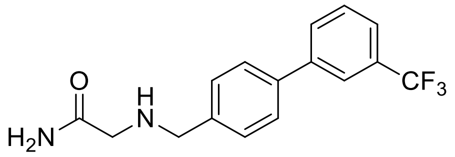


Using method B, glycinamide hydrochloridem (**3**) (0.21 g, 1.92 mmol), triethylamine (0.33 mL, 2.40 mmol), **2i** (0.40 g, 1.60 mmol) in MeOH (1.74 mL), and sodium cyanoborohydride (0.65 g, 6.39 mmol) in MeOH (1.60 mL) gave **10i** as a white solid (0.26 g, 52%); $R_f$ = 0.15 (EtOAc); mp 122–126 ˚C; $^{1}$H NMR (300 MHz, DMSO–$d_6$) $\delta$ 7.91–8.03 (m, 2Ar**H**), 7.64–7.78 (m, 4Ar**H**), 7.46 (d, $J$ = 8.0 Hz, 2Ar**H**), 7.31 (br s, C(O)N**H**H'), 7.06 (br s, C(O)NH**H'**), 3.73 (s, NHC**H**$_2$Ar), 3.06 (s, C(O)C**H**$_2$NH), 2.64 (br s, NH); $^{13}$C NMR (75 MHz, CDCl$_3$) $\delta$ 173.8 (**C**(O)), 141.6, 141.1, 137.4, 131.1, 130.5, 130.2 (q, $J_{C–F}$ = 31.4 Hz), 129.2, 127.2, 124.7 (q, $J_{C–F}$ = 270.8 Hz), 124.3 (q, $J_{C–F}$ = 3.7 Hz), 123.4 (q, $J_{C–F}$ = 3.8 Hz) (Ar**C**), 52.7 (C(O)**C**H$_2$NH), 51.7 (NH**C**H$_2$Ar).

**2-(((4'-(Trifluoromethyl)-[1,1'-biphenyl]-4-yl)methyl)amino)acetamide (10j)**

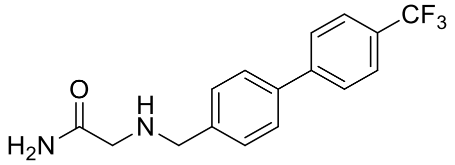


Using method B, glycinamide hydrochloride (**3**) (0.27 g, 2.40 mmol), triethylamine (0.42 mL, 3.0 mmol), **2j** (0.50 g, 2.0 mmol) in MeOH (2.17 mL), and sodium cyanoborohydride (0.49 g, 7.64 mmol) in MeOH (2.0 mL) gave **10j** as a white solid (0.28 g, 45%); $R_f$ = 0.10 (EtOAc); mp 172–173 ˚C; $^{1}$H NMR (300 MHz, DMSO–$d_6$) $\delta$ 7.89 (d, $J$ = 8.2 Hz, 2Ar**H**), 7.80 (d, $J$ = 8.2 Hz, 2Ar**H**), 7.69 (d, $J$ = 8.1 Hz, 2Ar**H**), 7.47 (d, $J$ = 8.1 Hz, 2Ar**H**), 7.31 (br s, C(O)N**H**H'), 7.06 (br s, C(O)NH**H'**), 3.73 (s, NHC**H**$_2$Ar), 3.05 (s, C(O)C**H**$_2$NH), 2.64 (br s, NH); $^{13}$C NMR (75

MHz, DMSO–$d_6$) $\delta$ 173.8 (**C**(O)), 144.5, 141.4, 137.4, 130.2, 128.1 (q, $J_{C–F}$ = 31.7 Hz), 127.7, 127.5, 126.2 (q, $J_{C–F}$ = 3.7 Hz), 124.8 (q, $J_{C–F}$ = 270.1 Hz) (Ar**C**), 52.7 (C(O)**C**$H_2$NH), 51.8 (NH**C**$H_2$Ar).

**2-(((3'-(Trifluoromethoxy)-[1,1'-biphenyl]-4-yl)methyl)amino)acetamide (10k)**

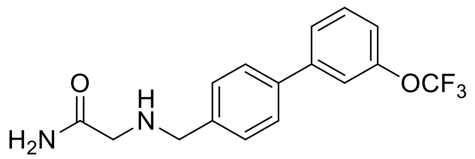

Using method B, glycinamide hydrochloride (**3**) (0.15 g, 1.40 mmol), triethylamine (0.24 mL, 1.70 mmol), **2k** (0.30 g, 1.10 mmol) in MeOH (1.22 mL), and sodium cyanoborohydride (0.30 g, 4.51 mmol) in MeOH (1.12 mL) gave **1k** as a white solid (0.03 g, 8%); $R_f$ = 0.13 (EtOAc); mp 124–127 ˚C; $^1$H NMR (300 MHz, DMSO–$d_6$) $\delta$ 7.55–7.77 (m, 5Ar**H**), 7.45 (d, $J$ = 7.9 Hz, 2Ar**H**), 7.35 (d, $J$ = 8.4 Hz, 2Ar**H**), 7.30 (br s, C(O)N**H**H'), 7.05 (br s, C(O)NH**H'**), 3.72 (s, NHC**H**$_2$Ar), 3.05 (s, C(O)C**H**$_2$NH), 2.85 (br s, N**H**); $^{13}$C NMR (75 MHz, DMSO–$d_6$) $\delta$ 173.8 (**C**(O)), 149.4 (**C**OC$F_3$), 142.9, 141.1, 137.3, 131.3, 129.1, 127.1, 126.1, 120.6 (q, $J_{C–F}$ = 254.5 Hz), 120.0, 119.5 (Ar**C**), 52.6 (C(O)**C**$H_2$NH), 51.7 (NH**C**$H_2$Ar).

**2-(((4'-(Trifluoromethoxy)-[1,1'-biphenyl]-4-yl)methyl)amino)acetamide (10l)**

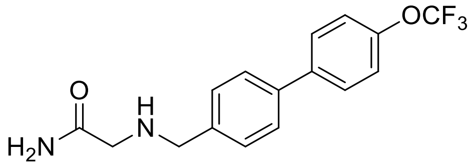

Using method B, glycinamide hydrochloride (**3**) (0.25 g, 2.26 mmol), triethylamine (0.39 mL, 2.83 mmol), **2l** (0.50 g, 1.88 mmol) in MeOH (2.04 mL), and sodium cyanoborohydride (0.76 g, 7.51 mmol) in MeOH (1.88 mL) gave **10l** as a white solid (0.33 g, 54%); $R_f$ = 0.10 (EtOAc); mp 170–171 ˚C; $^1$H NMR (300 MHz, DMSO–$d_6$) $\delta$ 7.78 (d, $J$ = 8.7 Hz, 2Ar**H**), 7.63 (d, $J$ = 8.1 Hz, 2Ar**H**), 7.39–7.49 (m, 4Ar**H**), 7.31 (br s, C(O)N**H**H'), 7.06 (br s, C(O)NH**H'**), 3.72 (s, NHC**H**$_2$Ar), 3.05 (s, C(O)C**H**$_2$NH), 2.63 (br s, N**H**); $^{13}$C NMR (75 MHz, DMSO–$d_6$) $\delta$ 173.8 (**C**(O)), 148.2 (**C**OC$F_3$), 140.7, 139.6, 137.6, 129.1, 128.9, 127.1, 121.9, 120.6 (q, $J_{C–F}$ = 254.5 Hz) (Ar**C**), 52.6 (C(O)**C**$H_2$NH), 51.7 (NH**C**$H_2$Ar).

**2-(((4'-Methoxy-[1,1'-biphenyl]-4-yl)methyl)amino)acetamide (10n)**

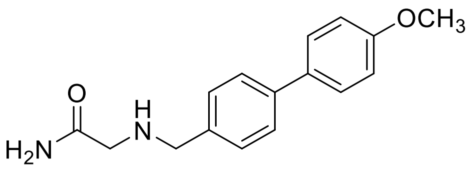

Using method B, glycinamide hydrochloride (**3**) (0.25 g, 2.26 mmol), triethylamine (0.39 mL, 2.83 mmol), **2n** (0.40 g, 1.80 mmol) in MeOH (2.05 mL), and sodium cyanoborohydride (0.76 g, 7.54 mmol) in MeOH (1.88 mL) gave **10n** as a light yellow solid (0.12 g, 23%); $R_f$ = 0.05 (EtOAc); mp 162–165 ˚C; $^{1}$H NMR (300 MHz, DMSO–$d_6$) $\delta$ 7.52–7.63 (m, 4Ar**H**), 7.38 (d, $J$ = 8.1 Hz, 2Ar**H**), 7.30 (br s, C(O)N**H**H'), 6.97–7.08 (m, 2ArH, C(O)N**H**H'), 3.79 (s, OC**H**$_3$), 3.69 (s, NHC**H**$_2$Ar) 3.04 (s, C(O)C**H**$_2$NH), 2.65 (br s, N**H**); $^{13}$C NMR (75 MHz, DMSO–$d_6$) $\delta$ 173.8 (**C**(O)), 159.2 (**C**OCH$_3$), 139.4, 138.7, 132.9, 130.0, 128.0, 126.4, 114.8 (Ar**C**), 55.6 (O**C**H$_3$), 52.7 (C(O)**C**H$_2$NH), 51.7 (NH**C**H$_2$Ar).

**(*S*)-2-(([1,1'-Biphenyl]-4-ylmethyl)amino)propanamide (11a)**

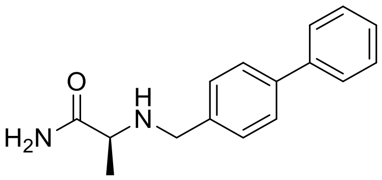

Using method B, L-alaninamide hydrochloride (**4**) (0.44 g, 3.51 mmol), triethylamine (0.57 mL, 2.98 mmol) **2a** (0.50 g, 2.74 mmol) in MeOH (2.98 mL) and sodium cyanoborohydride (0.69 g, 10.98 mmol) in MeOH (2.74 mL) gave **11a** as a white solid (0.28 g, 41%); $R_f$ = 0.10 (EtOAc); mp 157–159 ˚C; $^{1}$H NMR (300 MHz, DMSO–$d_6$) $\delta$ 7.57–7.68 (m, 4Ar**H**), 7.31–7.50 (m, 5ArH, C(O)N**H**H'), 6.99 (br s, C(O)NH**H'**), 3.58 and 3.73 (AB$_q$, $J$ = 13.6 Hz, C**H**$_2$), 3.04 (q, $J$ = 6.8 Hz, C**H**), 2.42 (br s, N**H**), 1.15 (d, $J$ = 6.8 Hz, C**H**$_3$); $^{13}$C NMR (75 MHz, DMSO–$d_6$) $\delta$ 177.4 (**C**(O)), 140.6, 140.3, 139.0, 129.4, 129.0, 127.7, 127.0, 126.9 (Ar**C**), 57.0 (**C**H), 51.2 (**C**H$_2$), 19.8 (**C**H$_3$).

**(*S*)-2-(((2'-Fluoro-[1,1'-biphenyl]-4-yl)methyl)amino)propanamide (11b)**

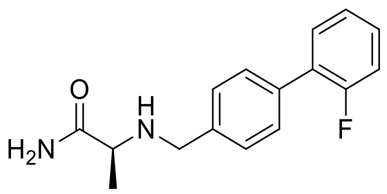

Using method B, L-alaninamide hydrochloride (**4**) (0.19 g, 1.52 mmol), triethylamine (0.21 mL, 1.52 mmol) **2b** (0.25 g, 1.27 mmol) and sodium cyanoborohydride (0.13 g, 1.90 mmol) in MeOH (1.3 mL) gave **11b** as a white solid (0.21 g, 62%); $R_f$ = 0.10 (EtOAc); mp 133–136 °C; $^1$H NMR (300 MHz, DMSO–$d_6$) $\delta$ 7.26–7.58 (m, 8Ar**H,** C(O)N**H**H'), 7.01 (br s, C(O)NH**H'**), 3.59 and 3.74 (AB$_q$, $J$ = 13.7 Hz, C**H**$_2$), 3.03 (q, $J$ = 6.8 Hz, C**H**), 2.43 (br s, N**H**), 1.15 (d, $J$ = 6.8 Hz, C**H**$_3$); $^{13}$C NMR (75 MHz, CDCl$_3$) $\delta$ 178.1 (**C**(O)), 159.8 (d, $J_{C–F}$ = 246.1 Hz), 139.0, 134.9, 130.7 (d, $J_{C–F}$ = 3.5 Hz), 129.2 (d, $J_{C–F}$ = 2.8 Hz), 128.8 (d, $J_{C–F}$ = 8.2 Hz), 128.1, 124.4 (d, $J_{C–F}$ = 3.7 Hz), 116.1 (d, $J_{C–F}$ = 22.6 Hz) (Ar**C**), 57.8 (**C**H), 52.2 (**C**H$_2$), 19.7 (**C**H$_3$).

**(*S*)-2-(((3'-Fluoro-[1,1'-biphenyl]-4-yl)methyl)amino)propanamide (11c)**

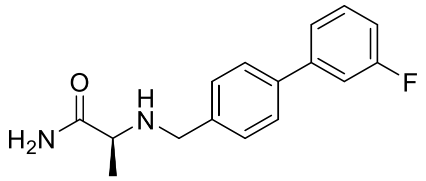

Using method B, L-alaninamide hydrochloride (**4**) (0.58 g, 4.63 mmol), triethylamine (0.81 mL, 5.78 mmol), **2c** (0.77 g, 3.85 mmol) in MeOH (4.19 mL), and sodium cyanoborohydride (0.38 g, 5.78 mmol) in MeOH (3.85 mL) gave **11c** as a white solid (0.48 g, 45%); $R_f$ = 0.20 (EtOAc); mp 128–130 °C; $^1$H NMR (300 MHz, DMSO–$d_6$) $\delta$ 7.65 (d, $J$ = 8.12 Hz, 2Ar**H**), 7.48–7.52 (m, 3Ar**H**), 7.43 (d, $J$ = 8.2 Hz, 2Ar**H**), 7.34 (br s, C(O)N**H**H'), 7.14–7.21(m, 1Ar**H**), 6.99 (br s, C(O)NH**H'**), 3.59 and 3.73 (AB$_q$, $J$ = 13.7 Hz, C**H**$_2$), 3.03 (q, $J$ = 6.9 Hz, C**H**), 2.43 (br s, N**H**), 1.15 (d, $J$ = 6.87 Hz, C**H**$_3$); $^{13}$C NMR (75 MHz, CDCl$_3$) $\delta$ 178.1 (**C**(O)), 163.2 (d, $J_{C–F}$ = 244.0 Hz), 143.1 (d, $J_{C–F}$ = 7.6 Hz), 139.3, 139.0 (d, $J_{C–F}$ = 2.2 Hz), 130.3 (d, $J_{C–F}$ = 8.3 Hz), 128.6, 127.3, 122.7 (d, $J_{C–F}$ = 2.7 Hz), 114.1 (d, $J_{C–F}$ = 21.0 Hz), 113.9 (d, $J_{C–F}$ = 21.9 Hz) (Ar**C**), 57.7 (**C**H), 52.2 (**C**H$_2$), 19.7 (**C**H$_3$).

**(*S*)-2-(((4'-Fluoro-[1,1'-biphenyl]-4-yl)methyl)amino)propanamide (11d)**

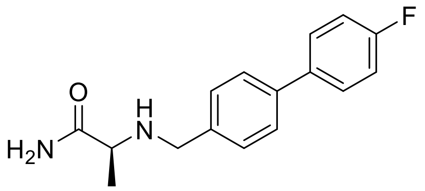

Using method B, L-alaninamide hydrochloride (**4**) (0.37 g, 3.0 mmol), triethylamine (1.04 mL, 7.49 mmol), **2d** (0.50 g, 2.50 mmol) in MeOH (2.71 mL), and sodium cyanoborohydride (0.28 g, 3.75 mmol) in MeOH (2.50 mL) gave **11d** as a white solid (0.35 g, 51%); $R_f$ = 0.20 (EtOAc); mp 129–133 °C; $^1$H NMR (300 MHz, DMSO–$d_6$) $\delta$ 7.65–7.72 (m, 2Ar**H**), 7.58 (d, $J$ = 8.2 Hz, 2Ar**H**), 7.35 (br s, C(O)N**H**H'), 7.24–7.31(m, 2Ar**H**), 6.99 (br s, C(O)NH**H'**), 3.58 and 3.72 (AB$_q$, $J$ = 13.7 Hz, C**H**$_2$), 3.03 (q, $J$ = 6.9 Hz, C**H** 1.15 (d, $J$ = 6.9 Hz, C**H**$_3$), the remaining peak was not detected and is believed to overlap with $H_2O$ signals; $^{13}$C NMR (100 MHz, DMSO–d$_6$) $\delta$ 177.4 (**C**(O)), 162.2 (d, $J_{C–F}$ = 181.9 Hz), 140.4, 138.0, 137.0 (d, $J_{C–F}$ = 2.3 Hz), 129.0, 126.9 (d, $J_{C–F}$ = 5.2 Hz), 116.2 (Ar**C**), 56.9 (**C**H), 51.1 (**C**H$_2$), 19.8 (**C**H$_3$), the remaining aromatic peak was not detected.

**(*S*)-2-(((2'-Chloro-[1,1'-biphenyl]-4-yl)methyl)amino)propanamide (11e)**

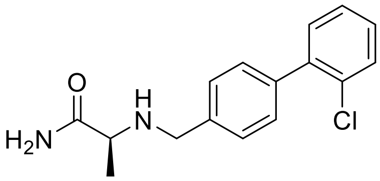

Using method B, L-alaninamide hydrochloride (**4**) (0.16 g, 1.44 mmol), triethylamine (0.25 mL, 3.53 mmol), **2e** (0.26 g, 1.20 mmol) in MeOH (1.30 mL), and sodium cyanoborohydride (0.32 g, 4.80 mmol) in MeOH (1.20 mL) gave **11e** as a white solid 0.16 g, 47%); R$_f$ = 0.15 (EtOAc); mp 91–94 °C; $^1$H NMR (300 MHz, DMSO–$d_6$) $\delta$ 7.53–7.58 (m, 1Ar**H**), 7.30–7.48 (m, 7ArH, C(O)N**H**H'), 6.98 (br s, C(O)NH**H'**), 3.60 and 3.74 (AB$_q$, $J$ = 13.6 Hz, C**H**$_2$), 3.06 (q, $J$ = 6.8 Hz, C**H**), 2.44 (br s, N**H**), 1.17 (d, $J$ = 6.8 Hz, C**H**$_3$); $^{13}$C NMR (75 MHz, DMSO–$d_6$) $\delta$ 177.4 (**C**(O)), 140.7, 140.2, 137.5, 132.0, 131.8, 130.3, 129.5, 129.4, 128.2, 128.0 (Ar**C**), 57.1 (**C**H), 51.2 (**C**H$_2$), 19.9 (**C**H$_3$).

**(*S*)-2-(((3'-Chloro-[1,1'-biphenyl]-4-yl)methyl)amino)propanamide (11f)**

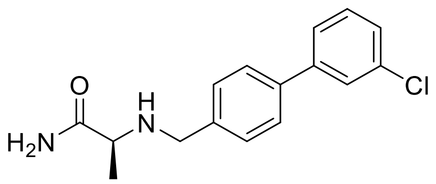

Using method B, L-alaninamide hydrochloride (**4**) (0.21 g, 1.66 mmol), triethylamine (0.23 mL, 1.66 mmol), **2f** (0.30 g, 1.38 mmol), and sodium cyanoborohydride (0.13 g, 2.08 mmol) in MeOH (1.36 mL) gave **11f** as a white solid (0.22 mg, 54%); $R_f$ = 0.10 (EtOAc); mp 121–123 °C; $^{1}$H NMR (400 MHz, DMSO–$d_6$) $\delta$ 7.70 (s, 1Ar**H**), 7.62–7.65 (m, 3Ar**H**), 7.48 (t, *J* = 7.8 Hz, 1Ar**H**), 7.40–7.48 (m, 3Ar**H**), 7.34 (br s, C(O)N**H**H'), 6.99 (br s, C(O)NH**H'**), 3.59 and 3.74 ($AB_q$, *J* = 13.7 Hz, C$\mathbf{H_2}$), 3.03 (q, *J* = 6.7 Hz, C**H**), 2.44 (br s, N**H**), 1.15 (d, *J* = 6.8 Hz, C$\mathbf{H_3}$); $^{13}$C NMR (75 MHz, DMSO–$d_6$) $\delta$ 177.4 (**C**(O)), 142.7, 141.1, 137.4, 134.2, 131.2, 129.0, 127.5, 127.0, 126.7, 125.7 (Ar**C**), 57.0(**C**H), 51.1(**C**$H_2$), 19.8 (**C**$H_3$).

**(*S*)-2-(((4'-Chloro-[1,1'-biphenyl]-4-yl)methyl)amino)propanamide(11g)**

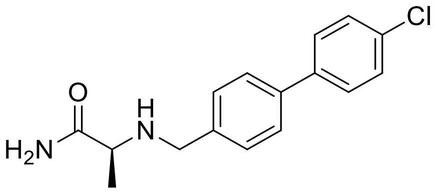

Using method B, L-alaninamide hydrochloride (**4**) (0.35 g, 3.00 mmol), triethylamine (0.97 mL, 6.92 mmol), **2g** (0.50 g, 2.31 mmol) in MeOH (2.51 mL), and sodium cyanoborohydride (0.23 g, 3.46 mmol) in MeOH (2.31 mL) gave **11g** as a white solid (386 mg, 58%); $R_f$ = 0.15 (EtOAc); mp 166–164 °C; $^{1}$H NMR (300 MHz, DMSO–$d_6$) $\delta$ 7.66–7.70 (m, 2Ar**H**), 7.61 (d, *J* = 8.2 Hz, 2Ar**H**), 7.49–7.53 (m, 2Ar**H**), 7.42 (d, *J* = 8.2 Hz, 2Ar**H**), 7.33 (br s, C(O)N**H**H'), 6.99 (br s, C(O)NH**H'**), 3.58 and 3.73 ($AB_q$, *J* = 13.7 Hz, C$\mathbf{H_2}$), 3.03 (q, *J* = 6.8 Hz, C**H**), 2.43 (br s, N**H**), 1.08 (d, *J* = 6.7 Hz, C$\mathbf{H_3}$); $^{13}$C NMR (75 MHz, DMSO–$d_6$) $\delta$ 177.4 (**C**(O)), 140.8, 139.3, 137.6, 132.6, 129.3, 129.0, 128.7, 126.9 (Ar**C**), 56.9 (**C**H), 51.1 (**C**$H_2$), 19.8 (**C**$H_3$).

**(*S*)-2-(((2'-(Trifluoromethyl)-[1,1'-biphenyl]-4-yl)methyl)amino)propanamide (11h)**

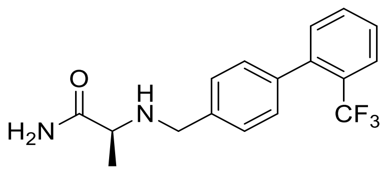

Using method B, L-alaninamide hydrochloride (**4**) (0.15 g, 1.34 mmol), triethylamine (0.23 mL, 1.68 mmol) **2h** (0.28 g, 1.11 mmol) in MeOH (1.22 mL) and sodium cyanoborohydride (0.30 g, 4.48 mmol) in MeOH (1.12 mL) gave **11h** as a white solid (0.17 g, 48%); $R_f$ = 0.15 (EtOAc); mp 109–111 °C; $^1$H NMR (400 MHz, DMSO–$d_6$) $\delta$ 7.82 (d, $J$ = 7.6 Hz, 1Ar**H**), 7.70 (t, $J$ = 7.6 Hz, 1Ar**H**), 7.59 (t, $J$ = 7.6 Hz, 1Ar**H**), 7.32–7.45 (m, 3Ar**H,** C(O)N**H**H'), 7.24 (d, $J$ = 7.8 Hz, 2Ar**H**), 7.00 (br s, C(O)NH**H'**), 3.60 and 3.73 (AB$_q$, $J$ = 13.7 Hz, C**H**$_2$), 3.05 (q, $J$ = 6.8 Hz, C**H**), 2.48 (br s, N**H**), 1.16 (d, $J$ = 6.8 Hz, C**H**$_3$); $^{13}$C NMR (75 MHz, DMSO–$d_6$) $\delta$ 177.4 (**C**(O)), 141.2, 140.7, 138.1, 132.7, 128.9, 128.4, 127.8, 127.3 (q, $J_{C–F}$ = 29.0 Hz), 126.4, 124.7 (q, $J_{C–F}$ = 273.6 Hz) (Ar**C**), 57.2 (**C**H), 51.2 (**C**H$_2$), 19.8 (**C**H$_3$).

**(*S*)-2-(((3'-(Trifluoromethyl)-[1,1'-biphenyl]-4-yl)methyl)amino)propanamide (11i)**

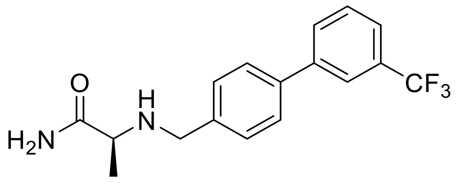

Using method B, L-alinamide hydrochloride (**4**) (2.11 g, 16.91 mmol), triethylamine (2.34mL, 16.91 mmol), **2i** (3.30 g, 14.10 mmol), and sodium cyanoborohydride (1.40 g, 21.14 mmol) in MeOH (14.09 mL) gave **11i** as a white solid (1.15 g, 26%); $R_f$ = 0.15 (EtOAc); mp 106–108 °C; $^1$H NMR (400 MHz, DMSO–$d_6$) $\delta$ 7.94–7.99 (m, 2Ar**H**), 7.69–7.71 (m, 4Ar**H**), 7.45–7.47 (m, 2Ar**H**), 7.36 (br s, C(O)N**H**H'), 7.02 (br s, C(O)NH**H'**), 3.60 and 3.75 (AB$_q$, $J$ = 13.8 Hz, NHC**H$_2$**Ar), 3.04 (q, $J$ = 6.8 Hz, C**H**), 1.16 (d, $J$ = 6.8 Hz, C**H$_3$**); $^{13}$C NMR (75 MHz, DMSO–d$_6$) $\delta$ 177.5 (**C**(O)), 141.6, 141.3, 137.4, 131.0, 130.4, 130.3 (q, $J_{C–F}$ = 31.2 Hz), 129.1, 127.1, 124.7 (q, $J_{C–F}$ = 270.8 Hz), 124.2 (q, $J_{C–F}$ = 3.6 Hz) 123.3 (q, $J_{C–F}$ = 3.7 Hz) (Ar**C**), 57.0 (**C**H), 51.1 (**C**H$_2$), 19.8 (**C**H$_3$).

**(*S*)-2-(((4'-(Trifluoromethyl)-[1,1'-biphenyl]-4-yl)methyl)amino)propanamide (11j)**

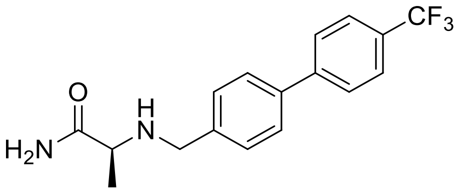

Using method B, L-alaninamide hydrochloride (**4**) (0.30 g, 2.4 mmol), triethylamine (0.42 mL, 4.12 mmol), **2j** (0.50 g, 2.0 mmol) in MeOH (2.17 mL), and sodium cyanoborohydride (0.30 g, 3.0 mmol) in MeOH (2.0 mL) gave **11j** as a white solid (0.33 g, 53%); $R_f$ = 0.10 (EtOAc); mp 144–147 °C; $^1$H NMR (300 MHz, DMSO–$d_6$) $\delta$ 7.89 (d, $J$ = 8.1 Hz, 2Ar**H**), 7.80 (d, $J$ = 8.3 Hz, 2Ar**H**), 7.69 (d, $J$ = 7.9 Hz, 2Ar**H**), 7.47 (d, $J$ = 8.0 Hz, 2Ar**H**), 7.34 (br s, C(O)N**H**H’), 6.99 (br s, C(O)NH**H**’), 3.60 and 3.75 ($AB_q$, $J$ = 13.8 Hz, C**H2**), 3.04 (q, $J$ = 6.8 Hz, C**H**), 1.15 (d, $J$ = 6.8 Hz, C**H3**), the remaining peak was not detected and is believed to overlap with $H_2O$ signals; $^{13}$C NMR (75 MHz, DMSO–$d_6$) $\delta$ 177.4 (**C**(O)), 144.5, 141.6, 137.4, 129.1, 128.1 (q, $J_{C–F}$ = 31.8 Hz), 127.7, 127.2, 126.2 (q, $J_{C–F}$ = 3.8 Hz), 124.8 (q, $J_{C–F}$ = 270.1 Hz) (Ar**C**), 57.0 (**C**H), 51.1 (**C**$H_2$), 19.8 (**C**$H_3$).

**(*S*)-2-(((3'-(Trifluoromethoxy)-[1,1'-biphenyl]-4-yl)methyl)amino)propanamide (11k)**

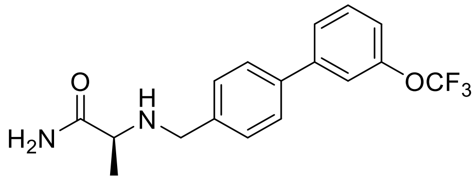

Using method B, L-alaninamide hydrochloride (**4**) (0.28 g, 2.25 mmol), triethylamine (0.39 mL, 2.82 mmol), **2k** (0.50 g, 1.88 mmol) in MeOH (2.04 mL) and sodium cyanoborohydride (0.17 g, 2.57 mmol) in MeOH (1.71 mL) gave **11k** as a white solid (0.34 g, 54%); $R_f$ = 0.10 (EtOAc); mp 95–97 °C; $^1$H NMR (300 MHz, DMSO–$d_6$) $\delta$ 7.55–7.76 (m, 5Ar**H**), 7.44 (d, $J$ = 7.8 Hz, 2Ar**H**), 7.30–7.38 (m, 1Ar**H**, C(O)N**H**H’), 6.99 (br s, C(O)NH**H’**), 3.59 and 3.74 ($AB_q$, $J$ = 13.7 Hz, C**H**$_2$), 3.03 (q, $J$ = 6.8 Hz, C**H**), 2.43 (br s, N**H**), 1.15 (d, $J$ = 6.8 Hz, C**H**$_3$); $^{13}$C NMR (75 MHz, $CDCl_3$) $\delta$ 178.0 (**C**(O)), 149.8(**C**O$CF_3$), 142.9, 139.4, 138.7, 130.1 128.6, 127.3, 125.4, 120.6 (q, $J_{C–F}$ = 255.7 Hz), 119.6 (Ar**C**), 57.7(**C**H), 52.2 (**C**$H_2$), 19.7 (**C**$H_3$).

**(*S*)-2-(((4'-(Trifluoromethoxy)-[1,1'-biphenyl]-4-yl)methyl)amino)propanamide (11l)**

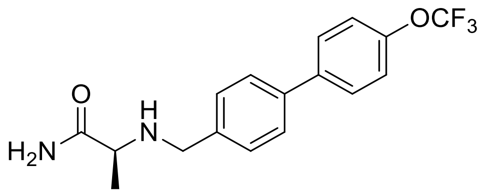


Using method B, L-alaninamide hydrochloride (**4**) (0.23 g, 1.81 mmol), triethylamine (0.25 mL, 1.81 mmol), **2l** (0.40 g, 1.51 mmol) and sodium cyanoborohydride (0.15 g, 2.26 mmol) in MeOH (1.5 mL) gave compound **11l** as a white solid (0.32 g, 62%); $R_f$ = 0.10 (EtOAc); mp 129–132 °C; $^1$H NMR (400 MHz, DMSO–$d_6$) $\delta$ 7.78 (d, $J$ = 8.6 Hz, 2Ar**H**), 7.63 (d, $J$ = 8.0 Hz, 2Ar**H**), 7.41–7.53 (m, 4Ar**H**), 7.34 (br s, C(O)N**H**H'), 7.00 (br s, C(O)NH**H'**), 3.59 and 3.73 ($AB_q$, $J$ = 13.7 Hz, C**H**$_2$), 2.98–3.11 (m, C**H**), 2.42 (br s, N**H**), 1.15 (d, $J$ = 6.8 Hz, C**H**$_3$); $^{13}$C NMR (75 MHz, $CDCl_3$) $\delta$ 178.1 (**C**(O)), 148.7 (**C**$OCF_3$), 139.6, 139.1, 128.9, 128.6, 128.4, 127.3, 121.3, 120.6 (q, $J_{C–F}$ = 255.4 Hz) (Ar**C**), 57.7 (**C**H), 52.2 (**C**$H_2$), 19.7 (**C**$H_3$).

**(*S*)-2-(((4'-Methoxy-[1,1'-biphenyl]-4-yl)methyl)amino)propanamide (11n)**

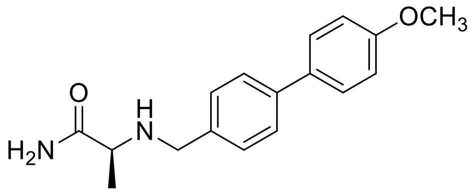


Using method B, L-alaninamide hydrochloride (**4**) (0.35 g, 2.83 mmol), triethylamine (0.49 mL, 3.53 mmol), **2n** (0.50 g, 2.36 mmol) in MeOH (2.56 mL), and sodium cyanoborohydride (0.23 g, 3.53 mmol) in MeOH (2.36 mL) gave **11n** as a white solid (0.11 g, 17%); $R_f$ = 0.15 (EtOAc); mp 160–162 ˚C; $^1$H NMR (300 MHz, DMSO–$d_6$) $\delta$ 7.54–7.60 (m, 4Ar**H**), 7.37 (d, $J$ = 7.92 Hz, 2Ar**H**), 7.33 (br s, C(O)N**H**H'), 6.96–7.09(m, 2Ar**H**, C(O)NH**H'**), 3.79 (s, OC**H**$_3$), 3.56 and 3.76 ($AB_q$, $J$ = 13.8 Hz, C**H**$_2$), 3.02 (q, $J$ = 6.4 Hz, C**H**), 2.36 (br s, N**H**), 1.15 (d, $J$ = 6.8 Hz, C**H**$_3$); $^{13}$C NMR (75 MHz, $CDCl_3$) $\delta$ 178.1 (**C**(O)), 159.2, 140.0, 137.9, 133.3, 128.4, 128.1, 126.9, 114.3 (Ar**C**), 57.8 (**C**H), 55.4 (O**C**$H_3$), 52.2 (**C**$H_2$), 19.7 (**C**$H_3$).

**(*S*)-2-(((4'-Isobutyl-[1,1'-biphenyl]-4-yl)methyl)amino)propanamide (11o)**

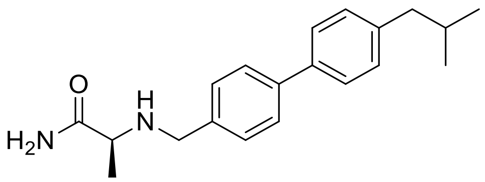

Using method B, L-alaninamide hydrochloride (**4**) (0.28 g, 2.52 mmol), triethylamine (0.44 mL, 3.15 mmol), **2o** (0.50 g, 2.10 mmol) in MeOH (2.28 mL), and sodium cyanoborohydride (0.56 g, 8.39 mmol) in MeOH (2.10 mL) gave **11o** as a white solid (0.40 g, 62%); $R_f$ = 0.20 ($CH_2Cl_2$/EtOAc 1/1); mp 149–150 ˚C; $^1$H NMR (300 MHz, DMSO–$d_6$) $\delta$ 7.54–7.60 (m, 4Ar**H**), 7.37 (d, *J* = 7.9 Hz, 2Ar**H**), 7.33 (br s, C(O)N**H**H'), 6.96–7.09 (m, 2Ar**H**, C(O)NH**H'**), 3.79 (s, OC**H**$_3$), 3.57 and 3.72 ($AB_q$, *J* = 13.8 Hz, C**H**$_2$), 3.02 (q, *J* = 6.4 Hz, C**H**), 2.36 (br s, N**H**), 1.15 (d, *J* = 6.8 Hz, C**H**$_3$); $^{13}$C NMR (75 MHz, DMSO–$d_6$) $\delta$ 177.4 (**C**(O)), 140.7, 140.0, 138.9, 138.0, 130.0, 128.9, 126.7, 126.6 (Ar**C**), 57.0 (C(O)**C**H), 51.2 (NH**C**$H_2$Ar), 44.7 ($CH_2$**C**H$(CH_3)_2$), 30.1 (Ph**C**$H_2$CH), 22.6 (2(**C**$H_3$)$_2$), 19.8 (C(O)CH**C**$H_3$).

**(*R*)-2-(((4'-(Trifluoromethyl)-[1,1'-biphenyl]-4-yl)methyl)amino)propanamide (12j)**

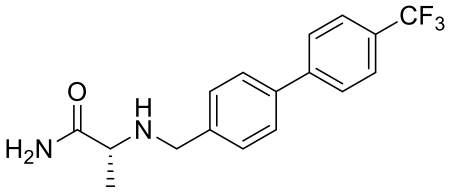

Using method B, D-alaninamide hydrochloride (**5**) (0.15 g, 1.20 mmol), triethylamine (0.21 mL, 1.50 mmol), **2j** (0.25 g, 1.00 mmol) in MeOH (1.09 mL) and sodium cyanoborohydride (0.26 g, 4.00 mmol) in MeOH (1.00 mL) gave **12j** as a white solid (0.21 g, 64%); $R_f$ = 0.10 (EtOAc); mp 143–145 ˚C; $^1$H NMR (300 MHz, DMSO–$d_6$) $\delta$ 7.89 (d, *J* = 8.3 Hz, 2Ar**H**), 7.80 (t, *J* = 8.3 Hz, 2Ar**H**), 7.69 (d, *J* = 8.1 Hz, 2Ar**H**), 7.47 (d, *J* = 8.1 Hz, 2Ar**H**), 7.35 (br s, C(O)N**H**H'), 7.00 (br s, C(O)NH**H'**), 3.60 and 3.75 ($AB_q$, *J* = 13.8 Hz, NHC**H**$_2$Ar), 3.03 (q, *J* = 6.8 Hz, C**H**), 2.44 (br s, N**H**), 1.15 (d, *J* = 6.8 Hz, C**H**$_3$); $^{13}$C NMR (75 MHz, $CDCl_3$) $\delta$ 177.9 (**C**(O)), 144.3, 139.7, 138.8, 129.4 (q, $J_{C–F}$ = 32.3 Hz), 128.6, 127.5, 127.3, 125.8 (q, $J_{C–F}$ = 3.7 Hz), 124.3 (q, $J_{C–F}$ = 270.1 Hz) (Ar**C**), 57.7 (**C**H), 52.2 (**C**$H_2$), 19.7 (**C**$H_3$).

**(*S*)-2-(((4'-(Trifluoromethyl)-[1,1'-biphenyl]-4-yl)methyl)amino)butanamide (13j)**

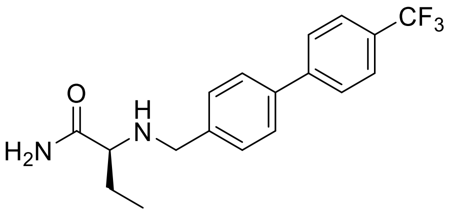

Using method B, (*S*)-2-aminobutanamide hydrochloride (**6**) (0.33 g, 2.40 mmol), triethylamine (0.42 mL, 3.00 mmol), **2j** (0.50 g, 2.00 mmol) in MeOH (2.17 mL), and sodium cyanoborohydride (0.53 g, 8.00 mmol) in MeOH (4.00 mL) gave **13j** as a white solid (0.26 g, 39%); $R_f$ = 0.15 (EtOAc/MeOH 30/1); mp 149–152 ˚C; $^1$H NMR (300 MHz, DMSO–$d_6$) $\delta$ 7.88 (d, *J* = 8.1 Hz, 2Ar**H**), 7.80 (d, *J* = 8.1 Hz, 2Ar**H**), 7.69 (d, *J* = 7.4 Hz, 2Ar**H**), 7.47 (d, *J* = 7.4 Hz, 2Ar**H**), 7.37 (br s, C(O)N**H**H'), 7.04 (br s, C(O)NH**H'**), 3.57 and 3.78 ($AB_q$, *J* = 13.9 Hz, NHC**H**$_2$Ar), 2.88 (t, *J* = 6.3 Hz, C(O)**C**H), 2.35 (br s, N**H**), 1.43–1.60 (m, CHC**H**$_2$CH$_3$), 0.88 (t, *J* = 7.2 Hz, C**H**$_3$); $^{13}$C NMR (75 MHz, DMSO–*d*$_6$) $\delta$ 176.7 (**C**(O)), 144.5, 141.7, 137.3, 129.1, 128.1 (q, $J_{C-F}$ = 31.7 Hz), 127.7, 127.2, 126.1 (q, $J_{C-F}$ = 3.8 Hz), 124.8 (q, $J_{C-F}$ = 270.2 Hz, **C**CF$_3$) (Ar**C**), 62.9 (C(O)**C**H), 51.2 (NH**C**H$_2$Ar), 26.8 (**C**H$_2$CH$_3$), 10.9 (CH$_2$**C**H$_3$).

**(*S*)-2-(((4'-(Trifluoromethoxy)-[1,1'-biphenyl]-4-yl)methyl)amino)butanamide (13l)**

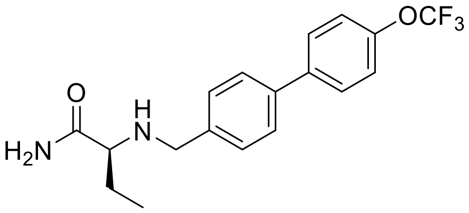

Using method B, (*S*)-2-aminobutanamide hydrochloride (**6**) (0.31 g, 2.25 mmol), triethylamine (0.39 mL, 2.82 mmol), **2l** (0.50 g, 1.88 mmol) in MeOH (2.00 mL), and sodium cyanoborohydride (0.50 g, 7.51 mmol) in MeOH (3.76 mL) gave **13l** as a white solid (0.42 g, 64%); $R_f$ = 0.20 (EtOAc); mp 143–145 ˚C; $^1$H NMR (300 MHz, DMSO–$d_6$) $\delta$ 7.77 (d, *J* = 8.8 Hz, 2Ar**H**), 7.62 (d, *J* = 8.2 Hz, 2Ar**H**), 7.40–7.48 (m, 4Ar**H**), 7.38 (br s, C(O)N**H**H'), 7.04 (br s, C(O)NH**H'**), 3.56 and 3.77 ($AB_q$, *J* = 13.8 Hz, NHC**H**$_2$Ar), 2.89 (t, *J* = 6.5 Hz, C(O)**C**H), 2.37 (br s, N**H**), 1.47–1.60 (m, CHC**H**$_2$CH$_3$), 0.88 (t, *J* = 7.4 Hz, C**H**$_3$); $^{13}$C NMR (75 MHz, DMSO–*d*$_6$) $\delta$ 176.7 (**C**(O)), 148.2 (**C**OCF$_3$), 141.0, 139.9, 137.5, 129.0, 128.8, 127.0, 122.3, 120.6 (q, $J_{C-F}$ = 254.5 Hz) (Ar**C**), 63.0 (C(O)**C**H), 51.2 (NH**C**H$_2$Ar), 26.8 (**C**H$_2$CH$_3$), 10.8 (CH$_2$**C**H$_3$).

**(*S*)-3-Methyl-2-(((3'-(trifluoromethyl)-[1,1'-biphenyl]-4-yl)methyl)amino)butanamide (14i)**

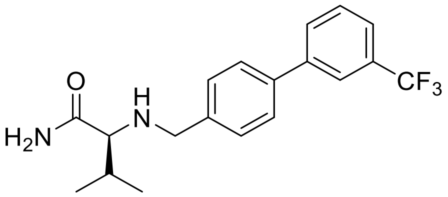


Using method B, L-valinamide hydrochloride (**7**) (0.29 g, 1.92 mmol), triethylamine (0.33 mL, 2.40 mmol) **2i** (0.40 g, 1.60 mmol) in MeOH (1.63 mL) and sodium cyanoborohydride (0.42 g, 6.39 mmol) in MeOH (3.20 mL) gave **14i** as a white solid (0.31 g, 55%); $R_f$ = 0.25 ($CH_2Cl_2$/EtOAc 4/1); mp 119–121 ˚C; $^1$H NMR (300 MHz, DMSO–$d_6$) $\delta$ 7.91–8.03 (m, 2Ar**H**), 7.63–7.79 (m, 4Ar**H**), 7.46 (d, *J* = 8.0 Hz, 2Ar**H**), 7.38 (br s, C(O)N**H**H’), 7.07 (br s, C(O)NH**H’**), 3.52 and 3.81 ($AB_q$, *J* = 13.9 Hz, C**H**$_2$), 2.69 (d, *J* = 6.2 Hz, C(O)C**H**), 2.34 (br s, N**H**), 1.69–1.85 (m, CHC**H**($CH_3$)$_2$), 0.84–0.98 (m, CH(C**H**$_3$) (C**H’**$_3$)); $^{13}$C NMR (75 MHz, DMSO–$d_6$) $\delta$ 176.3 (**C**(O)), 141.6, 141.5, 137.3, 131.1, 130.5, 130.2 (q, $J_{C–F}$ = 31.5 Hz), 129.1, 127.2, 124.7 (q, $J_{C–F}$ = 270.8 Hz), 124.3, 123.3 (q, $J_{C–F}$ = 3.7 Hz) (Ar**C**), 67.3 (C(O)CH), 51.5 (**C**H$_2$), 31.5 (C(O)CH**C**H), 20.1 (CH(**C**H$_3$) (C’H$_3$), 19.8 (CH(CH$_3$) (**C’**H$_3$)).

**(*S*)-3-Methyl-2-(((4'-(trifluoromethyl)-[1,1'-biphenyl]-4-yl)methyl)amino)butanamide (14j)**

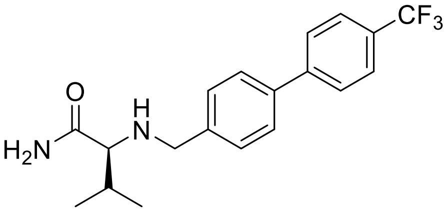


Using method B, L-valinamide hydrochloride (**7**) (0.08 g, 0.72 mmol), triethylamine (0.04 mL, 0.90 mmol) **2j** (0.15 g, 0.60 mmol) in MeOH (0.60 mL) and sodium cyanoborohydride (0.24 g, 3.60 mmol) in MeOH (0.60 mL) gave **14j** as a white solid (0.11 g, 51%); $R_f$ = 0.20 (MC/EtOAc 3/2); mp 138–140 ˚C; $^1$H NMR (400 MHz, DMSO–$d_6$) $\delta$ 7.89 (d, *J* = 8.3 Hz, 2Ar**H**), 7.80 (d, *J* = 8.3 Hz, 2Ar**H**), 7.69 (d, *J* = 8.1 Hz, 2Ar**H**), 7.47 (d, *J* = 8.1 Hz, 2Ar**H**),

7.37 (br s, C(O)N**H**H’), 7.06 (br s, C(O)NH**H’**), 3.53 and 3.81 ($AB_q$, $J$ = 13.9 Hz, C**H**$_2$), 2.69 (d, $J$ = 6.5 Hz, C(O)C**H**), 2.29 (br s, N**H**), 1.70–1.83 (m, CHC**H**$(CH_3)_2$), 0.85–0.98 (m, CH(C**H**$_3$) (C**H’**$_3$)); $^{13}$C NMR (75 MHz, DMSO–$d_6$) $\delta$ 176.3 (**C**(O)), 144.5, 141.8, 137.3, 129.1, 128.1 (q, $J_{C-F}$ = 31.6 Hz), 127.7, 127.2, 126.2 (q, $J_{C-F}$ = 3.7 Hz), 124.9 (q, $J_{C-F}$ = 270.1 Hz) (Ar**C**), 67.3 (C(O)**C**H), 51.5 (**C**H$_2$), 31.5 (C(O)CH**C**H), 20.1 (CH(**C**H$_3$) (C’H$_3$), 19.2 (CH(CH$_3$) (**C’**H$_3$)).

**(*S*)-2-(((3'-Fluoro-[1,1'-biphenyl]-4-yl)methyl)amino)-4-methylpentanamide (15c)**

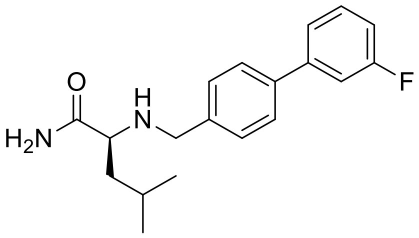


Using method B, L-leucinamide hydrochloride (**8**) (0.31 g, 1.80 mmol), triethylamine (0.31 mL, 2.25 mmol), **2c** (0.30 g, 1.50 mmol) in MeOH (1.63 mL), and sodium cyanoborohydride (0.15 g, 2.25 mmol) in MeOH (1.5 mL) gave **15c** as a white solid (0.19 g, 40%); $R_f$ = 0.30 (only EtOAc); mp 149–151 ˚C; $^1$H NMR (300MHz, DMSO–$d_6$) $\delta$ 7.65 (d, $J$ = 7.7 Hz, 2Ar**H**), 7.40–7.51 (m, 5Ar**H,** C(O)N**H**H’), 7.15–7.20 (m, 1Ar**H**), 6.99 (br s, C(O)NH**H’**), 3.53 and 3.76 ($AB_q$, $J$ = 13.8 Hz, NHC**H**$_2$Ar), 2.97 (t, $J$ = 7.1 Hz, C(O)C**H**NH), 2.27 (br s, N**H**), 1.69–1.85 (m, C**H**$(CH_3)_2$) 1.25–1.43 (m, CHC**H**$_2$CH), 0.88 (d, $J$ = 6.4 Hz, C**H**$_3$), 0.80 (d, $J$ = 6.4 Hz, C**H**$_3$); $^{13}$C NMR (75 MHz, $CDCl_3$) $\delta$ 178.2 (**C**(O)), 163.2 (q, $J_{C-F}$ = 243.9 Hz), 143.1 (q, $J_{C-F}$ = 7.6 Hz), 139.3, 139.0, 130.3 (q, $J_{C-F}$ = 31.6 Hz), 128.7, 127.3, 122.7 (q, $J_{C-F}$ = 2.7 Hz), 114.1 (q, $J_{C-F}$ = 21.2 Hz), 113.9 (q, $J_{C-F}$ = 21.8 Hz) (Ar**C**), 60.9 (C(O)**C**H), 52.5 (NH**C**H$_2$Ar), 43.0 ($CH_2$**C**H$(CH_3)_2$), 25.1 (CH**C**H$_2$CH), 23.3 (CH(**C**H$_3$)(C’H$_3$)), 21.8 (CH(CH$_3$)(**C’**H$_3$)).

**(*S*)-4-Methyl-2-(((3'-(trifluoromethyl)-[1,1'-biphenyl]-4-yl)methyl)amino)pentanamide (15i)**

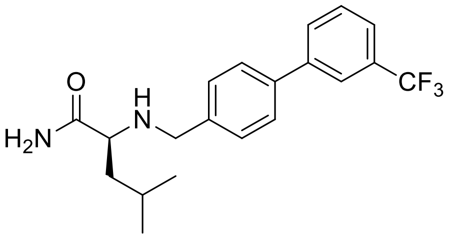


Using method B, L-leucinamide hydrochloride (**8**) (0.32 g, 1.92 mmol), triethylamine (0.33 mL, 2.40 mmol) **2i** (0.40 g, 1.60 mmol) in MeOH (1.74 mL) and sodium cyanoborohydride (0.42 g, 6.39 mmol) in MeOH (1.60 mL) gave **15i** as a white solid (0.27 g, 46%); $R_f$ = 0.25 (MC/EtOAc 4/1); mp 109–110 ˚C; $^1$H NMR (300 MHz, DMSO–$d_6$) $\delta$ 7.92–8.02 (m, 2Ar**H**), 7.65–7.76 (m, 4Ar**H**), 7.36–7.49 (m, 2Ar**H**, C(O)N**H**H'), 7.00 (br s, C(O)NH**H'**), 3.55 and 3.78 (AB$_q$, $J$ = 13.8 Hz, NHC**H**$_2$Ar), 2.98 (t, $J$ = 7.1 Hz, C(O)C**H**NH), 2.34 (br s, N**H**), 1.66–1.89 (m, CH$_2$C**H**(CH$_3$)$_2$), 1.39–1.42 (m, CHC**H**$_2$CH), 0.88 (d, $J$ = 6.6 Hz, CH(C**H**$_3$)(CH'$_3$)), 0.81 (d, $J$ = 6.6 Hz, CH(CH$_3$)(C**H'**$_3$)); $^{13}$C NMR (75 MHz, DMSO–$d_6$) $\delta$ 177.4 (**C**(O)), 141.5 (q, $J_{C–F}$ = 12.1 Hz), 137.3, 131.0, 130.5, 130.3 (q, $J_{C–F}$ = 94.2 Hz), 129.1, 127.2, 124.7 (q, $J_{C–F}$ = 270.9 Hz), 124.2, 123.3 (Ar**C**), 60.3 (C(O)**C**H), 51.2 (NH**C**H$_2$Ar), 43.3 (CH$_2$**C**H(CH$_3$)$_2$), 24.8 (CH**C**H$_2$CH), 23.4 (CH(**C**H$_3$)(C'H$_3$)), 22.7 (CH(CH$_3$)(**C'**H$_3$)).

**(*S*)-4-Methyl-2-(((4'-(trifluoromethyl)-[1,1'-biphenyl]-4-yl)methyl)amino)pentanamide (15j)**

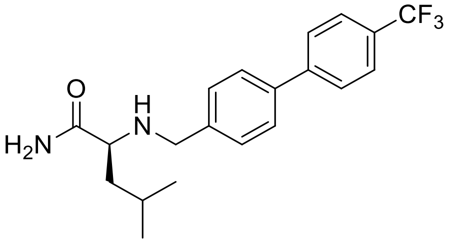


Using method B, L-leucinamide hydrochloride (**8**) (0.34 g, 2.88 mmol), triethylamine (0.50 mL, 3.60 mmol) **2j** (0.60 g, 2.40 mmol) in MeOH (2.61 mL) and sodium cyanoborohydride (0.63 g, 9.59 mmol) in MeOH (2.40 mL) gave **15j** as a white solid (0.18 g, 22%); $R_f$ = 0.25 ($CH_2Cl_2$/EtOAc 3/1); mp 145–148 ˚C; $^1$H NMR (300 MHz, DMSO–$d_6$) $\delta$ 7.88 (d, $J$ = 8.4 Hz, 2Ar**H**), 7.80 (d, $J$ = 8.4 Hz, 2Ar**H**), 7.69 (d, $J$ = 8.1 Hz, 2Ar**H**), 7.46 (d, $J$ = 8.1 Hz, 2Ar**H**), 7.41 (br s, C(O)N**H**H'), 7.00 (br s, C(O)NH**H'**), 3.55 and 3.78 (AB$_q$, $J$ = 13.8 Hz, NHC**H**$_2$Ar), 2.98 (t, $J$ = 7.1 Hz, C(O)C**H**NH), 2.33 (br s, N**H**), 1.69–1.86 (m, CH$_2$C**H**(CH$_3$)$_2$), 1.29–1.43

(m, CHC**H**$_2$CH), 0.88 (d, *J* = 6.6 Hz, CH(C**H$_3$**)(CH'$_3$)), 0.80 (d, *J* = 6.6 Hz, CH(CH$_3$)(C**H'$_3$**)); $^{13}$C NMR (75 MHz, DMSO–$d_6$) $\delta$ 177.4 (**C**(O)), 144.5, 141.7, 137.3, 129.1, 128.1 (q, $J_{C–F}$ = 31.6 Hz), 127.7, 127.2, 126.2 (q, $J_{C–F}$ = 3.8 Hz), 124.9 (q, $J_{C–F}$ = 270.2 Hz) (Ar**C**), 60.2 (C(O)**C**H), 51.2 (NH**C**H$_2$Ar), 43.3 (CH$_2$**C**H(CH$_3$)$_2$), 24.8 (CH**C**H$_2$CH), 23.4 (CH(**C**H$_3$)(C'H$_3$)), 22.7 (CH(CH$_3$)(**C'**H$_3$)).

**(*S*)-4-Methyl-2-(((3'-(trifluoromethoxy)-[1,1'-biphenyl]-4-yl)methyl)amino)pentanamide (15k)**

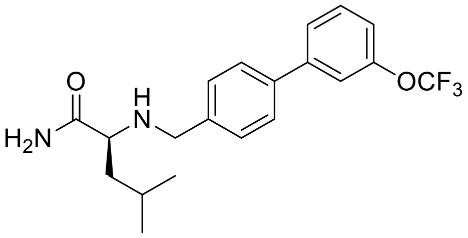


Using method B, L-leucinamide hydrochloride (**8**) (0.45 g, 2.70 mmol), triethylamine (0.47 mL, 3.38 mmol), **2k** (0.60 g, 2.25 mmol) in MeOH (2.45 mL), and sodium cyanoborohydride (0.22 g, 3.38 mmol) in MeOH (2.25 mL) gave **15k** as a white solid (0.25 g, 30%); R$_f$ = 0.30 (n-Hexane/EtOAc 10/1); mp 116–119 °C; $^1$H NMR (300 MHz, DMSO–$d_6$) $\delta$ 7.55–7.76 (m, 5Ar**H**), 7.31–7.48 (m, 3Ar**H**, C(O)N**H**H'), 7.00 (br s, C(O)NH**H'**), 3.53 and 3.77 (AB$_q$, *J* = 13.8 Hz, NHC**H$_2$**Ar), 2.97 (t, *J* = 6.9 Hz, C(O)C**H**), 2.28 (br s, N**H**), 1.69–1.85 (m, CH$_2$C**H**(CH$_3$)$_2$), 1.25–1.43 (m, CHC**H$_2$**CH), 0.88 (d, *J* = 6.6 Hz, CH(C**H$_3$**)(CH'$_3$)), 0.80 (d, *J* = 6.6 Hz, CH(CH$_3$)(C**H'$_3$**)$_2$); $^{13}$C NMR (75 MHz, CDCl$_3$) $\delta$ 178.2 (**C**(O)), 149.7, 142.9, 139.5, 138.7, 130.1, 128.7, 127.3, 125.4, 120.6 (q, $J_{C–F}$ = 255.4 Hz), 119.6 (q, $J_{C–F}$ = 4.0 Hz) (Ar**C**), 60.9 (C(O)**C**H), 52.4 (NH**C**H$_2$Ar), 43.0 (CH$_2$**C**H(CH$_3$)$_2$), 25.1 (CH**C**H$_2$CH), 23.2 (CH(**C**H$_3$)(C'H$_3$)), 21.8 (CH(CH$_3$)(**C'**H$_3$)).

**(*S*)-2-(((3'-Methoxy-[1,1'-biphenyl]-4-yl)methyl)amino)-4-methylpentanamide (15m)**

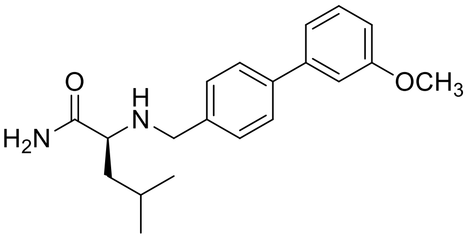


Using method D, L-leucinamide hydrochloride (**8**) (0.28 g, 1.70 mmol), triethylamine (0.30 mL, 2.12 mmol), **2m** (0.30 g, 1.41 mmol) in MeOH (1.54 mL), and sodium cyanoborohydride (0.14 g, 2.12 mmol) in MeOH (1.40 mL) gave **15m** as a white solid (0.22 g, 46%); $R_f$ = 0.20 (n-Hexane/EtOAc 1/2); mp 112–114 ˚C; $^1$H NMR (300 MHz, DMSO–$d_6$) $\delta$ 7.61 (d, $J$ = 7.8 Hz, 2Ar**H**), 7.31–7.46 (m, 3ArH, C(O)N**H**H'), 7.13–7.25 (m, 2Ar**H**), 7.00 (br s, C(O)N**H**H'), 6.87–6.96 (m, 1Ar**H**), 3.82 (s, OC**H$_3$**), 3.52 and 3.76 (AB$_q$, $J$ = 13.7 Hz, NHC**H$_2$**Ar), 2.97 (t, $J$ = 7.0 Hz), 2.28 (br s, N**H**), 1.68–1.88 (m, CH$_2$C**H**(CH$_3$)$_2$), 1.22–1.45 (m, CHC**H$_2$**CH), 0.88 (d, $J$ = 6.6 Hz, CH(C**H$_3$**)(CH'$_3$)), 0.80 (d, $J$ = 6.6 Hz, CH(CH$_3$)(C**H'$_3$**)); $^{13}$C NMR (100 MHz, CDCl$_3$) $\delta$ 177.4 (**C**(O)), 160.2 (**C**OCH$_3$), 142.1, 140.6, 138.9, 130.5, 128.9, 127.0, 119.3, 113.3, 122.5 (Ar**C**), 60.2 (C(O)**C**H), 55.6 (NH**C**H$_2$Ar), 51.2 (CH$_2$**C**H(CH$_3$)$_2$), 24.8 (CH**C**H$_2$CH), 23.5 (CH(**C**H$_3$)(C'H$_3$)), 22.7 (CH(CH$_3$)(**C'**H$_3$)).

**(2*S*,3*S*)-2-(((3'-Fluoro-[1,1'-biphenyl]-4-yl)methyl)amino)-3-methylpentanamide (16c)**

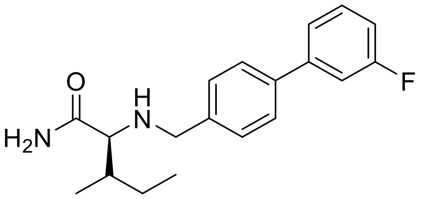


Using method B, L-isoleucinamide hydrochloride (**9**) (0.31 g, 1.80 mmol), triethylamine (0.31 mL, 2.25 mmol), **2c** (0.30 g, 1.50 mmol) in MeOH (1.63 mL), and sodium cyanoborohydride (0.15 g, 2.25 mmol) in MeOH (1.50 mL) gave **16c** as a white solid (0.25 g, 53%); $R_f$ = 0.15 (n-Hexane/EtOAc 1/1); mp 107–108 ˚C; $^1$H NMR (400 MHz, DMSO–$d_6$) $\delta$ 7.65 (d, $J$ = 8.0 Hz, 2Ar**H**), 7.48–7.51 (m, 3Ar**H**), 7.42 (d, $J$ = 8.0 Hz, 2Ar**H**), 7.36 (br s, C(O)N**H**H'), 7.15–7.19 (m, 1Ar**H**), 7.04 (br s, C(O)NH**H'**), 3.50 and 3.78 (AB$_q$, $J$ = 13.8 Hz, NHC**H$_2$**Ar), 2.73 (d, $J$ = 6.1 Hz, C(O)C**H**), 2.22 (br s, N**H**), 1.45–1.70 (m, CHC**H$_2$**CH$_3$), 1.08–1.20 (m, CHC**H**CH$_2$), 0.78–0.90 (m, CHC**H$_3$**, CH$_2$C**H$_3$**); $^{13}$C NMR (75 MHz, DMSO–$d_6$) $\delta$ 176.3 (**C**(O)), 163.2 (d, $J_{C–F}$ = 241.7 Hz), 143.0 (d, $J_{C–F}$ = 7.7 Hz), 141.3, 137.5 (d, $J_{C–F}$ = 2.1 Hz), 131.2 (d, $J_{C–F}$ = 8.5

Hz), 129.0, 127.0, 123.0 (d, $J_{C\text{–}F}$ = 2.3 Hz), 114.3 (d, $J_{C\text{–}F}$ = 21.0 Hz), 113.6 (d, $J_{C\text{–}F}$ = 21.8 Hz) (Ar**C**), 66.3 (C(O)**C**H), 51.5 (NH**C**$H_2$Ar), 38.0 (CH**C**H$CH_3$), 25.4 (CH**C**$H_2CH_3$), 16.3, 11.7 (CH**C**$H_3$, $CH_2$**C**$H_3$).

**(2*S*,3*S*)-2-(((3'-Chloro-[1,1'-biphenyl]-4-yl)methyl)amino)-3-methylpentanamide (16f)**

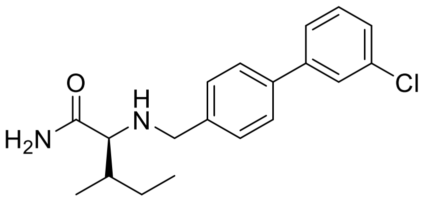


Using method B, L-isoleucinamide hydrochloride (**9**) (0.28 g, 1.66 mmol), triethylamine (0.29 mL, 2.08 mmol), **2f** (0.30 g, 1.38 mmol) in MeOH (1.51 mL), and sodium cyanoborohydride (0.14 g, 2.08 mmol) in MeOH (1.38 mL) gave **16f** as a white solid (0.25 mg, 56%); $R_f$ = 0.25 ($CH_2Cl_2$/EtOAc 1/1); mp 101–102 ˚C; $^1$H NMR (300 MHz, DMSO–$d_6$) $\delta$ 7.60–7.78 (m, 4Ar**H**), 7.35–7.53 (m, 4Ar**H**, C(O)N**H**H'), 7.06 (br s, C(O)NH**H'**), 3.50 and 3.78 ($AB_q$, $J$ = 13.8 Hz, NHC**$H_2$**Ar), 2.74 (d, $J$ = 6.7 Hz, C(O)C**H**), 2.25 (br s, N**H**), 1.45–1.73 (m, CHC**$H_2$**$CH_3$), 1.03–1.26 (m, CHC**H**$CH_2$), 0.77–0.92 (m, CHC**$H_3$**, $CH_2$C**$H_3$**); $^{13}$C NMR (75 MHz, $CDCl_3$) $\delta$ 176.6 (**C**(O)), 142.7, 139.4, 138.9, 134.7, 130.0, 128.7, 127.3, 127.3, 127.2, 125.2 (Ar**C**), 67.2 (C(O)**C**H), 53.0 (NH**C**$H_2$Ar), 38.3 (CH**C**H($CH_3$) ($CH_2CH_3$), 25.3 (CH**C**$H_2CH_3$), 15.9, 11.7 (CH**C**$H_3$, $CH_2$**C**$H_3$).

**(2*S*,3*S*)-2-(((4'-Chloro-[1,1'-biphenyl]-4-yl)methyl)amino)-3-methylpentanamide (16g)**

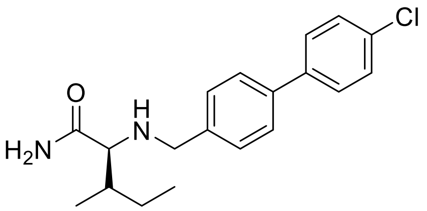


Using method B, L-isoleucinamide hydrochloride (**9**) (0.28 g, 1.66 mmol), triethylamine (0.29 mL, 2.08 mmol), **2g** (0.30 g, 1.38 mmol) in MeOH (1.51 mL), and sodium cyanoborohydride (0.14 g, 2.08 mmol) in MeOH (1.38 mL) gave **16g** as a white solid (0.20 mg, 44%); $R_f$ = 0.25 ($CH_2Cl_2$/EtOAc 1/1); mp 132–134 ˚C; $^1$H NMR (300 MHz, DMSO–$d_6$) $\delta$ 7.68 (d, $J$ = 8.4 Hz,

2Ar**H**), 7.61 (d, $J$ = 8.0 Hz, 2Ar**H**), 7.50 (d, $J$ = 8.4 Hz, 2Ar**H**), 7.42 (d, $J$ = 8.0 Hz, 2Ar**H**), 7.37 (br s, C(O)N**H**H'), 7.05 (br s, C(O)NH**H'**), 3.50 and 3.78 (AB$_q$, $J$ = 13.8 Hz, NHC**H$_2$**Ar), 2.73 (d, $J$ = 4.8 Hz, C(O)C**H**), 2.22 (br s, N**H**), 1.45–1.70 (m, CHC**H$_2$**CH$_3$), 1.02–1.25 (m, CHC**H**CH$_2$), 0.78–0.90 (m, CHC**H$_3$**, CH$_2$C**H$_3$**); $^{13}$C NMR (75 MHz, DMSO–$d_6$) $\delta$ 176.3 (**C**(O)), 141.0, 139.4, 137.6, 132.5, 129.3, 129.0, 128.7, 126.8 (Ar**C**), 66.2 (C(O)**C**H), 51.5 (NH**C**H$_2$Ar), 38.0 (CH**C**H(CH$_3$) (CH$_2$CH$_3$), 25.4 (CH**C**H$_2$CH$_3$), 16.3, 11.8 (CH**C**H$_3$, CH$_2$**C**H$_3$).

**(2*S*,3*S*)-3-Methyl-2-(((3'-(trifluoromethyl)-[1,1'-biphenyl]-4-yl)methyl)amino)pentanamide (16i)**

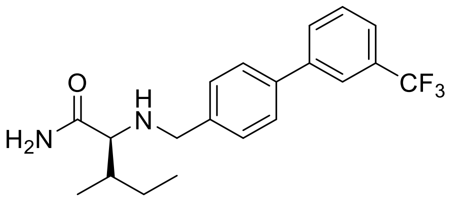


Using method B, L-isoleucinamide hydrochloride (**9**) (0.32 g, 1.92 mmol), triethylamine (0.33 mL, 2.40 mmol), **2i** (0.40 g, 1.60 mmol) in MeOH (1.74 mL), and sodium cyanoborohydride (0.42 g, 6.39 mmol) in MeOH (1.60 mL) gave **16i** as a white solid (0.18g, 31%); R$_f$ = 0.30 (CH$_2$Cl$_2$/EtOAc 4/1); mp 110–111 ˚C; $^1$H NMR (300 MHz, DMSO–$d_6$) $\delta$ 7.91–8.04 (m, 2Ar**H**), 7.65–7.78 (m, 4Ar**H**), 7.45 (d, $J$ = 8.1 Hz, 2Ar**H**), 7.38 (br s, C(O)N**H**H'), 7.05 (br s, C(O)NH**H'**), 3.52 and 3.80 (AB$_q$, $J$ = 13.9 Hz, NHC**H$_2$**Ar), 2.74 (d, $J$ = 6.7 Hz, C(O)C**H**), 2.30 (br s, N**H**), 1.32–1.43 (m, CHC**H$_2$**H$_3$), 1.02–1.25 (m, CHC**H**CH$_2$); $^{13}$C NMR (75 MHz, DMSO–$d_6$) $\delta$ 176.3 (**C**(O)), 141.6, 141.5, 137.3, 130.0, 130.4, 130.2 (q, $J_{C–F}$ = 31.4 Hz), 129.1, 127.1, 126.5, 124.7 (q, $J_{C–F}$ = 270.7 Hz), 124.2 (q, $J_{C–F}$ = 3.5 Hz), 123.3 (q, $J_{C–F}$ = 3.8 Hz) (Ar**C**), 66.3 (C(O)**C**H), 51.5 (NH**C**H$_2$Ar), 38.0 (CH**C**H(CH$_3$)(CH$_2$CH$_3$), 25.4 (CH**C**H$_2$CH$_3$), 16.3, 11.7 (CH**C**H$_3$, CH$_2$**C**H$_3$).

**General procedure for the preparation of methanesulfonic acid salt compounds (17a-23i) (Method C)**

The an EtOAc solution of free amine **(10a-16i)** was added methanesulfonic acid (1.0–1.2 equiv.) at room temperature for 1 h. The reaction mixture filtered *in vacuo* and washed with EtOAc and *n*-hexane. The washed filter cake was dried to give desired compounds without further purification.

**2-(([1,1'-Biphenyl]-4-ylmethyl)amino)acetamide methanesulfonate (17a)**

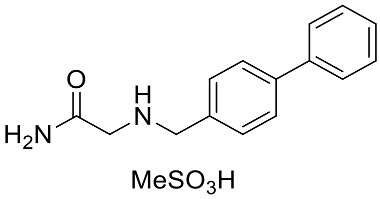


Using method C, to a solution of **10a** (0.15 g, 0.62 mmol), methanesulfonic acid (50.64 μL, 0.78 mmol) in EtOAc (1.24 mL) gave **17a** as a white solid (0.19 g, 90%); $R_f$ = 0.00 ($CH_2Cl_2$/MeOH 10/1); mp 233–236 ˚C; HPLC purity: 6.1 min, >99.9%; $^1$H NMR (300 MHz, DMSO–$d_6$) $\delta$ 9.20 (br s, $^+$N**H**$_2$), 7.86 (br s, C(O)N**H**H'), 7.67–7.79 (m, 4Ar**H**), 7.55–7.65(m, 2Ar**H**, C(O)NH**H'**), 7.48 (t, $J$ = 7.3 Hz, 2Ar**H**), 7.34–7.44 (m, 1Ar**H**), 4.22 (s, $^+NH_2C$**H**$_2$Ar), 3.68 (s, C(O)C**H**$_2$$^+NH_2$), 2.35 (s, SC**H**$_3$); $^{13}$C NMR (75 MHz, DMSO–$d_6$) $\delta$ 167.3 (**C**(O)), 141.2, 139.8, 131.2, 131.1, 129.5, 128.3, 127.4, 127.2 (Ar**C**), 50.0 (C(O)**C**H$^+NH_2$), 47.2 ($^+NH_2$**C**$H_2$Ar), 40.2 (S**C**$H_3$); LRMS (M + H)$^+$(ESI$^+$) 241.1 [M + H]$^+$ (calcd for $C_{15}H_{16}N_2OH^+$ 241.1)

**2-(((4'-Fluoro-[1,1'-biphenyl]-4-yl)methyl)amino)acetamide methanesulfonate (17d)**

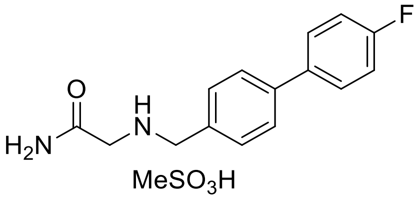


Using method C, to a solution of **10d** (0.10 g, 0.39 mmol), methanesulfonic acid (31.41 μL, 0.48 mmol) in EtOAc (0.77 mL) gave **17d** as a white solid (0.13 g, 97%); $R_f$ = 0.00 ($CH_2Cl_2$/MeOH 10/1); mp 238–242 ˚C; HPLC purity: 7.3 min, 99.6%; $^1$H NMR (300 MHz, DMSO–$d_6$) $\delta$ 9.20 (br s, $^+$N**H**$_2$), 7.85 (br s, C(O)N**H**H'), 7.68–7.81 (m, 3Ar**H**, C(O)NH**H'**), 7.54–7.63 (m, 3Ar**H**), 7.25–7.36 (m, 2Ar**H**), 4.21 (s, $^+NH_2C$**H**$_2$Ar), 3.67 (s, C(O)C**H**$_2$$^+NH_2$), 2.34 (s, SC**H**$_3$); $^{13}$C NMR (75 MHz, DMSO–$d_6$) $\delta$ 167.3 (**C**(O)), 162.5 (d, $J_{C–F}$ = 243.1 Hz), 140.2, 136.3 (d, $J_{C–F}$ = 3.1 Hz), 131.2, 131.1, 129.2 (d, $J_{C–F}$ = 8.2 Hz), 127.3, 116.3 (d, $J_{C–F}$ =

21.3 Hz) (Ar**C**), 49.9 (C(O)**C**H$^+$NH$_2$), 47.2 ($^+$NH$_2$**C**H$_2$Ar), 40.2 (S**C**H$_3$); LRMS (M + H)$^+$(ESI$^+$) 259.1 [M + H]$^+$ (calcd for $C_{15}H_{15}FN_2OH^+$ 259.1)

**2-(((3'-Chloro-[1,1'-biphenyl]-4-yl)methyl)amino)acetamide methanesulfonate (17f)**

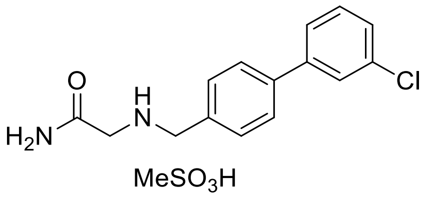


Using method C, to a solution of **10f** (0.26 g, 0.95 mmol), methanesulfonic acid (76.76 µL, 1.18 mmol) in EtOAc (1.89 mL) gave **17f** as a white solid (0.33 g, 95%); R$_f$ = 0.00 ($CH_2Cl_2$/MeOH 10/1); mp 165–168 ˚C; HPLC purity: 7.4 min, 98.6%; $^1$H NMR (300 MHz, DMSO–$d_6$) $\delta$ 9.21 (br s, $^+$N**H$_2$**), 7.90 (br s, C(O)N**H**H'), 7.34–7.80 (m, 3Ar**H** or 2ArH, C(O)NH**H'**), 7.68 (d, $J$ = 7.6 Hz, 1Ar**H**), 7.55–7.64 (m, 3Ar**H** or 2Ar**H**, C(O)NH**H'**), 7.42–7.55 (m, 2Ar**H**), 4.22 (s, $^+$NH$_2$C**H$_2$**Ar), 3.67 (s, C(O)C**H$_2$**$^+$NH$_2$), 2.34 (s, SC**H$_3$**); $^{13}$C NMR (75 MHz, DMSO–$d_6$) $\delta$ 167.3 (**C**(O)), 142.0, 139.6, 134.3, 131.9, 131.3, 128.0, 127.5, 126.9, 125.9 (Ar**C**), 50.0 (C(O)**C**H$^+$NH$_2$), 47.3 ($^+$NH$_2$**C**H$_2$Ar), 40.1 (S**C**H$_3$) ; LRMS (M + H)$^+$(ESI$^+$) 275.1 [M + H]$^+$ (calcd for $C_{15}H_{15}ClN_2OH^+$ 275.1)

**2-(((4'-Chloro-[1,1'-biphenyl]-4-yl)methyl)amino)acetamide methanesulfonate (17g)**

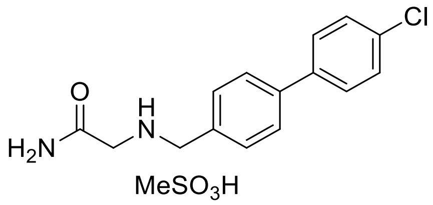


Using method C, to a solution of **10g** (0.15 g, 0.55 mmol), methanesulfonic acid (44.30 µL, 0.68 mmol) in EtOAc (5.46 mL) gave **17g** as a white solid (0.18 g, 89%); R$_f$ = 0.00 ($CH_2Cl_2$/MeOH 10/1); mp 262–266 ˚C; HPLC purity: 7.0 min, 98.4%; $^1$H NMR (300 MHz, DMSO–$d_6$) $\delta$ 9.21 (br s, $^+$N**H$_2$**), 7.85 (br s, C(O)N**H**H'), 7.72–7.77 (m, 4Ar**H**), 7.50–7.63 (m, 4ArH, C(O)NH**H'**), 4.21 (s, $^+$NH$_2$C**H$_2$**Ar), 3.67 (s, C(O)C**H$_2$**$^+$NH$_2$), 2.35 (s, SC**H$_3$**); $^{13}$C NMR (75 MHz, DMSO–$d_6$) $\delta$ 167.3 (**C**(O)), 139.8, 138.6, 133.2, 131.6, 131.3, 129.4, 129.0, 127.3

(Ar**C**), 49.9 (C(O)**C**H$^+$NH$_2$), 47.2 ($^+$NH$_2$**C**H$_2$Ar). The S**C**H$_3$ peak was overlapped with the DMSO signals; LRMS (M + H)$^+$(ESI$^+$) 275.1 [M + H]$^+$ (calcd for $C_{15}H_{15}ClN_2OH^+$ 275.1)

**2-(((3'-(Trifluoromethyl)-[1,1'-biphenyl]-4-yl)methyl)amino)acetamide methanesulfonate (17i)**

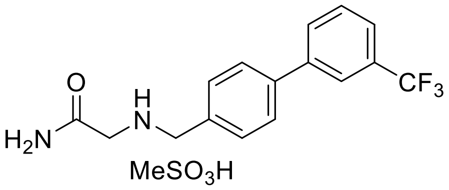


Using method C, to a solution of **10i** (0.15 g, 0.49 mmol), methanesulfonic acid (39.47 µL, 0.61 mmol) in EtOAc (0.97 mL) gave **17i** as a white solid (0.18 g, 91%); R$_f$ = 0.00 ($CH_2Cl_2$/MeOH 10/1); mp 168–169 ˚C; HPLC purity: 7.2 min, 99.9%; $^1$H NMR (300 MHz, DMSO–$d_6$) $\delta$ 9.20 (br s, $^+$N**H$_2$**), 8.00–8.05 (m, 2Ar**H** or 1Ar**H**, C(O)N**H**H'), 7.83–7.90 (m, 3Ar**H** or 2Ar**H**, C(O)N**H**H'), 7.68–7.80 (m, 2Ar**H**), 7.56–7.64 (m, 2ArH, C(O)NH**H'**), 4.23 (s, $^+$NH$_2$C**H$_2$**Ar), 3.67 (s, C(O)C**H$_2$**), 2.32 (s, SC**H$_3$**); $^{13}$C NMR (75 MHz, DMSO–$d_6$) $\delta$ 167.3 (**C**(O)), 140.9, 139.5, 132.1, 131.3, 130.2, 130.3 (q, $J_{C\text{–}F}$ = 31.4 Hz), 127.7, 124.8, 124.6 (q, $J_{C\text{–}F}$ = 270.8 Hz), 124.5 (q, $J_{C\text{–}F}$ = 3.7 Hz) (Ar**C**), 49.9 (C(O)**C**H$^+$NH$_2$), 47.2 ($^+$NH$_2$**C**H$_2$Ar), 40.1 (S**C**H$_3$); LRMS (M + H)$^+$(ESI$^+$) 309.1 [M + H]$^+$ (calcd for $C_{16}H_{15}F_3N_2OH^+$ 309.1)

**2-(((4'-(Trifluoromethylbiphenyl-4-yl)methyl)amino)acetamide methanesulfonate (17j)**

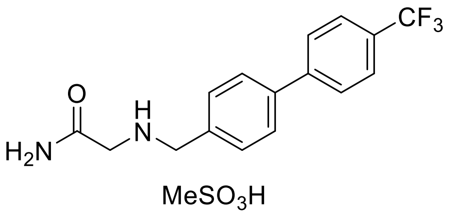


Using method C, to a solution of **10j** (0.23 g, 0.75 mmol), methanesulfonic acid (60.52 µL, 0.93 mmol) in EtOAc (1.50 mL) gave **17j** as a white solid (0.27 g, 90%); R$_f$ = 0.00 ($CH_2Cl_2$/MeOH 10/1); mp 255–256 ˚C; HPLC purity: 7.4 min, 99.9%; $^1$H NMR (300 MHz, DMSO–$d_6$) $\delta$ 9.26 (br s, $^+$N**H$_2$**), 7.91–7.93 (m, 2Ar**H**, C(O)N**H**H'), 7.79–7.82 (m, 4Ar**H**), 7.65 (d, $J$ = 7.2 Hz, 2Ar**H**), 7.58 (br s, C(O)NH**H'**), 4.25 (s, $^+$NH$_2$C**H$_2$**Ar), 3.71 (s, C(O)C**H$_2$**), 2.40

(s, SC**H**$_3$); $^{13}$C NMR (75 MHz, DMSO–$d_6$) $\delta$ 167.3 (**C**(O)), 143.8, 139.6, 132.3, 131.4, 128.6 (q, $J_{C–F}$ = 31.7 Hz), 128.0, 127.7, 126.3 (q, $J_{C–F}$ = 3.7 Hz), 124.8 (q, $J_{C–F}$ = 270.2 Hz) (Ar**C**), 49.9 (C(O)**C**H$^+$NH$_2$), 47.3 ($^+$NH$_2$**C**H$_2$Ar), 40.1 (S**C**H$_3$); LRMS (M + H)$^+$(ESI$^+$) 309.2 [M + H]$^+$ (calcd for $C_{16}H_{15}F_3N_2OH^+$ 309.2)

**2-(((3'-(Trifluoromethoxy)-[1,1'-biphenyl]-4-yl)methyl)amino)acetamide methanesulfonate (17k)**

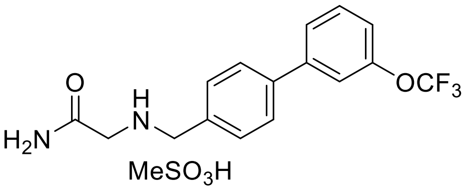


Using method C, to a solution of **10k** (0.02 g, 0.06 mmol), methanesulfonic acid (5.00 µL, 0.08 mmol) in EtOAc (0.12 mL) gave **17k** as a white solid (0.02 g, 66%); R$_f$ = 0.00 ($CH_2Cl_2$/MeOH 10/1); mp 152–156 ˚C; HPLC purity: 7.5 min, 99.2%; $^1$H NMR (300 MHz, CD$_3$OD–$d_4$) $\delta$ 7.76 (d, $J$ = 8.3 Hz, 2Ar**H**), 7.50–7.69 (m, 5Ar**H**), 7.27–7.34 (m, 1Ar**H**), 4.30 (s, $^+$NH$_2$C**H$_2$**Ar), 3.84 (s, C(O)C**H$_2$**$^+$NH$_2$), 2.70 (s, SC**H$_3$**); $^{13}$C NMR (75 MHz, CD$_3$OD–$d_4$) $\delta$ 167.0 (**C**(O)), 149.7, 142.3, 140.8, 130.6, 130.5, 130.4, 127.5, 125.5, 120.6 (q, $J_{C–F}$ = 254.1 Hz), 119.8, 119.2 (Ar**C**), 50.3 (C(O)**C**H$^+$NH$_2$), 38.1 ($^+$NH$_2$**C**H$_2$Ar). The S**C**H$_3$ peak was overlapped with the CD$_3$OD signals; LRMS (M + H)$^+$(ESI$^+$) 325.1 [M + H]$^+$ (calcd for $C_{16}H_{15}F_3N_2O_2H^+$ 325.1)

**2-(((4'-(Trifluoromethoxy)-[1,1'-biphenyl]-4-yl)methyl)amino)acetamide methanesulfonate (17l)**

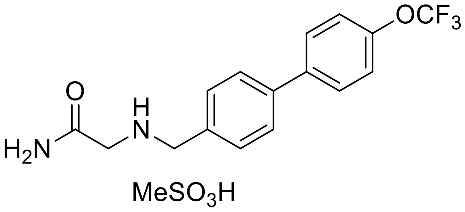


Using method C, to a solution of **10l** (0.30 g, 0.93 mmol), methanesulfonic acid (75.04 µL, 1.16 mmol) in EtOAc (0.93 mL) gave **17l** as a white solid (0.36 g, 94%); R$_f$ = 0.00 ($CH_2Cl_2$/MeOH 10/1); mp 217–220 ˚C; HPLC purity: 7.6 min, >99.9%; $^1$H NMR (300 MHz,

DMSO–*d*$_6$) *δ* 9.20 (br s, $^+$N**H$_2$**), 7.80–7.83 (m, 2Ar**H**, C(O)N**H**H'), 7.77 (d, *J* = 7.8 Hz, 2Ar**H**), 7.156–7.64 (m, 2Ar**H**, C(O)NH**H'**), 7.46 (d, *J* = 8.2 Hz, 2Ar**H**), 4.22 (s, $^+$NH$_2$C**H$_2$**Ar), 3.67 (s, C(O)C**H$_2$**$^+$NH$_2$), 2.34 (s, SC**H$_3$**); $^{13}$C NMR (75 MHz, DMSO–*d*$_6$) *δ* 167.3 (**C**(O)), 148.5 (**C**OCF$_3$), 139.7, 139.1, 131.7, 131.3, 129.1, 127.5, 121.9, 120.6 (q, $J_{C–F}$ = 254.6 Hz) (Ar**C**), 49.9 (C(O)**C**H$^+$NH$_2$), 47.2 ($^+$NH$_2$**C**H$_2$Ar), the S**C**H$_3$ peak was overlapped with the DMSO signals; LRMS (M + H)$^+$(ESI$^+$) 325.1 [M + H]$^+$ (calcd for $C_{16}H_{15}F_3N_2O_2H^+$ 325.1)

**2-(((4'-Methoxy-[1,1'-biphenyl]-4-yl)methyl)amino)acetamide methanesulfonate (17n)**

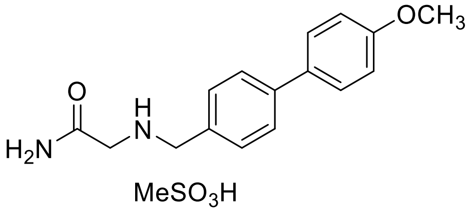


Using method C, to a solution of **10n** (0.08 g, 0.30 mmol), methanesulfonic acid (24.00 µL, 0.37 mmol) in EtOAc (2.96 mL) gave **17n** as a white solid (0.09 g, 81%); R$_f$ = 0.00 (CH$_2$Cl$_2$/MeOH 10/1); mp 243–245 ˚C; HPLC purity: 6.3 min, 99.1%; $^1$H NMR (300 MHz, DMSO–*d*$_6$) *δ* 9.10 (br s, $^+$N**H$_2$**), 7.81 (br s, C(O)N**H**H'), 7.70 (d, *J* = 8.0 Hz, 2Ar**H**), 7.65 (d, *J* = 8.5 Hz, 2Ar**H**), 7.57 (br s, C(O)NH**H'**), 7.53 (d, *J* = 8.0Hz, 2Ar**H**), 7.04 (d, *J* = 8.5 Hz, 2Ar**H**), 4.18 (s, $^+$NH$_2$C**H$_2$**Ar), 3.80 (s, OC**H$_3$**), 3.63 (s, C(O)C**H$_2$**$^+$NH$_2$), 2.30 (s, SC**H$_3$**); $^{13}$C NMR (100 MHz, DMSO–*d*$_6$) *δ* 167.3 (**C**(O)), 159.6 (**C**OCH$_3$), 140.9, 132.1, 131.2, 130.3, 128.3, 126.9, 114.9 (Ar**C**), 55.7 (O**C**H$_3$), 50.0 ($^+$NH$_2$**C**H$_2$Ar), 47.3 (C(O)**C**H$^+$NH$_2$), the S**C**H$_3$ peak was overlapped with the DMSO signals; LRMS (M + H)$^+$(ESI$^+$) 271.1 [M + H]$^+$ (calcd $C_{16}H_{18}N_2O_2H^+$ 271.1)

**(*S*)-2-(([1,1'-Biphenyl]-4-ylmethyl)amino)propanamide methanesulfonate (18a)**[d]

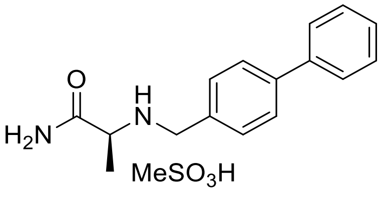


Using method C, to a solution of **11a** (0.15 g, 0.59 mmol), methanesulfonic acid (47.84 µL,

0.74 mmol) in EtOAc (1.18 mL) gave **18a** as a white solid (0.18 g, 86%); R*f* = 0.00 ($CH_2Cl_2$/MeOH 10/1); mp 252–256 ˚C; HPLC purity: 6.2 min, >99.9%; $^1$H NMR (300 MHz, DMSO–*d*6) *δ* 9.19 (br s, $^+$N**H**2), 7.98 (br s, C(O)N**H**H'), 7.75 (d, *J* = 8.0 Hz, 2Ar**H**), 7.70 (d, *J* = 7.8 Hz, 2Ar**H**), 7.65 (br s, C(O)NH**H'**), 7.60 (d, *J* = 8.0 Hz, 2Ar**H**), 7.48 (t, *J* = 7.5 Hz, 2Ar**H**), 7.34–7.43 (m, 1Ar**H**), 4.10–4.20 (m, C**H**2), 3.78–3.94 (m, C**H**), 2.37 (s, SC**H**3), 1.47 (d, *J* = 6.8 Hz, C**H**3); $^{13}$C NMR (75 MHz, DMSO–*d*6) *δ* 170.9 (**C**(O)), 141.2, 139.8, 131.3, 131.2, 129.5, 128.3, 127.3, 127.2 (Ar**C**), 55.0 (**C**H), 48.6 (**C**H2), 40.2 (S**C**H3), 16.4 (**C**H3); LRMS (M + H)$^+$(ESI$^+$) 255.2 [M + H]$^+$ (calcd for $C_{16}H_{18}N_2OH^+$ 255.2)

**(*S*)-2-(((2'-Fluorobiphenyl-4-yl)methyl)amino)propanamide methanesulfonate (18b)**[d]

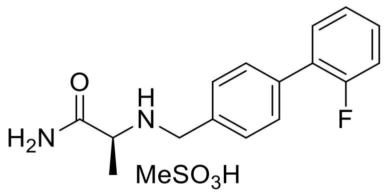


Using method C, to a solution of **11b** (0.10 g, 0.37 mmol), methanesulfonic acid (34.60 µL, 0.53 mmol) in EtOAc (1.50 mL) gave **18b** as a white solid (0.12 g, 90%); R*f* = 0.00 ($CH_2Cl_2$/MeOH 10/1); mp 131–135 ˚C; HPLC purity: 6.4 min, >99.9%; $^1$H NMR (300 MHz, DMSO–*d*6) *δ* 9.17 (br s, $^+$N**H**2), 7.94 (br s, C(O)N**H**H'), 7.30–7.94 (m, 8Ar**H**, C(O)NH**H'**), 4.08–4.25 (m, C**H**2), 3.80 (q, *J* = 6.7 Hz, C**H**), 2.30 (s, SC**H**3), 1.45 (d, *J* = 6.7 Hz, C**H**3); $^{13}$C NMR (75 MHz, DMSO–*d*6) *δ* 170.9 (**C**(O)), 159.5 (d, $J_{C–F}$ = 244.6 Hz), 136.2, 131.7, 131.2 (d, $J_{C–F}$ = 3.1 Hz), 130.8, 130.4 (d, $J_{C–F}$ = 5.4 Hz), 129.5 (d, $J_{C–F}$ = 2.7 Hz), 128.0 (d, $J_{C–F}$ = 13.0 Hz), 125.5 (d, $J_{C–F}$ = 3.4 Hz), 116.6 (d, $J_{C–F}$ = 22.3 Hz) (Ar**C**), 55.1 (**C**H), 48.7 (**C**H2), 16.4 (**C**H3), the S**C**H3 peak was overlapped with the DMSO signals; LRMS (M + H)$^+$(ESI$^+$) 237.1 [M + H]$^+$ (calcd for $C_{16}H_{17}FN_2OH^+$ 237.1)

**(*S*)-2-(((3'-Fluorobiphenyl-4-yl)methyl)amino)propanamide methanesulfonate (18c)**[d]

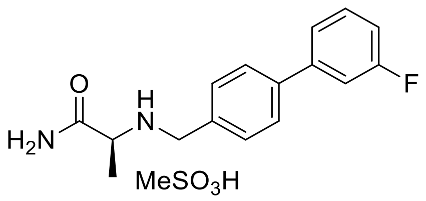

Using method C, to a solution of **11c** (0.30 g, 1.10 mmol), methanesulfonic acid (89.36 µL, 1.38 mmol) in EtOAc (2.50 mL) gave **18c** as a white solid (0.39 g, 97%); $R_f$ = 0.00 ($CH_2Cl_2$/MeOH 10/1); mp 235–238 ˚C; HPLC purity: 6.4 min, >99.9%; $^1$H NMR (300 MHz, DMSO–$d_6$) $\delta$ 9.15 (br s, $^+$N**H**$_2$), 7.92 (br s, C(O)N**H**H'), 7.81 (d, $J$ = 8.3 Hz, 2Ar**H**), 7.68 (br s, C(O)NH**H'**), 7.49–7.60 (m, 5Ar**H**), 7.21–7.27 (m, 1Ar**H**), 4.11–4.20 (m, C**H**$_2$), 3.76 (q, $J$ = 9.3 Hz, C**H**), 2.30 (s, SC**H**$_3$), 1.44 (d, $J$ = 9.3 Hz, C**H**$_3$); $^{13}$C NMR (75 MHz, DMSO–$d_6$) $\delta$ 171.0 (**C**(O)), 163.2 (d, $J_{C-F}$ = 241.9 Hz), 142.3, 139.8, 132.0, 131.5 (d, $J_{C-F}$ = 8.7 Hz), 131.2, 127.5, 123.3, 115.0 (d, $J_{C-F}$ = 21.1 Hz), 113.9 (d, $J_{C-F}$ = 21.9 Hz) (Ar**C**), 55.0 (**C**H), 48.6 (**C**H$_2$), 16.4 (**C**H$_3$), the S**C**H$_3$ peak was overlapped with the DMSO signals; LRMS (M + H)$^+$(ESI$^+$) 273.1 [M + H]$^+$ (calcd for $C_{16}H_{17}FN_2OH^+$ 273.1)

**(*S*)-2-(((4'-Fluorobiphenyl-4-yl)methyl)amino)propanamide methanesulfonate (18d)**[d]

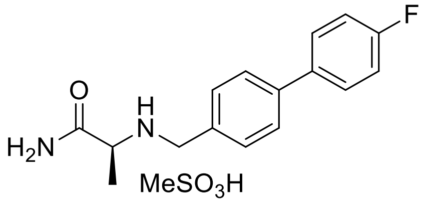


Using method C, to a solution of **11d** (0.20 g, 0.73 mmol), methanesulfonic acid (59.58 µL, 0.92 mmol) in EtOAc (1.46 mL) gave **18d** as a white solid (0.24 g, 88%); $R_f$ = 0.00 ($CH_2Cl_2$/MeOH 10/1); mp 162–164 ˚C; HPLC purity: 6.4 min, 97.6%; $^1$H NMR (300 MHz, DMSO–$d_6$) $\delta$ 9.18 (br s, $^+$N**H**$_2$), 7.95 (br s, C(O)N**H**H'), 7.72–7.77 (m, 4Ar**H**), 7.65 (br s, C(O)NH**H'**), 7.58 (d, $J$ = 8.2 Hz, 2Ar**H**), 7.28–7.34 (m, 2Ar**H**), 4.12–4.15 (m, C**H**$_2$), 3.78–3.84 (m, C**H**), 2.37 (s, SC**H**$_3$), 1.45 (d, $J$ = 6.9 Hz, C**H**$_3$); $^{13}$C NMR (75 MHz, DMSO–$d_6$) $\delta$ 171.0 (**C**(O)), 140.2, 136.3 (d, $J_{C-F}$ = 3.1 Hz), 131.2, 131.1, 129.2 (d, $J_{C-F}$ = 8.2 Hz), 127.3, 116.3 (d, $J_{C-F}$ = 21.2 Hz) (Ar**C**), 54.9 (**C**H), 48.5 (**C**H$_2$), 16.3 (**C**H$_3$), the S**C**H$_3$ peak was overlapped with the DMSO signals; LRMS (M + H)$^+$(ESI$^+$) 273.1 [M + H]$^+$ (calcd for $C_{16}H_{17}FN_2OH^+$ 273.1)

**(*S*)-2-(((2'-Chlorobiphenyl-4-yl)methyl)amino)propanamide methanesulfonate (18e)**[d]

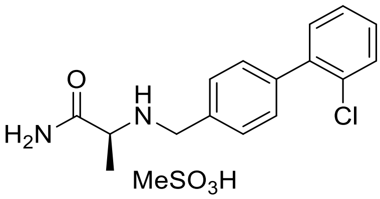


Using method C, to a solution of **11e** (0.10 g, 0.35 mmol), methanesulfonic acid (28.10 µL, 0.43 mmol) in EtOAc (3.46 mL) gave **18e** as a white solid (0.08 g, 63%); $R_f$ = 0.00 ($CH_2Cl_2$/MeOH 10/1); mp 216–220 ˚C; HPLC purity: 6.7 min, 97.8%; $^1$H NMR (300 MHz, DMSO–*d*$_6$) *δ* 9.18 (br s, $^+$N**H**$_2$), 7.96 (br s, C(O)N**H**H'), 7.67 (br s, C(O)NH**H'**), 7.59 (d, *J* = 8.1 Hz, 3Ar**H**), 7.52 (d, *J* = 8.2 Hz, 2Ar**H**), 7.39–7.47 (m, 3Ar**H**), 4.09–4.28 (m, C**H**$_2$), 3.86–3.90 (m, C**H**), 2.30 (s, SC**H**$_3$), 1.47 (d, *J* = 6.9 Hz, C**H**$_3$); $^{13}$C NMR (75 MHz, DMSO–*d*$_6$) *δ* 170.9 (**C**(O)), 139.8, 139.6, 131.9, 131.8, 131.7, 130.4, 130.0, 128.1, 55.2 (**C**H), 48.7 (**C**H$_2$), 16.4 (**C**H$_3$), the S**C**H$_3$ peak was overlapped with the DMSO signals. The remaining peak was not detected and is believed to overlap with the observed signals; LRMS (M + H)$^+$(ESI$^+$) 289.1 [M + H]$^+$ (calcd for $C_{16}H_{17}ClN_2OH^+$ 289.1)

**(*S*)-2-(((3'-Chlorobiphenyl-4-yl)methyl)amino)propanamide methanesulfonate (18f)**[d]

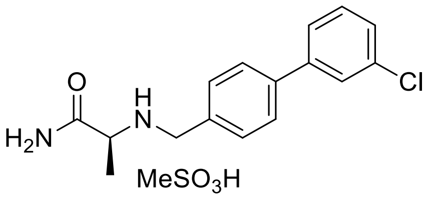


Using method C, to a solution of **11f** (0.15 g, 0.52 mmol), methanesulfonic acid (48.90 µL, 0.75 mmol) in EtOAc (0.52 mL) gave **18f** as a white solid (0.18 g, 90%); $R_f$ = 0.00 ($CH_2Cl_2$/MeOH 10/1); mp 220–223 ˚C; HPLC purity: 6.9 min, 97.3%; $^1$H NMR (400 MHz, DMSO–*d*$_6$) *δ* 9.19 (br s, $^+$N**H**$_2$), 7.96 (br s, C(O)N**H**H'), 7.80 (d, *J* = 8.14 Hz, 2Ar**H**), 7.77 (br s, C(O)NH**H'**), 7.70–7.66 (m, 2Ar**H**), 7.60 (d, *J* = 8.1 Hz, 2Ar**H**), 7.52 (t, *J* = 7.9 Hz, 1Ar**H**), 7.46 (d, *J* = 8.1 Hz, 1Ar**H**), 4.15–4.20 (m, C**H**$_2$), 3.82 (q, *J* = 6.7 Hz, C**H**), 2.34 (s, SC**H**$_3$), 1.47 (d, *J* = 6.7 Hz, C**H**$_3$); $^{13}$C NMR (75 MHz, DMSO–*d*$_6$) *δ* 170.9 (**C**(O)), 141.9, 139.5, 134.3, 132.0, 131.3, 131.2, 128.1, 127.5, 126.9, 125.9 (Ar**C**), 55.1 (**C**H), 48.6 (**C**H$_2$), 16.4 (**C**H$_3$), the S**C**H$_3$ peak was overlapped with the DMSO signals; LRMS (M + H)$^+$(ESI$^+$) 289.1 [M + H]$^+$ (calcd for $C_{16}H_{17}ClN_2OH^+$ 289.1)

**(*S*)-2-(((4'-Chlorobiphenyl-4-yl)methyl)amino)propanamide methanesulfonate (18g)**[d]

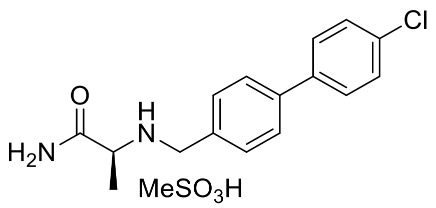


Using method C, to a solution of **11g** (0.15 g, 0.52 mmol), methanesulfonic acid (42.14 µL, 0.65 mmol) in EtOAc (1.04 mL) gave **18g** as a white solid (0.17 g, 84%); $R_f$ = 0.00 ($CH_2Cl_2$/MeOH 10/1); mp 220–223 ˚C; HPLC purity: 7.0 min, 98.2%; $^1$H NMR (300 MHz, DMSO–$d_6$) $\delta$ 9.17 (br s, $^+$N**H2**), 7.94 (br s, C(O)N**H**H'), 7.73–7.78 (m, 4Ar**H**), 7.66 (br s, C(O)NH**H'**), 7.53–7.60 (m, 4Ar**H**), 4.10–4.20 (m, C**H2**), 3.76–3.82 (m, C**H**), 2.32 (s, SC**H3**), 1.45 (d, *J* = 6.9 Hz, C**H3**); $^{13}$C NMR (100 MHz, DMSO–$d_6$) $\delta$ 170.9 (**C**(O)), 139.9, 138.6, 133.2, 131.7, 131.2, 129.4, 129.0, 127.3 (Ar**C**), 54.9, (**C**H), 48.5 (**C**H2), 16.3 (**C**H3), the S**C**H3 peak was overlapped with the DMSO signals; LRMS (M + H)$^+$(ESI$^+$) 289.1 [M + H]$^+$ (calcd for $C_{16}H_{17}ClN_2OH^+$ 289.1)

**(*S*)-2-(((2'-Trifluoromethylbiphenyl-4-yl)methyl)amino)propanamide methanesulfonate (18h)**[d]

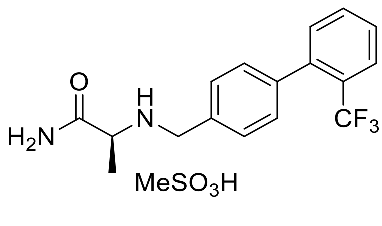


Using method C, to a solution of **11h** (0.12 g, 0.37 mmol), methanesulfonic acid (30.20 µL, 0.47 mmol) in EtOAc (3.72 mL) gave **18h** as a white solid (0.14 g, 87%); $R_f$ = 0.00 ($CH_2Cl_2$/MeOH 10/1); mp 224–227 ˚C; HPLC purity: 7.0 min, 95.8%; $^1$H NMR (300 MHz, DMSO–$d_6$) $\delta$ 9.20 (br s, $^+$N**H2**), 7.94 (br s, C(O)N**H**H'), 7.85 (d, *J* = 7.8 Hz, 1Ar**H**), 7.75 (t, *J* = 7.4 Hz, 1Ar**H**), 7.61–7.67 (m, 2Ar**H**), 7.57 (d, *J* = 7.2 Hz, 1Ar**H**), 7.39–7.41 (m, 2Ar**H**, C(O)NH**H'**), 4.11–4.22 (m, C**H2**), 3.86–3.88 (m, C**H**), 2.32 (s, SC**H3**), 1.47 (d, *J* = 6.7 Hz, C**H3**); $^{13}$C NMR (75 MHz, DMSO–$d_6$) $\delta$ 170.9 (**C**(O)), 140.5, 140.4, 132.8, 132.5, 131.9, 130.1, 129.4, 128.7, 127.3 (q, $J_{C–F}$ = 29.2 Hz), 126.5 (q, $J_{C–F}$ = 5.2 Hz), 124.6 (q, $J_{C–F}$ = 270.5 Hz),

55.4, (**C**H), 48.8 (**C**$H_2$), 40.2 (S**C**$H_3$), 16.4 (**C**$H_3$); LRMS (M + H)$^+$(ESI$^+$) 323.1 [M + H]$^+$ (calcd for $C_{17}H_{17}F_3N_2OH^+$ 323.1)

**(*S*)-2-(((3'-Trifluoromethylbiphenyl-4-yl)methyl)amino)propanamide methanesulfonate (18i)**[*d*]

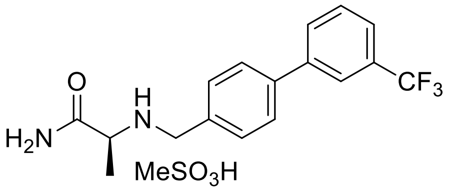


Using method C, to a solution of **11i** (0.09 g, 0.28 mmol), methanesulfonic acid (25.94 μL, 0.40 mmol) in EtOAc (1.10 mL) gave **18i** as a white solid (0.10 g, 92%); R$_f$ = 0.00 ($CH_2Cl_2$/MeOH 10/1); mp 215–217 ˚C; HPLC purity: 7.3 min, >99.9%; $^1$H NMR (400 MHz, DMSO–$d_6$) $\delta$ 9.16 (br s, $^+$N**H$_2$**), 7.98–8.02 (m, 2Ar**H**), 7.90 (br s, C(O)N**H**H'), 7.84 (d, $J$ = 8.1 Hz, 2Ar**H**), 7.69–7.76 (m, 2Ar**H**), 7.65 (br s, C(O)NH**H'**), 7.59 (d, $J$ = 8.1 Hz, 2Ar**H**), 4.14 (m, C**H$_2$**), 3.69–3.84 (m, C**H**), 2.27 (S, SC**H$_3$**), 1.43 (d, $J$ = 6.9 Hz, C**H$_3$**); $^{13}$C NMR (75 MHz, DMSO–$d_6$) $\delta$ 171.0 (**C**(O)), 140.9, 139.5, 132.2, 131.3, 130.6, 130.3 (q, $J_{C–F}$ = 31.4 Hz), 127.6, 124.8, 124.6 (q, $J_{C–F}$ = 270.7 Hz), 123.5 (q, $J_{C–F}$ = 3.6 Hz) (Ar**C**), 55.2 (**C**H), 48.6 (**C**$H_2$), 16.4 (**C**$H_3$). The S**C**$H_3$ peak was overlapped with the DMSO signals. The remaining peak was not detected and is believed to overlap with the observed signals; LRMS (M + H)$^+$(ESI$^+$) 323.1 [M + H]$^+$ (calcd for $C_{17}H_{17}F_3N_2OH^+$ 323.1)

**(*S*)-2-(((4'-Trifluoromethylbiphenyl-4-yl)methyl)amino)propanamide methanesulfonate (18j)**[*d*]

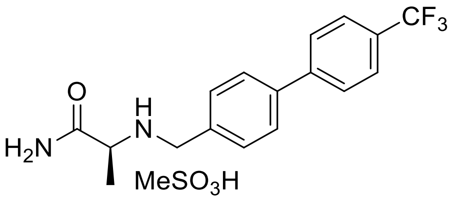


Using method C, to a solution of **11j** (1.12 g, 3.63 mmol), methanesulfonic acid (295.00 μL, 1.25 mmol) in EtOAc (7.26 mL) gave **18j** as a white solid (1.26 g, 83%); R$_f$ = 0.00

($CH_2Cl_2$/MeOH 10/1); mp 241–243 °C; HPLC purity: 7.4 min, 99.7%; $^1$H NMR (300 MHz, DMSO–$d_6$) $\delta$ 9.20 (br s, $^+$N**H**$_2$), 7.94 (d, $J$ = 7.7 Hz, 2Ar**H**, C(O)N**H**H'), 7.83–7.86 (m, 4Ar**H**), 7.63–7.66 (m, 2Ar**H**, C(O)NH**H'**), 4.12–4.23 (m, C**H**$_2$), 3.82 (q, $J$ = 6.4 Hz, C**H**), 2.33 (s, SC**H**$_3$), 1.46 (d, $J$ = 6.4 Hz, C**H**$_3$); $^{13}$C NMR (75 MHz, DMSO–$d_6$) $\delta$ 170.9 (**C**(O)), 143.8, 139.6, 132.4, 131.3, 128.6 (q, $J_{C–F}$ = 31.9 Hz), 128.0, 127.8, 126.3 (q, $J_{C–F}$ = 3.8 Hz), 124.8 (q, $J_{C–F}$ = 270.3 Hz) (Ar**C**), 55.0 (C(O)**C**H$^+$NH$_2$), 48.5 (**C**H$_2$), 40.2 (S**C**H$_3$), 16.4 (**C**H$_3$); LRMS (M + H)$^+$(ESI$^+$) 323.1 [M + H]$^+$ (calcd for $C_{17}H_{17}F_3N_2OH^+$ 323.1)

**(*S*)-2-(((3'-Trifluoromethoxybiphenyl-4-yl)methyl)amino)propanamide methanesulfonate (18k)**[d]

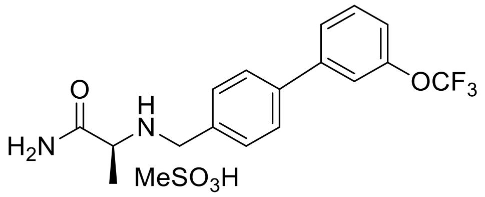


Using method C, to a solution of **11k** (0.25 g, 0.74 mmol), methanesulfonic acid (59.94 µL, 0.92 mmol) in EtOAc (1.48 mL) gave **18k** as a white solid (0.29 g, 90%); R$_f$ = 0.00 ($CH_2Cl_2$/MeOH 10/1); mp 209–212 °C; HPLC purity: 7.5 min, 98.8%; $^1$H NMR (300 MHz, DMSO–$d_6$) $\delta$ 9.16 (br s, $^+$N**H**$_2$), 7.92 (br s, C(O)N**H**H'), 7.83 (d, $J$ = 8.2 Hz, 2Ar**H**), 7.77 (d, $J$ = 8.2 Hz, 1Ar**H**), 7.59–7.69 (m, 4Ar**H**, C(O)NH**H'**), 7.39–7.42 (m, 1Ar**H**), 4.08–4.23 (m, C**H**$_2$), 3.77 (q, $J$ = 7.0 Hz, C**H**), 2.30 (s, SC**H**$_3$), 1.44 (d, $J$ = 7.0 Hz, C**H**$_3$); $^{13}$C NMR (100 MHz, DMSO–$d_6$) $\delta$ 170.9 (**C**(O)), 149.5, 142.2, 139.4, 132.2, 131.5, 131.2, 127.6, 126.3, 120.6 (q, $J_{C–F}$ = 254.7 Hz), 120.5, 119.7 (Ar**C**), 55.0, (**C**H), 48.5 (**C**H$_2$), 16.4 (**C**H$_3$), the S**C**H$_3$ peak was overlapped with the DMSO signals; LRMS (M + H)$^+$(ESI$^+$) 339.1 [M + H]$^+$ (calcd for $C_{17}H_{17}F_3N_2O_2H^+$ 339.1)

**(*S*)-2-(((4'-Trifluoromethoxybiphenyl-4-yl)methyl)amino)propanamide methanesulfonate (18l)**[d]

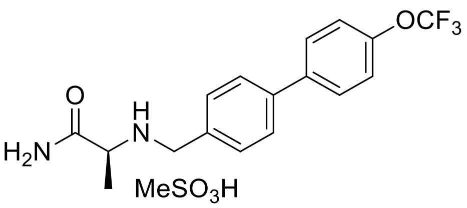


Using method C, to a solution of **11l** (0.10 g, 0.30 mmol), methanesulfonic acid (23.98 μL, 0.37 mmol) in EtOAc (0.59 mL) gave **18l** as a white solid (0.12 g, 92%); $R_f$ = 0.00 ($CH_2Cl_2$/MeOH 10/10); mp 256–259 °C; HPLC purity: 7.6 min, 99.8%; $^1$H NMR (400 MHz, DMSO–$d_6$) $\delta$ 9.19 (br s, $^+$N**H**$_2$), 7.96 (br s, C(O)N**H**H'), 7.84 (d, $J$ = 8.7 Hz, 2Ar**H**), 7.78 (d, $J$ = 8.2 Hz, 2Ar**H**), 7.67 (br s, C(O)NH**H'**), 7.61 (d, $J$ = 8.1 Hz, 2Ar**H**), 7.48 (d, $J$ = 8.2 Hz, 2Ar**H**), 4.16–4.20 (m, C**H**$_2$), 3.82–3.83 (m, C**H**), 2.34 (s, SC**H**$_3$), 1.46 (d, $J$ = 7.0 Hz, C**H**$_3$); $^{13}$C NMR (100 MHz, DMSO–$d_6$) $\delta$ 170.9 (**C**(O)), 148.5, 139.7, 139.1, 131.8, 129.1, 127.5, 122.0, 120.6 (q, $J_{C-F}$ = 254.7 Hz) (Ar**C**), 55.0 (**C**H), 48.6 (**C**H$_2$), 16.4 (**C**H$_3$), the S**C**H$_3$ peak was overlapped with the DMSO signals; LRMS $(M + H)^+(ESI^+)$ 339.1 $[M + H]^+$ (calcd for $C_{17}H_{17}F_3N_2O_2H^+$ 339.1)

**(*S*)-2-(((4'-Methoxybiphenyl-4-yl)methyl)amino)propanamide methanesulfonate (18n)**

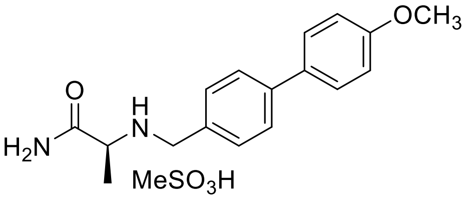


Using method C, to a solution of **11n (**0.09 g, 0.31 mmol), methanesulfonic acid (24.82 μL, 0.38 mmol) in EtOAc (0.61 mL) gave **18n** as a white solid (0.11 g, 94%); $R_f$ = 0.00 ($CH_2Cl_2$/MeOH 10/1); mp 231–233 °C; HPLC purity: 6.3 min, 99.6%; $^1$H NMR (300 MHz, DMSO–$d_6$) $\delta$ 9.14 (br s, $^+$N**H**$_2$), 7.92 (br s, C(O)N**H**H'), 7.64–7.72 (m, 4Ar**H**, C(O)NH**H'**), 7.54 (d, $J$ = 8.3 Hz, 2Ar**H**), 7.04 (d, $J$ = 8.8 Hz, 2Ar**H**), 4.06–4.22 (m, C**H**$_2$), 3.72–3.89 (m, OC**H**$_3$, C**H**), 2.31 (s, SC**H**$_3$), 1.44 (d, $J$ = 7.0 Hz, C**H**$_3$); $^{13}$C NMR (75 MHz, DMSO–$d_6$) $\delta$ 170.9 (**C**(O)), 159.7 (**C**OCH$_3$), 140.9, 132.1, 131.1, 130.4, 128.3, 126.8, 114.9 (Ar**C**), 55.7 (**C**H), 54.8 (O**C**H$_3$), 48.6 (**C**H$_2$), 40.2 (S**C**H$_3$), 16.4 (**C**H$_3$); LRMS $(M + H)^+(ESI^+)$ 285.2 $[M + H]^+$ (calcd for $C_{17}H_{20}N_2O_2H^+$ 285.2)

**(*S*)-2-(((4'-Isobutyl-[1,1'-biphenyl]-4-yl)methyl)amino)propanamide methanesulfonate (18o)**

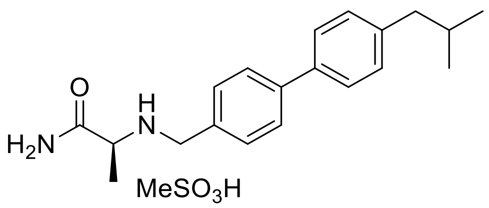


Using method C, to a solution of **11o** (0.25 g, 0.81 mmol), methanesulfonic acid (65.33 µL, 1.00 mmol) in EtOAc (8.05 mL) gave **18o** as a white solid (0.28 g, 86%); $R_f$ = 0.00 ($CH_2Cl_2$/MeOH 10/1); mp 253–256 ˚C; HPLC purity: 8.4 min, >99.9%; $^1$H NMR (300 MHz, DMSO–$d_6$) $\delta$ 9.15 (br s, $^+$N**H**$_2$), 7.94 (br s, C(O)N**H**H'), 7.74 (d, *J* = 8.2 Hz, 2Ar**H**), 7.53–7.68 (m, 4Ar**H**, C(O)NH**H'**), 7.26 (d, *J* =8.1Hz, 2Ar**H**), 4.03–4.26 (m, $^+NH_2C\mathbf{H_2}Ph$), 3.82 (q, *J* = 6.8 Hz, C**H**), 2.50 (d, *J* = 7.0 Hz, C$\mathbf{H_2}$CH), 2.34 (s, SC$\mathbf{H_3}$), 1.80–1.94 (m, $CH_2$C**H**), 1.46 (d, *J* = 6.9 Hz, CHC$\mathbf{H_3}$), 0.89 (d, *J* = 6.6 Hz, CH(C$\mathbf{H_3}$)$_2$); $^{13}$C NMR (75 MHz, DMSO–$d_6$) $\delta$ 170.9 (**C**(O)), 141.3, 141.1, 137.2, 131.1, 130.9, 130.1, 127.1, 126.9 (Ar**C**), 55.0 (**C**H), 48.7 (**C**$H_2$), 40.9 (**C**H$CH_2$), 40.2 (S**C**$H_3$), 30.1 (CH**C**$H_2$), 22.6 (CH(**C**$H_3$)$_2$), 16.4 (**C**$H_3$); LRMS (M + H)$^+$(ESI$^+$) 311.2 [M + H]$^+$ (calcd for $C_{20}H_{26}N_2OH^+$ 311.2).

**(*R*)-2-(((4'-(Trifluoromethylbiphenyl-4-yl)methyl)amino)propanamide methanesulfonate (19j)**[d]

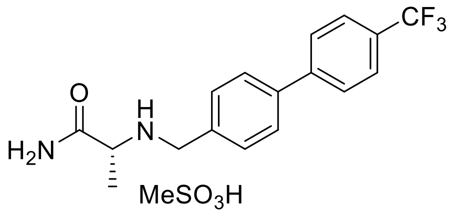


Using method C, to a solution of **12j** (0.17 g, 0.53 mmol), methanesulfonic acid (42.80 µL, 0.66 mmol) in EtOAc (1.05 mL) gave **19j** as a white solid (0.19 g, 87%); $R_f$ = 0.00 ($CH_2Cl_2$/MeOH 10/1); mp 241–244 ˚C; HPLC purity: 7.4 min, 99.8%; $^1$H NMR (300 MHz, DMSO–$d_6$) $\delta$ 9.18 (br s, $^+$N**H**$_2$), 7.93–7.95 (m, 2Ar**H**, C(O)N**H**H'), 7.84 (d, *J* = 7.9 Hz, 4Ar**H**), 7.62–7.66 (m, 2Ar**H**, C(O)NH**H'**), 4.12–4.22 (m, C$\mathbf{H_2}$), 3.80 (q, *J* = 6.5 Hz, C**H**), 2.31 (s, SC$\mathbf{H_3}$), 1.45 (d, *J* = 6.5 Hz, C$\mathbf{H_3}$); $^{13}$C NMR (75 MHz, DMSO–$d_6$) $\delta$ 170.9 (**C**(O)), 143.8, 139.5, 132.4, 131.3, 128.6 (q, $J_{C\text{–}F}$ = 31.7 Hz) (Ar**C**), 55.1 (**C**H), 48.6 (**C**$H_2$), 40.2 (S**C**$H_3$), 16.4 (**C**$H_3$);

LRMS $(M + H)^+(ESI^+)$ 323.1 $[M + H]^+$ (calcd for $C_{17}H_{17}F_3N_2OH^+$ 323.1)

**(*S*)-2-(((4'-(Trifluoromethyl)-[1,1'-biphenyl]-4-yl)methyl)amino)butanamide methanesulfonate (20j)**

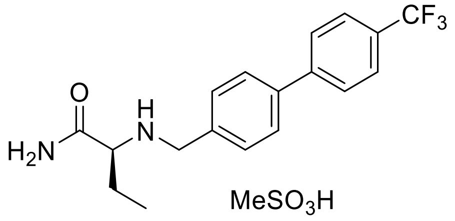


Using method C, to a solution of **13j** (0.18 g, 0.54 mmol), methanesulfonic acid (43.40 µL, 0.67 mmol) in EtOAc (1.07 mL) gave **20j** as a white solid (0.21 g, 91%); $R_f$ = 0.00 ($CH_2Cl_2$/MeOH 10/1); mp 253–256 ˚C; HPLC purity: 7.5 min, >99.9%; $^1$H NMR (300 MHz, DMSO–$d_6$) $\delta$ 9.23 (br s, $^+$N**H**$_2$), 7.95 (br s, C(O)N**H**H'), 7.93 (d, $J$ = 8.1 Hz, 2Ar**H**), 7.82 (d, $J$ = 7.9 Hz, 4Ar**H**), 7.77 (br s, C(O)NH**H'**), 7.64 (d, $J$ = 8.1 Hz, 2Ar**H**), 4.05–4.28 (m, C**H**$_2$), 3.68–3.86 (m, C**H**), 2.30 (s, SC**H**$_3$), 1.52–2.03 (m, CHC**H**$_2$), 0.92 (t, $J$ = 7.4 Hz, C**H**$_3$); $^{13}$C NMR (75 MHz, DMSO–$d_6$) $\delta$ 169.5 (**C**(O)), 143.8, 139.6, 132.2, 131.5, 128.6 (q, $J_{C–F}$ = 31.7 Hz), 128.0, 127.7, 126.3 (q, $J_{C–F}$ = 3.7 Hz), 124.8 (q, $J_{C–F}$ = 270.4 Hz) (Ar**C**), 60.2 (C(O)**C**H$^+$NH$_2$), 49.0 ($^+$NH$_2$**C**H$_2$Ar), 23.4 (CH**C**H$_2$), 9.4 (**C**H$_3$), the S**C**H$_3$ peak was overlapped with the DMSO signals; LRMS $(M + H)^+(ESI^+)$ 337.2 $[M + H]^+$ (calcd for $C_{18}H_{19}F_3N_2OH^+$ 337.2)

**(*S*)-2-(((4'-(Trifluoromethoxy)-[1,1'-biphenyl]-4-yl)methyl)amino)butanamide methanesulfonate (20l)**

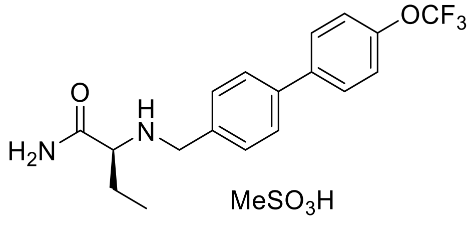


Using method C, to a solution of **13l** (0.20 g, 0.57 mmol), methanesulfonic acid (46.04 µL, 0.71 mmol) in EtOAc (1.13 mL) gave **20l** as a white solid (0.14 g, 57%); $R_f$ = 0.00

($CH_2Cl_2$/MeOH 10/1); mp 253–257 ˚C; HPLC purity: 7.8 min, >99.9%; $^{1}$H NMR (300 MHz, DMSO–$d_6$) $\delta$ 9.20 (br s, $^{+}$N**H2**), 8.05 (br s, C(O)N**H**H'), 7.83 (d, $J$ = 8.2 Hz, 2Ar**H**), 7.71–7.78 (m, 2Ar**H**, C(O)NH**H'**), 7.61 (d, $J$ = 7.7 Hz, 2Ar**H**), 7.46 (d, $J$ = 8.0 Hz, 2Ar**H**), 4.05–4.23 (m, $^{+}NH_2$C**H2**Ar), 3.71–3.79 (m, C**H**), 2.38 (s, SC**H3**), 1.71–2.01 (m, CHC**H2**), 0.92 (t, $J$ = 7.1 Hz, C**H3**); $^{13}$C NMR (75 MHz, DMSO–$d_6$) $\delta$ 169.5 (**C**(O)), 148.5(**C**O$CF_3$), 139.7, 139.1, 131.6, 131.4, 129.1, 127.4, 121.9, 120.6 (q, $J_{C–F}$ = 254.6 Hz) (Ar**C**), 60.2 (C(O)**C**H$^{+}NH_2$), 49.1 ($^{+}NH_2$**C**$H_2$Ar), 40.2 (S**C**$H_3$), 23.4 (CH**C**$H_2$), 9.4 (**C**$H_3$); LRMS (M + H)$^{+}$(ESI$^{+}$) 353.2 [M + H]$^{+}$ (calcd for $C_{18}H_{19}F_3N_2O_2H^+$ 353.2)

**(*S*)-3-Methyl-2-(((3'-(trifluoromethyl)-[1,1'-biphenyl]-4-yl)methyl)amino)butanamide methanesulfonate (21i)**

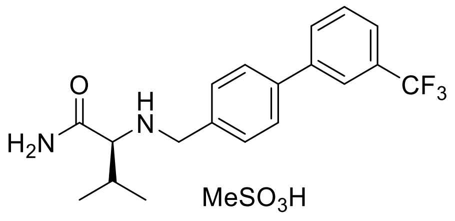


Using method C, to a solution of **14i** (0.20 g, 0.57 mmol), methanesulfonic acid (46.30 µL, 0.71 mmol) in EtOAc (1.14 mL) gave **21i** as a white solid (0.24 g, 95%); R$_f$ = 0.00 ($CH_2Cl_2$/MeOH 10/1); mp 274–278 ˚C; HPLC purity: 9.9 min, 99.7% ; $^{1}$H NMR (300 MHz, DMSO–$d_6$) $\delta$ 9.20 (br s, $^{+}$N**H**H'), 8.99 (br s, $^{+}$NH**H'**), 7.98–8.07 (m, 2Ar**H**), 7.94 (br s, C(O)N**H**H'), 7.86 (d, $J$ = 7.7 Hz, 2Ar**H**), 7.79 (br s, C(O)NH**H'**), 7.69–7.76 (m, 2Ar**H**), 7.64 (d, $J$ = 7.7 Hz, 2Ar**H**), 4.02–4.22 (m, C**H2**), 3.51–3.65 (m, C(O)C**H**), 2.34 (s, SC**H3**), 2.11–2.29 (m, CHC**H**$(CH_3)_2$), 0.88–1.09 (m, 2C**H3**); $^{13}$C NMR (75 MHz, DMSO–$d_6$) $\delta$ 168.4 (**C**(O)), 140.8, 139.5, 131.6, 131.5, 131.3, 130.7, 130.3 (q, $J_{C–F}$ = 31.5 Hz), 127.6, 124.8, 124.7 (q, $J_{C–F}$ = 270.8 Hz), 123.5 (q, $J_{C–F}$ = 3.6 Hz) (Ar**C**), 64.1 (C(O)**C**H$^{+}NH_2$), 49.6 ($^{+}NH_2$**C**$H_2$Ar), 29.3 (C(O)CH**C**H), 19.1 (CH(**C**$H_3$) (C'H3)), 18.2 (CH(CH3) (**C'**$H_3$)), the S**C**$H_3$ peak was overlapped with the DMSO signals; LRMS (M + H)$^{+}$(ESI$^{+}$) 351.2 [M + H]$^{+}$ (calcd for $C_{16}H_{15}F_3N_2OH^+$ 351.2)

**(*R*)-3-Methyl-2-(((4'-(trifluoromethylbiphenyl-4-yl)methyl)amino)butanamide methanesulfonate (21j)**

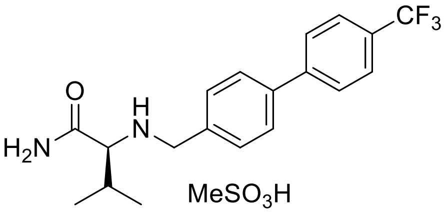


Using method C, to a solution of **14j** (0.08 g, 0.23 mmol), methanesulfonic acid (18.52 µL, 0.29 mmol) in EtOAc (0.46 mL) gave **21j** as a white solid (0.08 g, 74%); R$_f$ = 0.00 ($CH_2Cl_2$/MeOH 10/1); mp 266–267 ˚C; HPLC purity: 7.7 min, 99.4%; $^1$H NMR (300 MHz, DMSO–$d_6$) $\delta$ 9.20 (br s, $^+$N**H**H'), 8.95 (br s, $^+$NH**H'**), 7.78–7.96 (m, 6Ar**H**, C(O)N**H$_2$**), 7.60–7.65 (m, 2Ar**H**), 4.02–4.18 (m, C**H$_2$**), 3.47–3.69 (m, C(O)C**H**), 2.30 (s, SC**H$_3$**), 2.16–2.22 (m, CHC**H**($CH_3$)$_2$), 0.92–1.00(m, 2C**H$_3$**); $^{13}$C NMR (75 MHz, DMSO–$d_6$) $\delta$ 168.4 (**C**(O)), 143.8, 139.6, 131.8, 131.7, 128.6 (q, $J_{C–F}$ = 31.8 Hz), 126.3 (q, $J_{C–F}$ = 3.7 Hz), 124.8 (q, $J_{C–F}$ = 270.2 Hz), 64.1 (C(O)**C**H$^+$NH$_2$), 49.7 ($^+$NH$_2$**C**H$_2$Ar), 40.2 (S**C**H$_3$), 29.3 (C(O)CH**C**H), 19.1 (CH(**C**H$_3$) (C'H$_3$)), 18.1 (CH(CH$_3$) (**C'**H$_3$)); LRMS (M + H)$^+$(ESI$^+$) 351.2 [M + H]$^+$ (calcd for $C_{19}H_{21}F_3N_2OH^+$ 351.2)

**(*S*)-2-(((3'-Fluoro-[1,1'-biphenyl]-4-yl)methyl)amino)-4-methylpentanamide methanesulfonate (22c)**

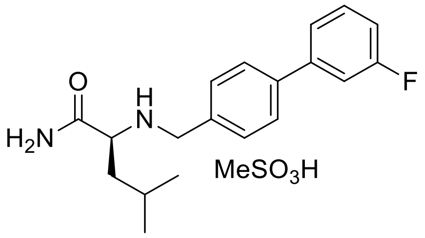


Using method C, to a solution of **15c** (0.10 g, 0.32 mmol), methanesulfonic acid (25.80 µL, 0.40 mmol) in EtOAc (0.64 mL) gave **22c** as a white solid (0.12 g, 90%); R$_f$ = 0.00 ($CH_2Cl_2$/MeOH 10/1); mp 264–265 ˚C; HPLC purity: 7.1 min, >99.9%; $^1$H NMR (300 MHz, DMSO–$d_6$) $\delta$ 9.30 (br s, $^+$N**H**H'), 9.18 (br s, $^+$NH**H'**), 8.05 (br s, C(O)N**H**H'), 7.81 (d, $J$ = 8.01 Hz, 2Ar**H**), 7.76 (br s, C(O)NH**H'**), 7.49–7.59 (m, 5Ar**H**), 7.20–7.26 (m, 1Ar**H**), 4.06–4.18 (m, $^+$NH$_2$C**H$_2$**Ar), 3.60–3.75 (m, C(O)C**H**), 2.31 (s, SC**H$_3$**), 1.58–1.78 (m, C**H**, CHC**H$_2$**), 0.92 (d, $J$ = 5.43 Hz, CH(C**H$_3$**) (CH'$_3$)), 0.88 (d, $J$ = 5.43 Hz, CH(CH$_3$) (C**H'$_3$**)); $^{13}$C NMR (75 MHz, DMSO–$d_6$) $\delta$ 170.0 (**C**(O)), 163.2 (d, $J_{C–F}$ = 241.8 Hz), 142.2 (d, $J_{C–F}$ = 7.7 Hz), 139.7 (d, $J_{C–F}$ = 2.1 Hz), 131.7, 131.4 (d, $J_{C–F}$ = 8.5 Hz), 131.3, 123.2 (d, $J_{C–F}$ = 2.5 Hz), 115.0 (d, $J_{C–F}$ = 20.8

Hz), 113.8 (d, $J_{C\text{–}F}$ = 22.0 Hz) (Ar**C**), 58.5 (C(O)**C**H$^+$NH$_2$), 49.0 ($^+$NH$_2$**C**H$_2$Ar), 40.2 (S**C**H$_3$), 39.2 (**C**H(CH$_3$)$_2$), 24.4 (CH**C**H$_2$), 23.5 (CH(**C**H$_3$) (C'H$_3$)), 22.3 (CH(CH$_3$) (**C'**H$_3$)); LRMS (M + H)$^+$(ESI$^+$) 315.2 [M + H]$^+$ (calcd for $C_{19}H_{23}FN_2OH^+$ 315.2)

**(*S*)-4-Methyl-2-(((3'-(trifluoromethyl)-[1,1'-biphenyl]-4-yl)methyl)amino)pentanamide methanesulfonate (22i)**

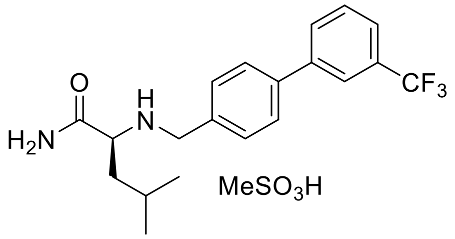


Using method C, to a solution of **15i** (0.15 g, 0.41 mmol), methanesulfonic acid (33.40 μL, 0.52 mmol) in EtOAc (0.82 mL) gave **22i** as a white solid (0.16 g, 86%); R$_f$ = 0.00 ($CH_2Cl_2$/MeOH 10/1); mp 239–243 ˚C; HPLC purity: 8.0 min, >99.9%; $^1$H NMR (300 MHz, DMSO–$d_6$) $\delta$ 9.30 (br s, $^+$N**H**H'), 9.11 (br s, $^+$NH**H'**), 8.10 (br s, C(O)N**H**H'), 7.98–8.07 (m, 2Ar**H** or 1Ar**H**, C(O)NH**H'**), 7.83–7.90 (m, 2Ar**H**), 7.71–7.81 (m, 3Ar**H** or 2Ar**H**, C(O)NH**H'**), 7.58–7.66 (m, 2Ar**H**), 4.02–4.21 (m, $^+$NH$_2$C**H$_2$**Ar), 3.65–3.71 (m, C(O)C**H**), 2.35 (s, SC**H$_3$**), 1.55–1.78 (m, C**H**(CH$_2$), CHC**H$_2$**), 0.78–1.03 (m, CH(C**H$_3$**)$_2$); $^{13}$C NMR (75 MHz, DMSO–$d_6$) $\delta$ 170.0 (**C**(O)), 140.8, 139.5, 132.0, 131.4, 131.3, 130.6, 130.3 (q, $J_{C\text{–}F}$ = 31.5 Hz), 127.6, 124.8, 124.6 (q, $J_{C\text{–}F}$ = 270.7 Hz), 123.5 (q, $J_{C\text{–}F}$ = 3.7 Hz) (Ar**C**), 58.5 ($^+$NH$_2$**C**H$_2$Ar), 49.0 (S**C**H$_3$), 39.2 (**C**H(CH$_3$)$_2$), 24.5 (CH**C**H$_2$), 23.5 (CH(**C**H$_3$) (CH'$_3$)), 22.2 (CH(CH$_3$) (**C'**H$_3$), the S**C**H$_3$ peak was overlapped with the DMSO signals; LRMS (M + H)$^+$(ESI$^+$) 365.2 [M + H]$^+$ (calcd for $C_{20}H_{23}F_3N_2OH^+$ 365.2)

**(*R*)-4-Methyl-2-(((4'-(trifluoromethylbiphenyl-4-yl)methyl)amino)pentanamide methanesulfonate (22j)**

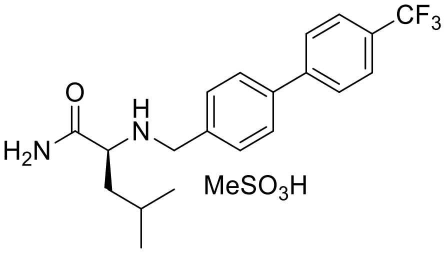


Using method C, to a solution of **15j** (0.13 g, 0.36 mmol), methanesulfonic acid (42.86 μL, 0.45 mmol) in EtOAc (0.71 mL) gave **22j** as a white solid (0.13 g, 77%); $R_f$ = 0.00 ($CH_2Cl_2$/MeOH 10/1); mp 267–270 ˚C; HPLC purity: 10.5 min, 98.6%; $^1$H NMR (300 MHz, DMSO–*d*6) *δ* 9.30 (br s, $^+$N**H**H'), 9.16 (br s, $^+$NH**H'**), 8.13 (br s, C(O)N**H**H'), 7.94 (d, *J* = 7.85 Hz, 2Ar**H**), 7.84 (d, *J* = 7.80 Hz, 4Ar**H**), 7.79 (br s, C(O)NH**H'**), 7.63 (d, *J* = 7.85 Hz, 2Ar**H**), 4.05–4.25 (m, $^+NH_2C\mathbf{H_2}Ar$), 3.70–3.83 (m, C(O)C**H**), 2.35 (s, SC**H3**), 1.59–1.79 (m, C**H**($CH_2$), CHC**H2**), 0.80–1.03 (m, CH(C**H3**)2); $^{13}$C NMR (75 MHz, DMSO–*d*6) *δ* 169.9 (**C**(O)), 143.8, 139.6, 132.2, 131.4, 128.6 (q, $J_{C–F}$ = 31.9 Hz), 128.4, 127.7, 126.3 (q, $J_{C–F}$ = 3.7 Hz), 124.8 (q, $J_{C–F}$ = 270.3 Hz) (Ar**C**), 58.4 (C(O)**C**H$^+NH_2$), 49.0 ($^+NH_2$**C**$H_2$Ar), 40.2 (S**C**$H_3$), 24.4 (CH**C**$H_2$), 23.5 (CH(**C**$H_3$) (CH'$_3$)), 22.3 (CH($CH_3$) (**C'**$H_3$), the **C**H($CH_3$)$_2$ peak was overlapped with the DMSO signals; LRMS (M + H)$^+$(ESI$^+$) 365.2 [M + H]$^+$ (calcd for $C_{20}H_{23}F_3N_2OH^+$ 365.2)

**(*S*)-4-Methyl-2-(((3'-(trifluoromethoxy)-[1,1'-biphenyl]-4-yl)methyl)amino)pentanamide methanesulfonate (22k)**

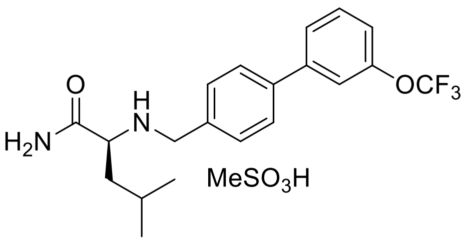


Using method C, to a solution of **15k** (0.15 g, 0.39 mmol), methanesulfonic acid (31.80 μL, 0.49 mmol) in EtOAc (0.78 mL) gave **22k** as a white solid (0.17 g, 93%); $R_f$ = 0.00 ($CH_2Cl_2$/MeOH 10/1); mp 232–233 ˚C; HPLC purity: 8.2 min, 99.3%; $^1$H NMR (300 MHz, DMSO–*d*6) *δ* 9.30 (br s, $^+$N**H**H'), 9.10 (br s, $^+$NH**H'**), 8.08 (br s, C(O)N**H**H'), 7.76–7.84 (m, 4Ar**H** or 3ArH, C(O)NH**H'**), 7.59–7.69 (m, 4ArH or 3ArH, C(O)NH**H'**), 7.41 (d, *J* = 8.2 Hz, 1Ar**H**), 4.03–4.21 (m, $^+NH_2C\mathbf{H_2}Ar$), 3.62–3.76 (m, C(O)C**H**), 2.31 (s, SC**H3**), 1.59–1.75 (m, C**H**, CHC**H2**), 0.82–1.01 (m, 2C**H3**); $^{13}$C NMR (75 MHz, DMSO–*d*6) *δ* 169.9 (**C**(O)), 149.5,

142.2, 139.4, 131.9, 131.5, 131.4, 127.5, 126.3, 120.6 (q, $J_{C–F}$ = 254.8 Hz), 120.5, 119.7 (Ar**C**), 58.4 (C(O)**C**H$^+$NH$_2$), 49.0 ($^+$NH$_2$**C**H$_2$Ar), 40.2 (S**C**H$_3$), 24.4 (CH**C**H$_2$), 23.5 (CH(**C**H$_3$) (C'H$_3$)), 22.3 (CH(CH$_3$) (**C'**H$_3$)), the **C**H(CH$_3$)$_2$ peak was overlapped with the DMSO signals; LRMS (M + H)$^+$(ESI$^+$) 381.2 [M + H]$^+$ (calcd for $C_{20}H_{23}F_3N_2O_2H^+$ 381.2)

**(*S*)-2-(((3'-Methoxy-[1,1'-biphenyl]-4-yl)methyl)amino)-4-methylpentanamide methanesulfonate (22m)**

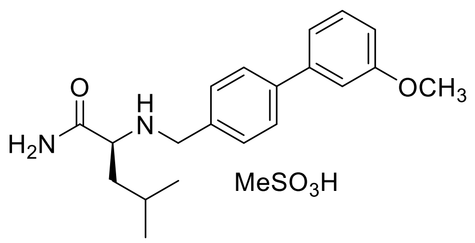


Using method C, to a solution of **15m** (0.15 g, 0.46 mmol), methanesulfonic acid (37.30 µL, 0.57 mmol) in EtOAc (0.91 mL) gave **22m** as a white solid (0.18 g, 93%); R$_f$ = 0.00 ($CH_2Cl_2$/MeOH 10/1); mp 240–242 °C; HPLC purity: 7.1 min, 99.4%; $^1$H NMR (300 MHz, DMSO–*d*$_6$) $\delta$ 9.28 (br s, $^+$N**H**H'), 9.10 (br s, $^+$NH**H'**), 8.11 (br s, C(O)N**H**H'), 7.21–7.82 (m, 2Ar**H**, C(O)NH**H'**), 7.57 (d, *J* = 7.5 Hz, 2Ar**H**), 7.40 (t, *J* = 7.5 Hz, 1Ar**H**), 7.20–7.29 (m, 2Ar**H**), 6.97 (d, *J* = 8.1 Hz, 1Ar**H**), 4.01–4.21 (m, $^+$NH$_2$C**H$_2$**Ar), 3.83 (s, OC**H$_3$**), 3.64–3.79 (m, C(O)C**H**), 2.34 (s, SC**H$_3$**), 1.55–1.77 (m, C**H**, CHC**H$_2$**), 0.80–1.02 (m, CH(C**H$_3$**)**$_2$**); $^{13}$C NMR (75 MHz, DMSO–*d*$_6$) $\delta$ 170.0 (**C**(O)), 160.3(**C**OCH$_3$), 141.3, 141.1, 131.2, 131.1, 130.6, 127.4, 119.5, 113.8, 112.7 (Ar**C**), 58.3 (C(O)**C**H$^+$NH$_2$), 55.6 (O**C**H$_3$), 49.0 ($^+$NH$_2$**C**H$_2$Ar), 40.2 (S**C**H$_3$), 24.4 (CH**C**H$_2$), 23.5 CH(**C**H$_3$) (C'H$_3$)), 22.3 CH(CH$_3$) (**C'**H$_3$)), the **C**H(CH$_3$)$_2$ peak was overlapped with the DMSO signal; LRMS (M + H)$^+$(ESI$^+$) 327.2 [M + H]$^+$ (calcd for $C_{20}H_{26}N_2O_2H^+$ 327.2)

**(2*S*,3*S*)-2-(((3'-Fluoro-[1,1'-biphenyl]-4-yl)methyl)amino)-3-methylpentanamide methanesulfonate (23c)**

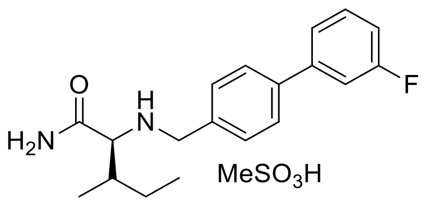


Using method C, to a solution of **16c** (0.15 g, 0.48 mmol), methanesulfonic acid (57.32 µL, 0.60 mmol) in EtOAc (1.19 mL) gave **23c** as a white solid (0.18 g, 94%); $R_f$ = 0.00 ($CH_2Cl_2$/MeOH 10/1); mp 252–255 ˚C; HPLC purity: 7.2 min, >99.9%; $^1$H NMR (300 MHz, DMSO–$d_6$) $\delta$ 9.20 (br s, $^+$N**H**H'), 8.90 (br s, $^+$NH**H'**), 7.87 (br s, C(O)N**H**H'), 7.81 (d, $J$ = 8.25 Hz, 2Ar**H**), 7.77 (br s, C(O)NH**H'**), 7.49–7.61 (m, 5Ar**H**), 7.23–7.28 (m, 1Ar**H**), 4.06 and 4.18 ($AB_q$, $J$ = 13.5 Hz, $^+NH_2C\mathbf{H_2}$Ar), 2.31 (s, $SC\mathbf{H_3}$), 1.80–1.98 (m, C(O)CHC**H**), 1.50–1.58 (m, CHC**H**H'), 1.10–1.20 (m, CHCH**H'**), 0.80–0.93 (m, $CHC\mathbf{H_3}$, $CH_2C\mathbf{H_3}$), the remaining peak was not detected and is believed to overlap with the $H_2O$ signals; $^{13}$C NMR (75 MHz, DMSO–$d_6$) $\delta$ 168.4 (**C**(O)), 163.2 (d, $J_{C–F}$ = 241.8 Hz), 142.2 (d, $J_{C–F}$ = 7.8 Hz), 139.7 (d, $J_{C–F}$ = 2.2 Hz), 131.6, 131.4 (d, $J_{C–F}$ = 8.7 Hz), 131.4, 127.4, 123.2 (d, $J_{C–F}$ = 2.5 Hz), 115.0 (d, $J_{C–F}$ = 20.8 Hz), 113.8 (d, $J_{C–F}$ = 22.0 Hz) (Ar**C**), 63.4 (C(O)**C**$H^+NH_2$), 49.7 ($^+NH_2\mathbf{C}H_2$Ar), 40.2 (S**C**$H_3$), 35.8 (C(O)CH**C**H), 25.6 (CH**C**$H_2$), 14.6 (CH**C**$H_3$), 12.0 (CH2**C**$H_3$); LRMS (M + H)$^+$(ESI$^+$) 315.2 [M + H]$^+$ (calcd for $C_{19}H_{23}FN_2OH^+$ 315.2)

**(*2S*,*3S*)-2-(((3'-Chloro-[1,1'-biphenyl]-4-yl)methyl)amino)-3-methylpentanamide methanesulfonate (23f)**

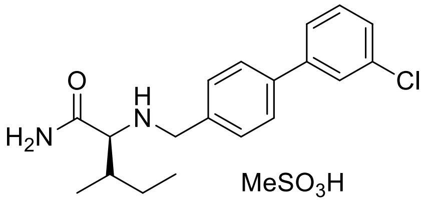


Using method C, to a solution of **16f** (0.20 g, 0.61 mmol), methanesulfonic acid (49.00 µL, 0.76 mmol) in EtOAc (1.20 mL) gave **23f** as a white solid (0.24 g, 95%); $R_f$ = 0.00 ($CH_2Cl_2$/MeOH 10/1); mp 248–250 ˚C; HPLC purity: 7.6 min, 91.7% ; $^1$H NMR (300 MHz, DMSO–$d_6$) $\delta$ 9.20 (br s, $^+$N**H**H'), 8.93 (br s, $^+$NH**H'**), 7.88 (br s, C(O)N**H**H'), 7.77–7.82 (m, 2Ar**H**, C(O)NH**H'**), 7.69 (d, $J$ = 7.6 Hz, 1Ar**H**), 7.59 (d, $J$ = 7.9 Hz, 2Ar**H**), 7.45–7.55 (m, 2Ar**H**), 4.02–4.20 (m, $^+NH_2C\mathbf{H_2}$Ar), 3.51–3.63 (m, C(O)C**H**), 2.31 (s, $SC\mathbf{H_3}$), 1.81–1.98 (m, C(O)CHC**H**), 1.45–1.63 (m, CHC**H**H'), 1.05–1.24 (m, CHCH**H'**), 0.80–0.99 (m, $CHC\mathbf{H_3}$,

$CH_2C\mathbf{H_3}$); [13]C NMR (75 MHz, DMSO–*d*$_6$) $\delta$ 168.4 (**C**(O)), 141.9, 139.6, 134.3, 131.6, 131.4, 131.3, 128.1, 127.4, 126.9, 125.9 (Ar**C**), 63.3 (C(O)**C**H$^+$NH$_2$), 49.6 ($^+$NH$_2$**C**H$_2$Ar), 35.8 (C(O)CH**C**H), 25.6 (CH**C**H$_2$), 14.6 (CH**C**H$_3$), 12.0 (CH$_2$**C**H$_3$), the S**C**H$_3$ peak was overlapped with the DMSO signals; LRMS (M + H)$^+$(ESI$^+$) 331.2 [M + H]$^+$ (calcd for $C_{19}H_{23}ClN_2OH^+$ 331.2)

**(2*S*,3*S*)-2-(((4'-Chloro-[1,1'-biphenyl]-4-yl)methyl)amino)-3-methylpentanamide methanesulfonate (23g)**

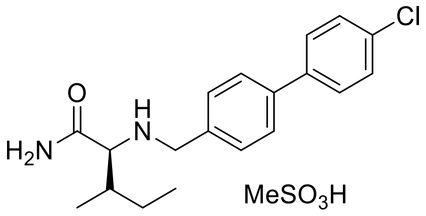


Using method C, to a solution of **16g** (0.14 g, 0.41 mmol), methanesulfonic acid (33.10 µL, 0.51 mmol) in EtOAc (0.82 mL) gave **23g** as a white solid (0.15g, 88; R$_f$ = 0.00 ($CH_2Cl_2$/MeOH 10/1); mp 260–262 ˚C; HPLC purity: 7.7 min, 99.3%; [1]H NMR (300 MHz, DMSO–*d*$_6$) $\delta$ 9.19 (br s, $^+$N**H**H'), 8.92 (br s, $^+$NH**H'**), 7.88 (br s, C(O)N**H**H'), 7.77–7.82 (m, 5Ar**H**, or 4Ar**H**, C(O)NH**H'**), 7.53–7.60 (m, 4Ar**H** or 3Ar**H**, C(O)NH**H'**), 4.06 and 4.17 (AB$_q$, $J$ = 12.4 Hz, $^+$NH$_2$C**H$_2$**Ar), 3.51–3.64 (m, C(O)C**H**), 2.32 (s, SC**H$_3$**), 1.82–1.97 (m, C(O)CHC**H**), 1.46–1.64 (m, CHC**H**H'), 1.02–1.27 (m, CHCH**H'**), 0.81–0.99 (m, CHC**H$_3$**, CH$_2$C**H$_3$**); [13]C NMR (75 MHz, DMSO–*d*$_6$) $\delta$ 168.4 (**C**(O)), 139.8, 138.6, 133.1, 131.6, 131.1, 129.4, 128.9, 127.2 (Ar**C**), 63.4 (C(O)**C**H$^+$NH$_2$), 49.7 ($^+$NH$_2$**C**H$_2$Ar), 35.8 (C(O)CH**C**H), 25.6 (CH**C**H$_2$), 14.6 (CH**C**H$_3$), 12.0 (CH$_2$**C**H$_3$), the S**C**H$_3$ peak was overlapped with the DMSO signals; LRMS (M + H)$^+$(ESI$^+$) 331.2 [M + H]$^+$ (calcd for $C_{19}H_{23}ClN_2OH^+$ 331.2)

**(2*S*,3*S*)-3-Methyl-2-(((3'-(trifluoromethyl)-[1,1'-biphenyl]-4-yl)methyl)amino)pentanamide (23i)**

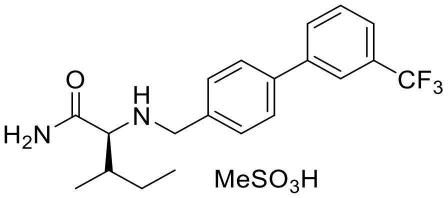


Using method C, to a solution of **16i** (0.10 g, 0.27 mmol), methanesulfonic acid (22.26 µL, 0.34 mmol) in EtOAc (0.55 mL) gave **23i** as a white solid (0.12 g, 97%); $R_f$ = 0.00 ($CH_2Cl_2$/MeOH 10/1); mp 263–265 ˚C; HPLC purity: 8.0 min, 98.8%; $^1$H NMR (300 MHz, DMSO–$d_6$) $\delta$ 9.20 (br s, $^+$N**H**H'), 8.93 (br s, $^+$NH**H'**), 7.98–8.08 (m, 1Ar**H**, C(O)N**H**H'), 7.82–7.92 (m, 2Ar**H**, C(O)NH**H'**), 7.69–7.80 (m, 3ArH), 7.63 (d, $J$ = 7.8 Hz, 2Ar**H**) 4.08 and 4.19 ($AB_q$, $J$ = 11.9 Hz, $^+NH_2C$**$H_2$**Ar), 3.50–3.59 (m, C(O)C**H**), 2.31 (s, SC**H3**), 1.80–1.98 (m, C(O)CHC**H**), 1.44–1.68 (m, CHC**H**H'), 1.09–1.26 (m, CHCH**H'**), 0.80–0.99 (m, CHC**H3**, $CH_2$C**H3**); $^{13}$C NMR (75 MHz, DMSO–$d_6$) $\delta$ 168.4 (**C**(O)), 140.8, 139.5, 131.7, 131.6, 131.3, 130.7, 130.3 (q, $J_{C–F}$ = 31.4 Hz), 127.6, 124.8, 124.6 (q, $J_{C–F}$ = 270.8 Hz), 123.5 (q, $J_{C–F}$ = 3.7 Hz) (Ar**C**), 63.4 (C(O)**C**H$^+NH_2$), 49.6 ($^+NH_2$**C**$H_2$Ar), 35.8 (C(O)CH**C**H), 27.6 (CH**C**$H_2$), 14.6 (CH**C**$H_3$), 12.0 (CH2**C**$H_3$), the S**C**H3 peak was overlapped with the DMSO signals; LRMS $(M + H)^+(ESI^+)$ 365.2 $[M + H]^+$ (calcd for $C_{20}H_{23}F_3N_2OH^+$ 365.2)
